# Machine Learning-Assisted Evolution of Broadly Functional Enzyme Libraries

**DOI:** 10.64898/2026.07.23.740427

**Authors:** Ravi G. Lal, Jason Yang, Ziyan Zhang, Frances H. Arnold

## Abstract

Biocatalysis offers sustainable solutions to pressing challenges in chemical synthesis by exploiting the remarkable efficiency and selectivity of enzymes. Importantly, enzymes are able to accommodate non-native substrates and mediate transformations outside of their natural repertoire. Enzymes can be engineered for diverse applications by harnessing these ‘promiscuous’ activities and optimizing them using directed evolution (DE). The success of a DE campaign, however, depends on the availability of a protein starting point that displays detectable levels of the desired function. To find a starting point, researchers often screen libraries of protein variants for novel activities, typically with low rates of success. Here, instead, we diversified the active site of a desirable ‘parent’ protein and applied machine learning to generate informed, promiscuous libraries of protein variants. Specifically, we tested 26 different carbene and nitrene transfer reactions and used active learning-assisted directed evolution (ALDE) to generate optimized protoglobin variants with high activity across multiple reactions. We observed improvements in activity and selectivity for every reaction performed by the parent enzyme in at least one member of the ALDE-predicted libraries. Moreover, variants from these libraries can catalyze 5 out of 10 reactions not catalyzed by the parent protoglobin. These results indicate that supervised machine learning can help guide the construction of high-value enzyme libraries with expanded catalytic scope.

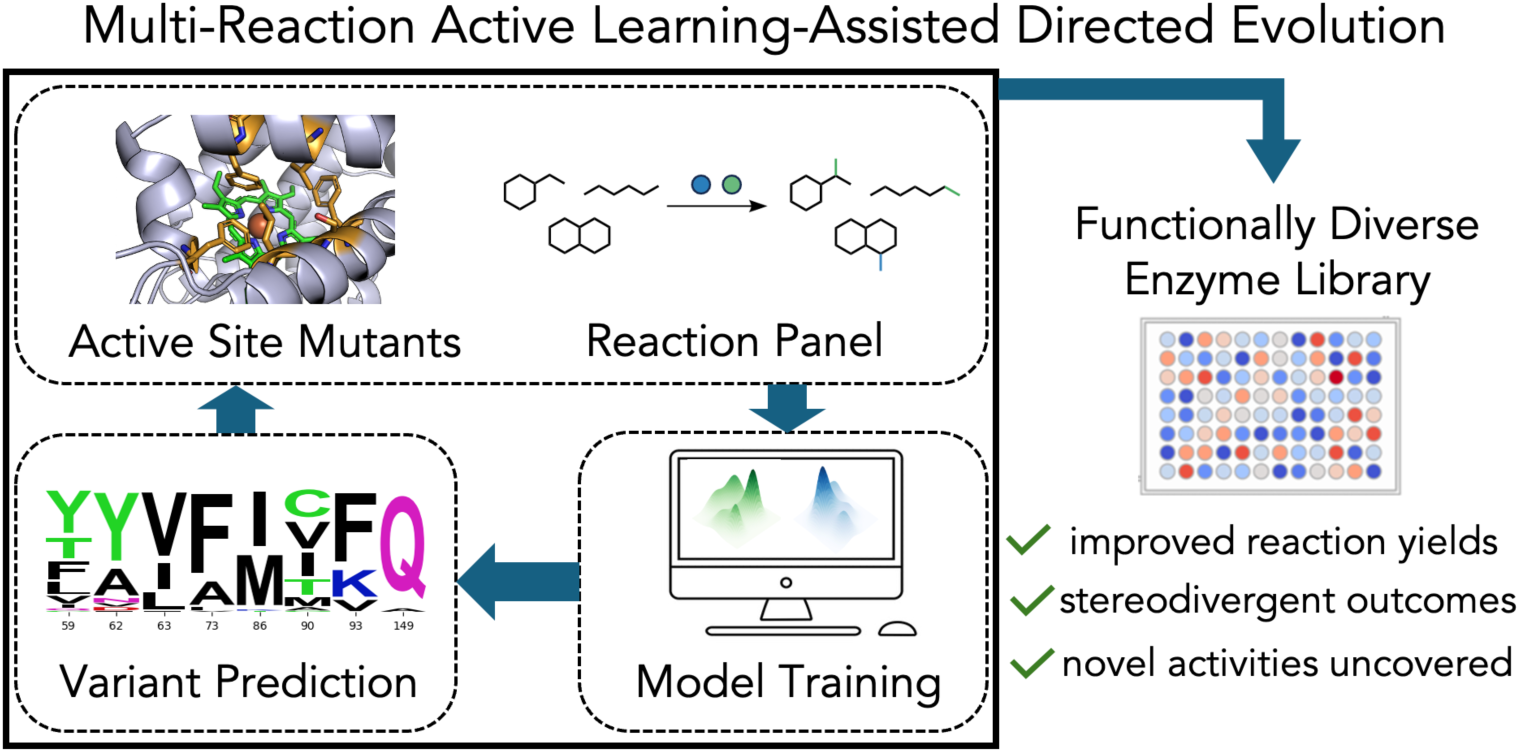

## INTRODUCTION

Biocatalysis harnesses enzymes, Nature’s catalysts, to perform chemical transformations with remarkable efficiency and selectivity, offering sustainable solutions to challenges in medicine, agriculture, and industry.^1^ A key feature underpinning this utility is the capacity of enzymes to accommodate non-native substrates (substrate promiscuity) or to catalyze reactions via non-native mechanisms (catalytic promiscuity).^2,3^ Researchers have leveraged enzyme promiscuity to develop valuable biocatalytic transformations. However, promiscuous activities are rarely optimal and thus require improvement using directed evolution (DE), in which sequential rounds of diversification and screening are used to enhance the desired activity. Such approaches have delivered biocatalysts with broadened substrate scopes^4,5^ and even entirely new-to-nature reaction modes^6–10^.

Directed evolution of a novel enzymatic activity starts with a protein that displays at least trace levels of the desired function such that a round of mutation and screening can reliably generate improvements. While significant advances have been made in optimization strategies and laboratory practices to accelerate and improve DE outcomes,^11^ the discovery of starting points displaying a target function remains a bottleneck, as it is challenging to predict promiscuous functions from sequence alone.^12,13^ This challenge is especially pronounced for new-to-nature biocatalytic transformations, where protein fitness data, which could be used to infer promiscuous functions, are scarce and rarely describe activity for more than one substrate.^14^ Given these challenges, the identification of new activities commonly relies on experimental testing of diverse but somewhat random collections of proteins (**Figure 1A**); in the best case these may be informed by biochemical intuition and prior protein engineering experience.^15^ Nonetheless, these mostly uninformed “shots in the dark” often result in low success rates for finding the desired functions. Recently, there have been efforts to design and build *informed* libraries of enzymes which display broad promiscuity, with the goal of improving the probability of discovering novel enzymatic functions upon screening.^14,16–21^ One strategy has been to generate libraries of enzyme variants which contain diverse active-site mutations but still retain their functionality. The driving hypothesis is that diverse active sites within an enzyme scaffold already capable of a desired reaction mode could display a sort of *ensemble promiscuity* by offering variants which span a greater breadth of the protein fitness landscapes.^22^ The efficient development of such libraries, however, is nontrivial as simultaneous incorporation of multiple mutations can destabilize the protein.^23,24^ Furthermore, epistasis, the phenomenon whereby the effects of a mutation depend on the context in which it is introduced, makes it challenging to predict how combinations of several mutations will behave.^25^ The computational package FuncLib developed by Fleishman and coworkers has been one effective answer to these challenges.^26^ Broadly, this tool predicts functional enzyme variants containing mutations at user-defined active-site residues by (1) assessing the stability of all possible variants using physics-based models and (2) designing combinations of interacting residues that will be compatible with function based on these same models and evolutionary information inferred from existing sequence diversity. Although effective,^27,28^ this method performs zero-shot design without the ability to learn from real-world assay labels or specific substrate/reaction promiscuity goals.

**Figure 1.**
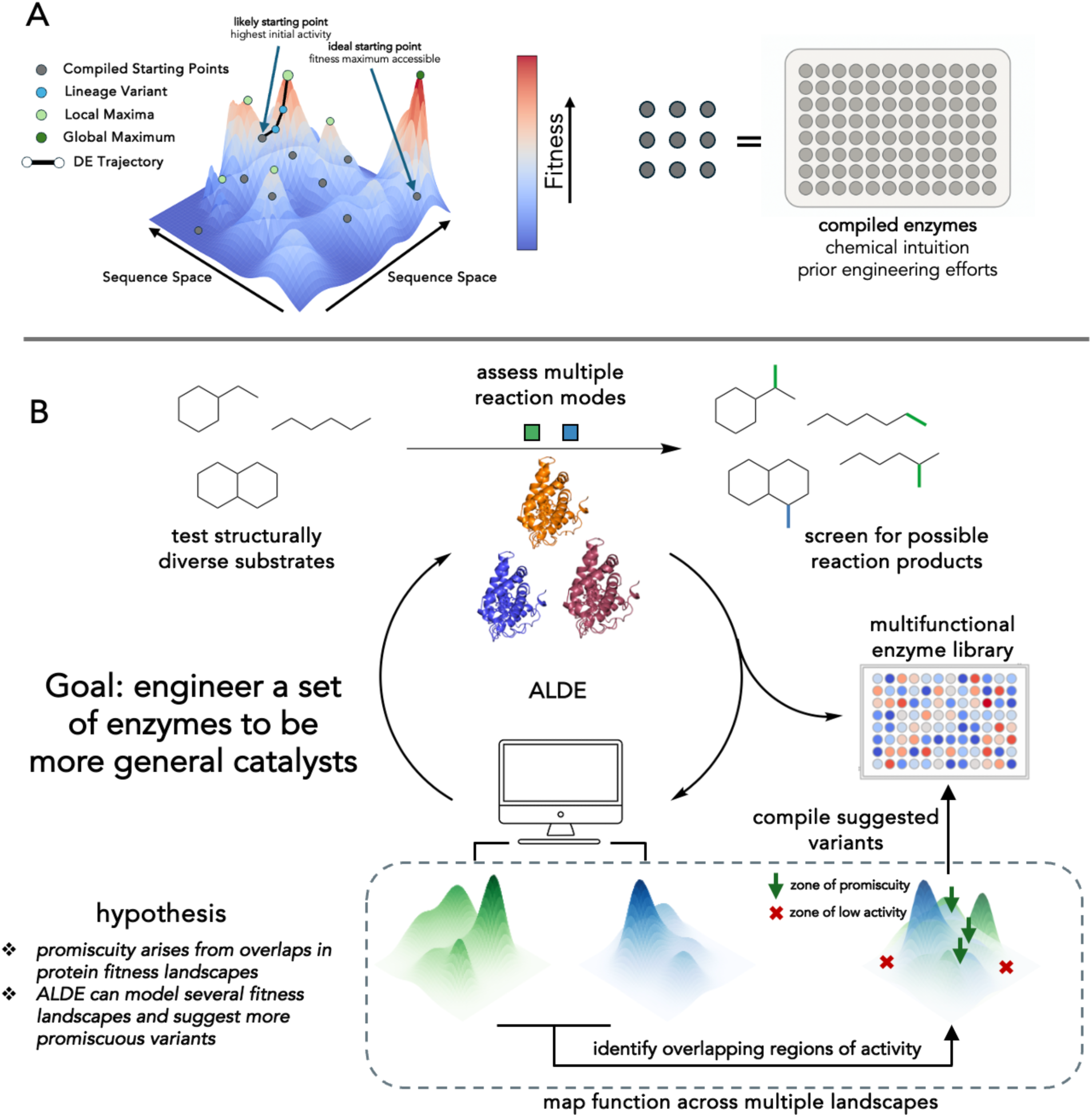
Promiscuity in directed evolution. **(A)** Directed evolution is a greedy uphill walk in a protein sequence- function landscape. Starting points for DE are selected from compiled libraries of enzymes. The starting point influences the outcomes of DE. **(B)** Proposed framework for using ALDE to optimize a family of enzymes for multiple activities. An ALDE model is trained using data from active-site mutants of a parent enzyme assayed for multiple reactions. The model then suggests variants which it predicts have improved activity across reactions seen in training. The predicted enzymes are compiled and assessed for their ability to catalyze reactions both in and out of the training set.

Emerging machine learning (ML) methods have the potential to enable the generation or optimization of more-promiscuous enzymes.^29,30^ We focus here on machine learning-assisted directed evolution (MLDE) methods, which are being used to predict combinations of beneficial mutations given sequence-fitness data on protein variants with multiple mutations, including at sites which leverage positive epistatic interactions.^31–36^ Recently, we described active learning-assisted directed evolution (ALDE), a framework in which an ML model is updated in an iterative manner after collecting new data, and applied it to optimize yields of non-native enzymatic carbene transfer reactions catalyzed by protoglobins, a class of microbial globin.^37,38^ While previous MLDE efforts have been aimed toward navigating sequence-function landscapes for a single catalytic task, we reasoned that sequence optimization across several reaction types could enable the discovery of enzyme variants which exist at overlaps of these landscapes, which we propose are ‘zones of promiscuity’ (**Figure 1B**). Our hypothesis in this work was that such promiscuous variants would have a higher probability of displaying activity for related reactions and substrates in future screens.

Here, we extend ALDE to multi-objective optimization and describe ‘<u>m</u>ulti-reaction <u>A</u>ctive <u>L</u>earning-assisted <u>D</u>irected <u>E</u>volution’ (mr-ALDE), in which a machine learning model is trained on functional data spanning multiple reactions assayed on a set of variants harboring active-site mutations. This model is then prompted to predict multi-mutation variants that will be more active toward both carbene and nitrene reactions. We reasoned that this framework could furnish enzyme collections well suited for initial screening campaigns seeking activities related to but different from the training reactions (**Figure 1B**). To test this strategy, we used mr-ALDE to develop protoglobin variants broadly functional toward new-to-nature carbene and nitrene transfer chemistries. Using an ML-model trained on function data for over 500 multi-mutation variants originating from a single ‘parent’ protoglobin, we predicted and tested 127 protoglobins with mutations in the active site. These variants, when considered as an ensemble, were capable of performing nearly every tested reaction in higher yield than the starting sequence and could also access reactions which the initial sequence could not deliver with detectable yield. Thus, supervised machine learning can be used to suggest protoglobin variants with active sites that are optimized for promiscuity.

## STUDY DESIGN

The goal of this study was to develop an ALDE-based workflow for constructing enzyme libraries exhibiting broad activity across new-to-nature enzymatic carbene and nitrene transfer reactions.^39^ The approach uses sequence–function data from multi-mutation variants to train predictive models that guide the selection of broadly functional enzymes. Our implementation began with the selection of a parent enzyme and proceeded through three stages: 1) training data collection, 2) round 1 prediction assessment, and 3) round 2 prediction assessment, each separated by *in silico* model training and proposal of new variants to test. We set out to apply ALDE-based enzyme diversification to simultaneously access variants with improved activity toward an expanded scope of both carbene and nitrene transfer reaction modes. This engineering goal would enable us to investigate mr-ALDE’s ability to deliver enzymes with both expanded substrate range and mechanistic scope.

To initiate the study, we first determined a ‘parent’ enzyme which could (1) access a broad scope of chemical reactivities and (2) stand up to a mutational load of up to eight amino acid substitutions. Protoglobins originating from Archaea have been engineered for a variety of carbene and nitrene transfer activities in previous studies^40–45^ and are highly thermostable.^24,38,46^ Furthermore, we have shown that active-site residues in protoglobins exhibit epistasis toward non-native activities and that ALDE can effectively predict synergistic mutation combinations for specific catalytic tasks.^37^ Because protoglobins have no known native catalytic function, we believed that zero-shot mutation effect prediction methods for designing variants of these proteins would not be effective for accessing catalytically functional enzymes, particularly for activities such as carbene and nitrene transfer, for which there is no fitness information encoded in protein sequences produced by natural evolution.

We focused on various new-to-nature reactions which had previously been demonstrated in hemoproteins.^40,42–45,47,48^ This set of <u>I</u>nitial <u>T</u>raining <u>R</u>eactions (ITRs) comprised four carbene-mediated cyclopropanation reactions and four nitrene-based reactions spanning C–H bond functionalization, aromatic desymmetrization, and boronic acid amination mechanisms (**Scheme 1**). These reactions were selected to challenge the active site along two axes: the cyclopropanation set tests whether a common carbene-transfer mechanism can be applied to alkene substrates of varying steric and electronic character, while the interrogation of both carbene and nitrene transfer reactions examines the enzyme’s catalytic promiscuity toward distinct mechanistic manifolds.

**Scheme 1.**
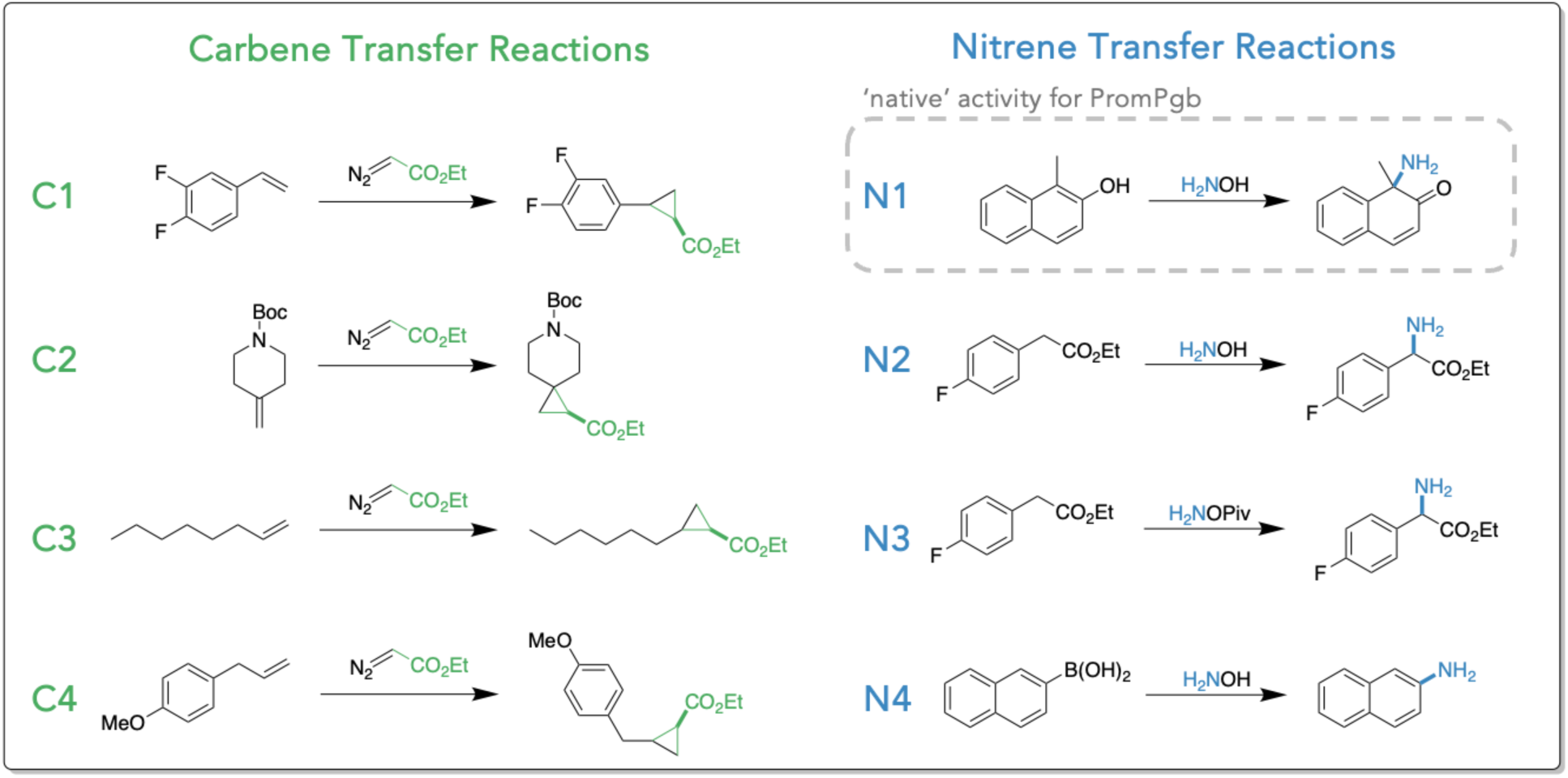
Initial training reactions (ITRs) used to identify a promiscuous protoglobin and first train the ALDE model.

Through screening the ITRs with the precompiled panel of 84 wild-type and engineered protoglobin analogs engineered previously for reactions related to the ITRs, we identified a variant of *Aeropyrum pernix*protoglobin (*Ape*Pgb), evolved previously for dearomatization chemistry (**N1 in Scheme 1**),^45^ that demonstrated measurable activity for all ITRs except **N3** (C–H insertion with a hydroxylamine-derived nitrenoid), with product yields varying across the different transformations. This variant, designated PromPgb (<u>Prom</u>iscuous <u>P</u>roto<u>g</u>lo<u>b</u>in), was selected as the parent enzyme for application of mr-ALDE.

Before ALDE is initiated, the design space—defined by a selected set of *k* residues—is specified, corresponding to 20*^k^* potential sequence variants. For this study, we identified eight positions, which are facing toward the distal side of the heme cofactor (where substrates bind): L59, W62, F63, F73, L86, S90, F93, and S149 (denoted LWFFLSFS, **Figure 2A**). Single mutations at some of these residues were shown to impact non-native function in previous engineering efforts;^43,44^ this prior knowledge is helpful but not necessarily essential for choosing active site residues, as they can also be selected based on MSAs and/or structural criteria. The combinatorial space of all variants containing mutations at these positions is > 2.5 x 10^10^. Although a library of variants containing mutations at all design sites would better capture the sequence–function landscapes we sought to model, random sampling of this space was expected to produce mostly non-functional enzymes. Thus, for the training library we tested double- and triple-site variants containing mutations at subsets of these positions (**Figure 2B**) in order to reduce the overall mutational load of the initial training set. Because the majority of functional epistatic outcomes in proteins can be attributed to pairwise interactions,^49,50^ we reasoned that randomly selected protoglobin variants with 2–3 mutations would be less likely to be inactive toward the ITRs while still reporting on key relationships between active-site residues. Twelve double-site libraries and six triple-site libraries were designed based on structural proximity (assessed on a structure prepared using AlphaFold) and ease of construction (**Table S8** of **Supporting Information**).

**Figure 2.**
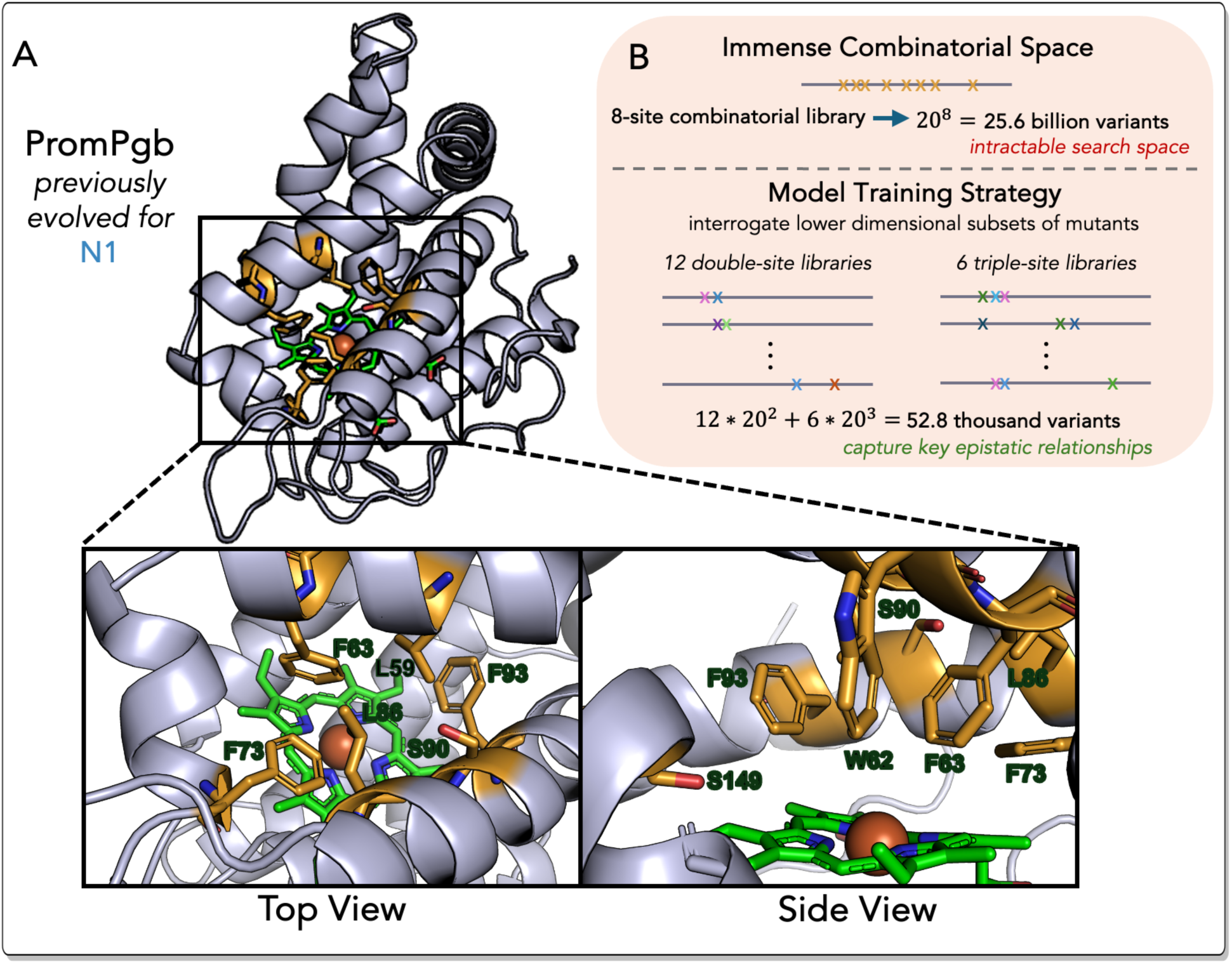
**(A)** The active site of PromPgb has several interacting residues above the distal face of the heme cofactor. Eight residues were selected for the design space in this study: L59, W62, F63, F73, L86, S90, F93, and S149. **(B)** To reduce the deleterious effects of increasing numbers of mutations, the initial round of sequences was built as a combined library of selected double- and triple-site combinatorial libraries.

These libraries covered 22 of the 28 possible combinations of the selected eight residues. Libraries were constructed using multi-site saturation mutagenesis techniques (**Cloning Protocols and Results** of **Supporting Information**).^51^ Twenty-two unique variants from each double-site library and 44 from each triple-site library, characterized by nanopore-based multiplexed sequencing (LevSeq),^52^ were arrayed, affording 528 sequences (1% of the possible search space) for initial reaction screening (**Table S9** of **Supporting Information**).

This collection of 528 PromPgb variants was evaluated against the eight ITRs, yielding a comprehensive dataset of >4,000 sequence-function pairs (Training Data Collection, **Figure 3B**). The parent-normalized fitness data for the ITRs were grouped into two objectives, carbene and nitrene reactions, for ALDE model training. Essentially, for each training library variant, the average change in activity from PromPgb for the carbene transfer ITRs (**C1**-**C4**) and the nitrene transfer ITRs (**N1**-**N4**) were separately computed as fitness objectives (**Machine Learning Details** of **Supporting Information**). We then trained an ensemble of multi-task supervised ML models to predict both fitness objectives from encodings of sequence. Afterward, the expected hypervolume improvement acquisition function was applied to the trained model to rank candidate sequences in the design space from most to least likely to improve **both** fitness objectives, balancing *exploration* of new areas of sequence space (mutation combinations with high uncertainty) with *exploitation* of variants that are predicted to have high fitness. We focused on the mean improvement to carbene and nitrene activity as the two objectives to optimize for two reasons: (1) to increase regularization during training of the multi-task ensemble of surrogate models and (2) to reduce the computational cost of calculating the expected hypervolume improvement acquisition function.^53^ The proposed variants (63 distinct sequences; Round 1 Predictions) were tested in the wet lab with 14 new reactions alongside the ITRs. The function data for these 22 reactions (13 carbene transfer reactions and nine nitrene transfer reactions) were then used to update the learning model for a second round of predictions to obtain further improved variants (**Figure 3A**). This final round of variants (64 distinct sequences; Round 2 Predictions) was then interrogated against a panel of the previous 22 reactions and four additional reactions (**Scheme 2A**). Across these two rounds of model-guided experimentation, we evaluated the catalytic performance of predicted protoglobin variants and used these data to assess mr-ALDE’s ability to generate a set of functional enzymes which can access a broader range of activities than the parent.

**Figure 3.**
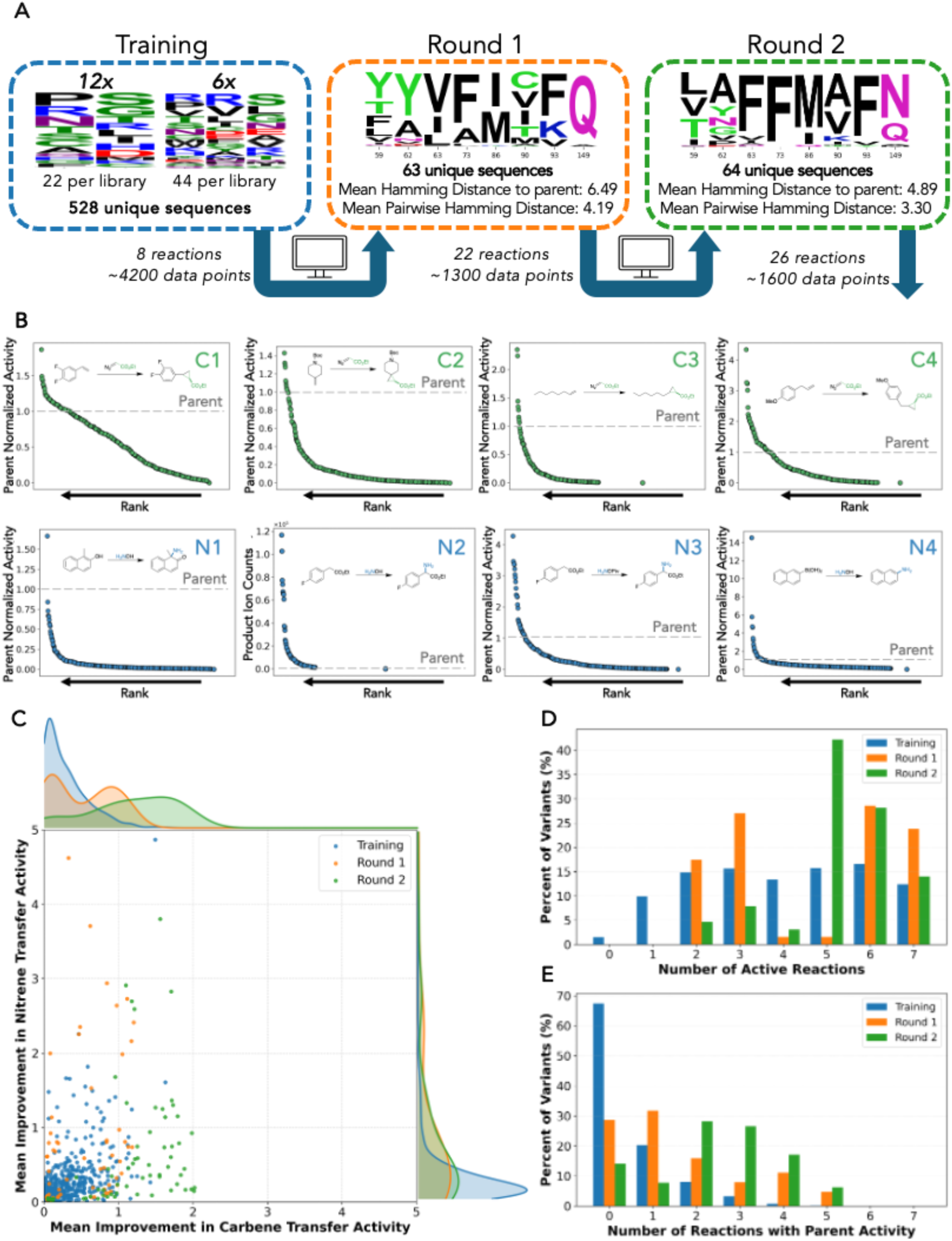
**(A)** After training data were collected, a panel of PromPgb variants harboring mutations at the eight positions were proposed through the ALDE algorithm. These multi-mutants (Round 1) were screened on 22 carbene and nitrene transfer reactions for further model training. After a second round (Round 2) of mutants was predicted, a total of 26 reactions was assessed. **(B)** Initial round of activity data collected for eight chemical reactions with random 2–3 site mutants of PromPgb. Reactions C1–C4 were assessed by gas chromatography with detection with a flame ionization detector (GC-FID). Reactions N1–N4 were assessed by liquid chromatography with mass- spectrometric detection (LC-MS). For reactions C1, C3, and C4 activities are taken as the sum of the yields of the two possible cyclopropane diastereomers. **(C)** The predicted variants display yields that generally represent higher values of the objective functions used to train the model. The evaluated objective functions shown here are computed only with data for reactions seen only by the original training set. Improvement is normalized across all reactions, with 1 referring to parent activity. **(D)** Counts of the total number of ITRs which each training-library or model-predicted design was capable of facilitating with detectable yield. Reaction **N2** is not considered in these counts as PromPgb does not perform this reaction. **(E)** Counts of the number of ITRs which each training-library or model-predicted design catalyzed with greater yield than the parent sequence, PromPgb.

## RESULTS

### Model Training is Informed by an Extensive Functional Landscape

Initial model training was based on experimental data obtained by screening a library of 264 double-site and 264 triple-site variants of PromPgb against the eight ITRs. Among the 8 x 19 = 152 possible single amino-acid substitutions at the eight design-space positions, 151 were present in the constructed library at least once (**Figure S21** of **Supporting Information**). Screening revealed that mutations within the training library substantially altered catalytic performance across all eight ITRs (**Figure 3B**). Additionally, we found that the identities of residues in the active site could drastically change the stereochemical outcomes of the cyclopropanation reactions **C1**, **C3**, and **C4** (**Figures S24**, **S28**, and **S31** of **Supporting Information**). Excitingly, nearly 10% of variants in the training library were capable of catalyzing C–H amination using hydroxylamine as a nitrene precursor (reaction **N2**), the only ITR not catalyzed by the parent protoglobin PromPgb. While members of the training library already displayed broadened specificity, applying ALDE should allow us to distill this information into a smaller, more diverse set of enzyme designs requiring fewer resources to screen. Overall, we were encouraged to see that combinations of mutations at these positions could positively impact activities toward all eight of the ITRs.

Before proceeding with model training, we examined how changes in activity correlated across reactions for each tested variant. A Pearson correlation analysis (**Figure S68** of **Supporting Information**) revealed that the seven ITRs accessible to PromPgb were generally, albeit quite weakly, correlated (average r = 0.28). This suggests that mutations influencing enzyme performance often affect multiple reactions, but some underlying structural and mechanistic determinants can remain distinct. These correlations likely also capture global effects of mutations on enzyme expression and stability, suggesting that part of the shared activity behavior arises from changes in overall protein abundance or folding. Such information remains valuable for model training, as these are desirable traits that we aim to retain and propagate in model-suggested variants. Reaction **N2**—the only ITR that PromPgb could not catalyze with detectable yield— displayed the weakest correlations overall, including below-average correlations with **N1** and **N3**, which share its nitrene precursor and organic co-substrate, respectively. These trends highlight the mechanistic complexity of introducing new chemistries into enzymes and emphasize the importance of sampling diverse active-site configurations to uncover novel functions.

### ALDE Generates Diverse Libraries of Functional Protoglobins

We performed two rounds of mr-ALDE, which included model training, experimental validation, and model updating, to explore the defined design space (**Figure 3A**). In each round, an oligo pool-based cloning strategy was applied to achieve exact model-predicted multi-mutants of PromPgb. Details regarding DNA sequence design are described in the supplementary materials (**Cloning Protocols and Results** of **Supporting Information**). In the first round, 63 of the 96 top-ranked variants were successfully cloned and characterized by LevSeq sequencing.^52^ These sequences had a mean Hamming distance of 6.49 amino acids from PromPgb and a mean pairwise distance of 4.19 amino acids. In the second round, informed by updated activity data from Round 1, 64 of the 96 top-ranked variants were obtained and characterized, with a mean Hamming distance of 4.89 amino acids from PromPgb and a mean pairwise distance of 3.30 amino acids. While sequence diversity varied between the two rounds of prediction, the mean number of mutations per variant in both libraries remained higher than the maximum of three mutations present in the training set.

To understand the performance of the modeling strategy, we first evaluated how well ALDE- predicted protoglobin variants retained their function by comparing their activities on ITRs with those of the initial training variants. Here we only consider the seven ITRs for which PromPgb demonstrates detectable yield (reactions **C1–C4**, **N1**, **N3**, and **N4**). Examination of these seven reactions showed that, in general, predicted variants demonstrated improved average activity toward carbene and nitrene chemistries when compared to the parent PromPgb and members of the training library (**Figure 3C**). Across both rounds, ALDE-predicted variants more frequently retained basal activity (≥5% of parent yield) toward the parent-accessible ITRs than did the random multi-mutants in the training library, with median accessibility of six and five reactions in the first and second rounds, respectively, versus four in the training library (**Figure 3D**). Even more excitingly, members of the designed libraries were far more likely to demonstrate activities higher than parent for a greater number of reactions than training variants (**Figure 3E**). While only 33% of training library members were capable of performing any ITR with greater yield than PromPgb, nearly 70% of protoglobin variants in both predicted libraries were capable of catalyzing at least one ITR with greater yield than PromPgb. This enrichment of protoglobin variants which have improved retention of function and even heightened activities demonstrates that MLDE is effective at synthesizing information from a multi-reaction dataset to predict diverse combinations of active-site mutations.

PromPgb was previously engineered for reaction **N1,** and this activity can be considered the “native” function of the protein. We found no predicted sequences which could catalyze this dearomatization transformation as well as PromPgb. The best mutant for reaction **N1** (VAVFIAFN; **Table 1**, Entry 6) could only catalyze this reaction with 11% yield (one fifth the yield of PromPgb), indicating a tradeoff between the improvement of a promiscuous activity and the “native” function.^54,55^ In the training data for reaction **N1**, we only observe one multi-variant that is capable of catalyzing this reaction with higher yield than PromPgb. As a result, the model has not been trained with sufficient data to predict mutations which will improve this activity. Nevertheless, the identification of variants retaining activity toward **N1** while gaining activity in other reactions underscores the potential to balance tradeoffs through strategic library design to achieve a compilation of enzymes with generally heightened catalytic competency.

**Table 1.**
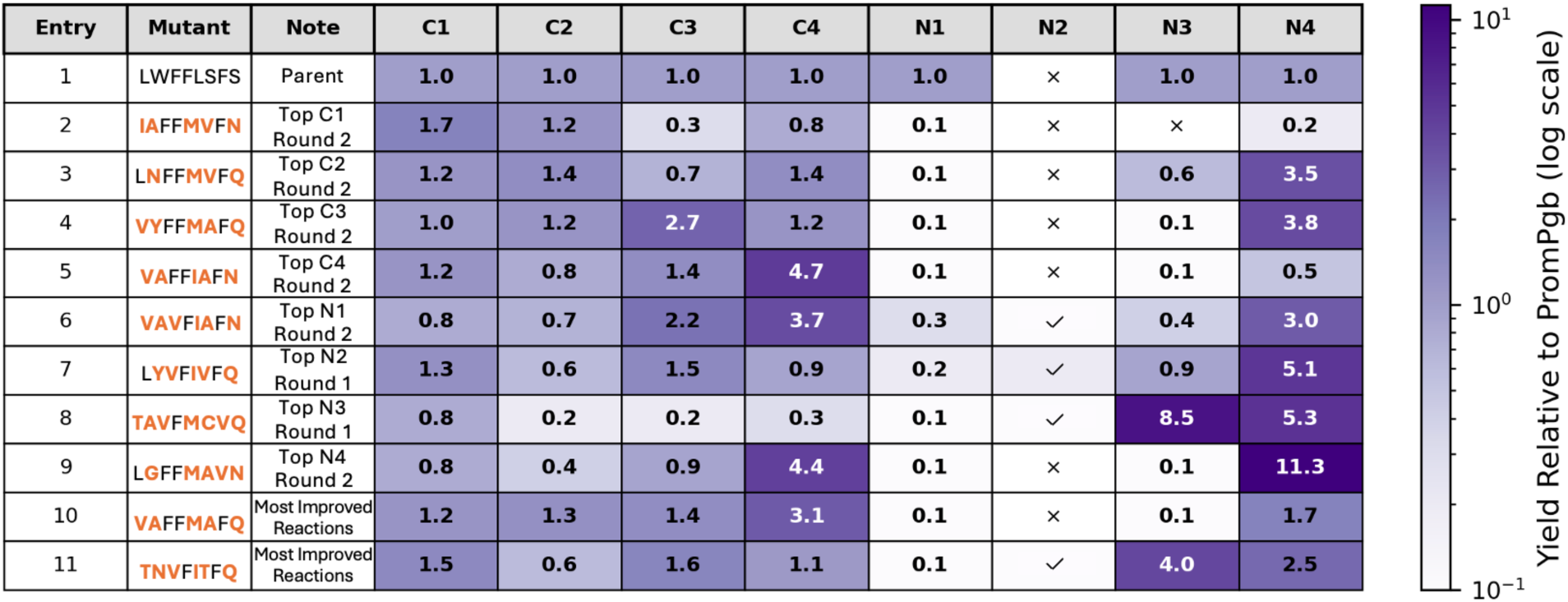
Heatmap of fold-improvements for ITRs for select variants from predicted protoglobin libraries. Cell coloring is shown as the log-fold change in yield relative to PromPgb. Absolute fold-changes in reaction yield are given in each cell. Rows 2–9 represent the top-yielding variant for reactions **C1**–**C4** and **N1**–**N4,** respectively (top reaction boxed). Rows 10 and 11 show variants show the variants for which the greatest number of activities were improved relative to PromPgb by the greatest degree from each round of predictions. For variants displaying activity for reaction **N2**, a check mark indicates the formation of the C–H amination product. Shading in cells for reaction **N2** is calculated based on a variant’s yield for this reaction normalized to the yield of PromPgb for reaction **N3**, which shares the same product.

Consideration of the top-performing ALDE design for each of the eight ITRs reveals varying degrees of specialization. The variants which demonstrated the highest activities for carbene transfer ITRs (reactions **C1**–**C4**) tended to maintain their activities for other carbene transfer reactions, while taking major losses in activity for all nitrene transfer ITRs (**N1**–**N4**) besides **N4 (Table 1**, Entries 2–5). In contrast, multi-mutants which displayed the highest yields toward nitrene transfer chemistries tended to maintain carbene transfer activities to some degree **(Table 1**, Rows 6-9). Notably, the two designs which catalyzed the greatest number of ITRs with higher yields than PromPgb (VAFFMAFQ and TNVFITFQ; **Table 1**, Entries 10 and 11) were not the highest performers for any single ITR.

Together, these findings underscore the power of ALDE-based library design to uncover both specialized and broadly functional catalysts within a single protein scaffold. By sampling mutational combinations that enhance reactivity across mechanistically distinct reactions, this approach identifies variants that not only expand the accessible catalytic repertoire but also reveal generalist sequences from which new activities can be evolved. Achieving comparable functional breadth through conventional directed evolution would demand several separate campaigns, each targeting a distinct reaction, whereas this ALDE design cycle furnished 127 catalytically active and mechanistically diverse variants upon testing only 655 sequences.

### Model-Predicted Variants Demonstrate Broadened Substrate Range as an Ensemble

Having established that the mr-ALDE framework generates protoglobin variants with enhanced activity toward catalytic functions accessible to PromPgb, we next examined how this functional diversity translates to substrate range. To this end, we evaluated the ensemble of model-predicted variants across an expanded substrate panel representing electronically and sterically diverse carbene and nitrene acceptors (**Scheme 2A**). This set of reactions comprised 18 transformations not seen in initial model training. Broadly, the reactions interrogated can be classified into two primary mechanistic categories: additions to π-systems and insertions into X–H bonds. Four additional transformations (reactions **N1**, **N4**, **C10**, and **C11**) likely proceed through distinct mechanisms that fall outside of these two classes. The collected data for all transformations are provided in the Supporting Information and are available in full on GitHub.

For every transformation that PromPgb could catalyze with detectable product formation, we identified at least one model-predicted variant that was more active than the parent. Notably, these improvements extended even to reactions that are mechanistically distinct from those represented among the ITRs, indicating that the sequence features enriched during mr-ALDE can generalize to new reactivity modes. For example, variant FYIFMMFQ catalyzes the intramolecular C(*sp³*)– H amination of alkyl azide **1** to afford pyrrolidine **2** in 3% yield (**Reaction N6** in **Scheme 2B**), whereas PromPgb furnishes only trace product under identical conditions. Strikingly, this variant emerged in the first round of ALDE, trained solely on ITR data, despite the fact that reaction **N6** requires both azide activation and an intramolecular cyclization—mechanistic steps not present in the ITR panel. We also observe cases that highlight the value of the iterative model-updating component of ALDE. For reactions **C4** and **C8**, second-round predictions yielded improvements in a larger fraction of variants compared to the first round (**Figures S30** and **S39** of **Supporting Information**). In the case of reaction **C13** (intermolecular carbene C–H insertion of pyrrolidine **3**), the second round yielded variant VVFFMAFN, which delivers product **4** in 6% yield, representing a six-fold increase relative to PromPgb (**Reaction C13** in **Scheme 2B**). To achieve the observed levels of activity for reactions **N6** and **C13** through conventional directed evolution of PromPgb would likely require multiple rounds of mutagenesis and screening. Thus, although the absolute yields for these reactions remain modest, the top-performing variants identified here constitute improved starting points for subsequent directed evolution efforts.

None of the data used in either round of ALDE model training encoded the stereochemical outcomes of any training reactions. All reactions with multiple diastereomeric products were recorded as the sum of both stereoisomers. Nevertheless, by training on substrates with varied steric and electronic profiles, ALDE guided the exploration of active-site environments that differ in configuration, ultimately producing variants that favor different stereochemical trajectories of the same reaction. For nearly all of the cyclopropanation reactions tested in this work, the complete ensemble of model-predicted variants is capable of delivering stereodivergent outcomes toward distinct diastereomeric products (**Scheme 2C**). For reaction **C1,** representing the formation of a cyclopropane core present in the antithrombotic therapeutic ticagrelor,^56^ PromPgb displays little to no selectivity for formation of the possible *cis*-**5** and *trans*-**5** cyclopropane products. Across both predicted libraries we find variants that are capable of selectively forming both possible diastereomers without significant losses in yield. Importantly, the synthesis of ticagrelor requires *trans*-**5**. To our delight, we identified variant VYIAIVFQ, which could deliver this diastereomer with a 9:1 preference for the *trans*- product. This emergence of stereodivergent behavior extended to other substrate classes: for the formation of alkylidene cyclopropanes ***Z*-6** and ***E*-6**, several variants not only display significantly improved yields for the PromPgb-favored *E*-isomer, but we also found that variant TAIFMVFQ inverts the selectivity toward formation of the *Z*-product (**Reaction C8** in **Scheme 2C**). Access to stereodivergent outcomes early in screening is particularly advantageous, as it provides multiple differentiated starting points from which independent DE campaigns can be initiated for desired stereoisomers, as would be the case for the synthesis of the ticagrelor cyclopropane precursor *trans***-5**.

There were 10 transformations in the full set of tested reactions which PromPgb could not catalyze with detectable yield (**Scheme 2A**). We were able to identify variants that catalyzed five of these previously inaccessible transformations. For reactions **N6** and **N7**, which involve C–H nitrene insertion using NH_2_OPiv at positions less activated or more hindered than for reaction **N3**, we identified a few variants in the first round of predictions that are capable of catalyzing these reactions in trace yield. The three designs capable of catalyzing reaction **N6** all follow the motif LYLFXXKQ, notably containing Phe to Lys substitutions at position 93, a site which was particularly recalcitrant to mutation for ITRs. This result highlights the importance of the balance of exploiting training data versus the exploration of sequence space in the ALDE algorithm. Additionally, we find that over a third of model-predicted mutants are capable of catalyzing the *α*- heteroatom C–H carbene insertion reactions **C14** and **C16**. Potentially, one could access reactions **C14** and **C16** through DE using a strategy known as a ‘substrate walk’ to enhance yield for the existing activity **C13**, which is mechanistically similar to these transformations, in the hope that would pick up activity for **C14** through **C16**. Here, however, ALDE directly proposes a diverse set of variants that are broadly competent for carbene transfer, derived solely from information encoded in a predefined reaction panel. As a result, directed evolution can be initiated with ALDE designs on substrates of greater synthetic or mechanistic interest without the need to first optimize an easier reaction (or intermediate substrate).

**Scheme 2.**
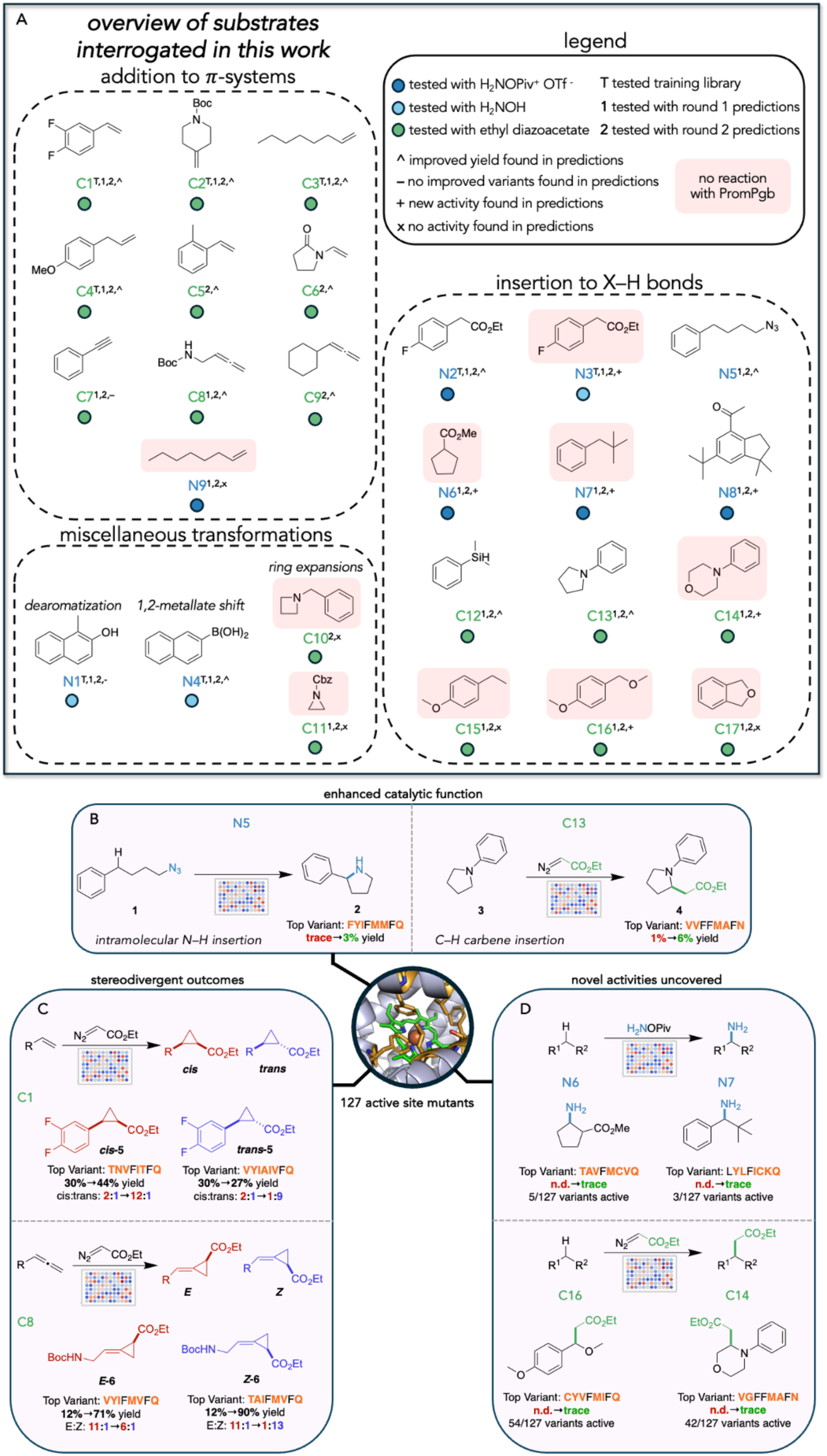
**(A)** Twenty-six carbene and nitrene transfer reactions representing electronically and sterically diverse acceptors were evaluated, including additions to π-systems and X–H insertion reactions. Symbols denote co- substrates for each reaction, rounds in which reaction was tested, and outcomes of library screening. **(B)** For nearly every tested reaction which PromPgb could perform, ALDE delivered improved variants. **(C)** Model- predicted variants unlock access to stereodivergent outcomes as an ensemble. **(D)** ALDE designs were capable of performing four of the nine assayed reactions which PromPgb could not perform.

## DISCUSSION

In this work, we present mr-ALDE, an implementation of ALDE that can be used to generate enzyme libraries which are capable of accessing a broader scope of chemical transformations. Starting from a thermostable protoglobin scaffold and a training set that captured double- and triple-mutant epistasis across a limited “basis set” of carbene and nitrene transfer reactions, ALDE generated multi-mutation variants that (1) retained basal activity across more training reactions than did multi-mutants seen in initial training, despite a higher mutational load, (2) frequently exceeded parent yields for at least one tested reaction, and (3) in several cases outperformed the parent on mechanistically distinct transformations not represented in initial training. In short, this data-driven approach effectively delivers enzyme variants that maintain functionality toward reactions represented in training while introducing diverse active-site environments that provide distinct solutions to related catalytic challenges.

Overall, we believe that mr-ALDE is a complementary approach to other experimental and computational strategies for generating enzyme libraries for finding detectable activity for a target reaction. While this method is inherently reliant on the construction of moderate sequence-function datasets, we believe that it presents some advantages in comparison to computational library generation methods which rely on zero-shot prediction or design. Firstly, this method is particularly applicable to the generation of enzyme libraries primed for the discovery of new-to-nature biocatalytic reactions, for which there is presumably little evolutionary information contained in existing protein sequence information. Methods such as FuncLib or the MODIFY software package developed by Ding and coworkers^14^ use sequence patterns from proteins found in nature to identify combinations of mutations which maintain stability and other biochemical properties likely important for function. However, new-to-nature chemistries often operate through catalytic principles and transition-state geometries that have not been sampled during natural evolution, as demonstrated by the amino acid identities of optimal variants in this study. Computational methods are particularly useful here because DE, while well suited for optimizing a single objective, is not suited to building libraries of proteins satisfying multiple objectives. Directed evolution also only considers a limited scope of single mutations when interrogating a diverse protein search space. By contrast, a direct MLDE-based approach leverages experimentally measured sequence–function data to identify variants that are empirically optimized for the desired reactivity, enabling the discovery of sequence solutions that lie outside the constraints imposed by evolutionary precedent.

Our findings regarding changes in population behavior in the first and second rounds of ALDE- predicted variants illustrate a practical strength of mr-ALDE: once an initial model is trained on a broad dataset, later rounds can focus on small, model-informed panels. Data collected from future engineering campaigns on members of ALDE-predicted libraries can be used to further update the model, enabling the continuous design of functional, diverse enzyme sequences. Similarly, the data generated through this ALDE framework have the potential to inform future machine learning-assisted directed evolution campaigns initiated from the designs generated in this work. In the course of this work, we assessed the function of 653 protoglobin variants harboring multiple active-site mutations against a panel of up to 26 distinct carbene and nitrene transfer reactions, resulting in over 7,000 unique sequence-function paired datapoints, which we have made publicly available at https://github.com/fhalab/mrALDE. To our knowledge this is the largest annotated dataset of non-native enzyme function, containing multiple substrate and reaction classes. This dataset may provide a useful reference for future MLDE training and validation studies, helping to inform broader efforts in data-driven enzyme engineering and design.

While we expect that mr-ALDE can be applied to other enzyme classes, we acknowledge that a limitation of this study is the focus on a single enzyme class that was particularly well positioned for success. Our efforts were initiated with a thermostable protein which we knew to be capable of promiscuously catalyzing both carbene and nitrene transfer chemistries: it would be difficult to perform mr-ALDE without a parent enzyme satisfying these criteria. Furthermore, our lab has extensive knowledge of engineering *Ape*Pgb variants, and we have previously applied ALDE to the active site of a related protoglobin homolog. As a result, we were able to confidently select active-site residues for the ALDE design space that influence non-native reactivity and exhibit epistatic effects. The generalizability of this approach thus remains to be demonstrated on systems where less prior knowledge is available for choosing residues to mutate.

Future studies will determine how well mr-ALDE can be extended to other enzyme and reaction classes and determine best practices for choosing training reaction sets, library designs, and ML-model parameters. Firstly, it is not clear how one should optimally design the original set of training reactions in mr-ALDE. It is possible that including fewer initial training reactions, or reactions spanning a greater chemical scope could lead to improved model training for predicting promiscuous enzyme variants. One could also envision a design framework in which the model is provided with the chemical details for training reactions. Such a model could possibly be used to explicitly suggest variants with activity for a specific substrate by combining existing function data with information on chemical similarity to the desired transformation. Additionally, we are curious to see whether mr-ALDE can be applied to broaden the capabilities of an enzyme to accommodate unnatural substrates for its native reaction mechanism, a common task in biocatalysis. Future studies will also require the investigation of datasets spanning diverse substrates within a *single* reaction class to generate libraries with expanded substrate promiscuity. Finally, in this study we selected the mean improvements in carbene and nitrene transfer activities as objective functions for multi-objective optimization. However, this encoding is apparently sensitive to variants seen in training which display atypically large improvements in activity for a particular reaction. This is evidenced by the strong representation of the amide-containing amino acids asparagine and glutamine at position 149 in model-predicted designs, likely due to the presence of these mutations in high-performing variants for reaction **N4**. We anticipate that subsequent studies must be conducted to develop ideal objective-function encodings which can avoid bias from outliers seen in model training data and to more effectively deliver diverse enzyme libraries.

Altogether, the discovery of 127 catalytically active variants encompassing both carbene and nitrene transfer reactions illustrates the efficiency of coupling machine learning with laboratory evolution principles. Where conventional directed evolution would have required several discrete engineering campaigns to achieve similar functional coverage, mr-ALDE accomplishes this in a single, integrated workflow. The ability of mr-ALDE to rapidly generate stable, mechanistically diverse, and functionally enriched enzyme libraries highlights its potential for accelerating the discovery of new biocatalytic activities across an expanding range of chemical transformations.^57^

## Contributions

R.G.L.: conceptualization, methodology, investigation, analysis, writing – original draft, writing – editing

J.Y. : conceptualization, methodology, software, investigation, analysis, writing – original draft, writing – editing

Z.Z.: investigation, writing – original draft, writing – editing

F.H.A.: resources, writing – editing, supervision, funding

## Acknowledgements

This work was supported by the U.S. Army Research Office cooperative agreement for the Institute for Collaborative Biotechnologies (W911NF-19-2-0026 to F.H.A.). This report was prepared as an account of work sponsored by an agency of the United States Government. Neither the United States Government nor any agency thereof, nor any of their employees, makes any warranty, express or implied, or assumes any legal liability or responsibility for the accuracy, completeness, or usefulness of any information, apparatus, product, or process disclosed, or represents that its use would not infringe privately owned rights. Reference herein to any specific commercial product, process, or service by trade name, trademark, manufacturer, or otherwise does not necessarily constitute or imply its endorsement, recommendation, or favoring by the United States Government or any agency thereof. The views and opinions of authors expressed herein do not necessarily state or reflect those of the United States Government or any agency thereof. This work was also supported by the NSF Division of Chemical, Bioengineering, Environmental and Transport Systems (CBET 1937902). J.Y. and R.G.L. are partially supported by National Science Foundation Graduate Research Fellowships and J.Y. is supported by the Google PhD Fellowship. The authors thank Kathleen M. Sicinski, Jae L. Kennemur, Yueming Long, Edwin Alfonzo, and Deirdre M. Hanley for providing guidance on analytical method development for training reactions. The authors thank Raul Astudillo for providing guidance on the use and implementation of expected hypervolume improvement. The authors thank Francesca Zhoufan-Li for help with data visualization and analysis.

## SUPPORTING INFORMATION

Information about materials, experimental methods, computational methods, compound characterization data and supplementary data can be found in the included supporting information. All experimental and simulation data that support the findings of this study are available at https://github.com/fhalab/mrALDE. All code that accompanies this study is available at https://github.com/fhalab/mrALDE under the MIT license.

