## Supporting Information for "Machine Learning-Assisted Evolution of Broadly Functional Enzyme Libraries"

### Contents


| | |
|---|---|
| General Information | S3 |
| Cloning and Sequence Design | S5 |
| Cloning Protocols and Results | S11 |
| Protoglobin Library Screening Protocols | S54 |
| Library Screening Details and Data | S57 |
| Computed Summary Statistics | S97 |
| Machine Learning Details | S98 |
| Quantitative Analysis of Select PromPgb Variants | S101 |
| Preparation of Calibration Curves for Analytical Yield Determination | S103 |
| Preparation of Authentic Standards | S106 |
| Preparative-Scale Enzymatic Reactions | S117 |
| NMR Spectra of Authentic Standards and Isolated Products | S120 |
| References | S130 |

### General Information

**Safety Statement:**

All chemical transformations were performed in a well-ventilated fume hood to avoid inhalation and exposure. Other than that, no unexpected or unusually high safety concerns were raised with these methods. Safety notes for individual synthetic procedures will be documented alongside the procedure.

**Chemicals:**

All chemical transformations were performed in a well-ventilated fume hood to avoid inhalation and exposure to chemicals. Reagents and solvents were obtained commercially (Sigma-Aldrich, Alfa Aesar, VWR, Fisher Scientific, Matrix Scientific, Oakwood Chemical, TCI America, and other suppliers) and used without prior purification unless otherwise stated.

**Instrumentation:**

Organic solutions were concentrated under reduced pressure on an IKA RV 10 rotary evaporator. Thin-layer chromatography (TLC) was performed on commercial Millipore Silica Gel 60 plates containing the F254 fluorescent indicator. Visualization of the developed chromatographs was performed by irradiation with UV light or by treatment with an appropriate TLC staining solution (e.g., Ceric Ammonium Molybdate, $KMnO_4$, or Bromocresol Green) followed by heating if necessary. Chromatographic purification was accomplished by flash chromatography using a Biotage Isolera One instrument.

All NMR spectra were obtained at the Caltech Liquid NMR Facility. NMR spectra were collected on a Bruker Prodigy 400 MHz instrument equipped with a cryoprobe operating at 400 MHz and 101 MHz for $^{1}H$ and $^{13}C$, respectively. $^{1}H$ spectra are referred to residual $CDCl_3$ solvent signals referenced at δ 7.26 ppm. Data for $^{1}H$ NMR are reported as follows: chemical shift (δ ppm), integration, multiplicity (s = singlet, d = doublet, t = triplet, q = quartet, p = pentad, sext = sextet, hept = heptet, m = multiplet, br s = broad singlet), and coupling constant (Hz).

Gas chromatography (GC) was performed on an Agilent Technologies 7820A GC system equipped with a split-mode capillary injection system. For achiral analyses, an Agilent J&W HP-5 Column was used as the stationary phase. Reaction systems displaying sufficient yield were subsequently analyzed with a flame-ionization detector. When target analytes were only produced in trace yield, samples were analyzed by an Agilent Technologies 5977B mass spectrometer.

Liquid chromatography (LC) was performed on an Agilent Technologies 1260 Infinity HPLC-MS system (Agilent 6120 quadrupole mass spectrometer) equipped with an Agilent C18 column (Poroshell 120 ESC18, 4.6 x 50 mm, 2.7-*μ*m packing) with a Poroshell 120 guard column (2.7-*μ*m packing, 2.1 x 5 mm). Water and acetonitrile modified with 0.1% acetic acid were used as eluents.

### Cloning and Sequence Design

#### Relevant Protoglobin Sequences:

**Table S1.** The amino acid sequences of wild-type *Ape*Pgb and the previously engineered variant, PromPgb. Residues in each sequence highlighted in yellow are the sites which were investigated in this work for wet-lab experimentation: L59, W62, F63, F73, L86, S90, F93, and S149 in PromPgb.

| Protein Variant | Amino Acid Sequence |
|---|---|
| wild-type *Aeropyrum pernix* Protoglobin (*Ape*Pgb) | MTPSDIPGYDYGRVEKSPITDLEFDLLKKTVM<br>LGEKDVMYLKKACDVLKDQVDEILDLWYG<br>WVASNEHLIYYFSNPDTGEPIKEYLERVRARF<br>GAWILDTTCRDYNREWLDYQYEVGLRHHRS<br>KKGVTDGVRTVPHIPLRYLIAFIYPITATIKPFL<br>AKKGGSPEDIEGMYNAWFKSVVLQVAIWSHP<br>YTKENDW |
| *Aeropyrum pernix* Protoglobin D10G C45Y L56I W59L Y60G V63F R90S F145G I149S F156L Y189H (PromPgb) | MTPSDIPGYGYGRVEKSPITDLEFDLLKKTVM<br>LGEKDVMYLKKAYDVLKDQVDEIIDLLGGW<br>FASNEHLIYYFSNPDTGEPIKEYLERVSARFGA<br>WILDTTCRDYNREWLDYQYEVGLRHHRSKK<br>GVTDGVRTVPHIPLRYLIAGIYPSTATIKPLLA<br>KKGGSPEDIEGMYNAWFKSVVLQVAIWSHPH<br>TKENDW |

**Table S2.** DNA Sequence of PromPgb, including C-terminal 6xHis-tag and stop codon. The region highlighted in green represents the region of the gene in which exact, predicted sets of mutations were incorporated in an oligo pool.

| |
|---|
| ATGACTCCCTCGGACATCCCGGGATATGGTTATGGGCGTGTCGAGAAGTCACCCATCACG<br>GACCTTGAGTTTGACCTTCTGAAGAAGACTGTCATGTTAGGTGAAAAGGACGTAATGTAC<br>TTGAAAAAGGCGTATGACGTTCTGAAAGATCAAGTTGATGAGATCATTGACTTGCTGGGT<br>GGTTGGTTTGCATCAAATGAGCATTTGATTTATTACTTCTCCAATCCGGATACAGGAGAGC<br>CTATTAAGGAATACCTGGAACGTGTAAGCGCTCGCTTTGGAGCCTGGATTCTGGACACTAC<br>CTGCCGCGACTATAACCGTGAATGGTTAGACTACCAGTACGAAGTTGGGCTTCGTCATCAC<br>CGTTCAAAGAAAGGGGTCACAGACGGAGTACGCACCGTGCCCCATATCCCACTTCGTTAT<br>CTTATCGCAGGTATCTATCCTAGTACCGCCACTATCAAGCCACTTTTGGCTAAGAAAGGTG<br>GCTCTCCGGAAGACATCGAAGGGATGTACAACGCTTGGTTCAAGTCTGTAGTTTTACAAGT<br>TGCCATCTGGTCACACCCTCATACTAAGGAGAATGACTGGCTCGAGCACCACCACCACCA<br>CCACTGAG |

**Table S3.** Codons utilized when ALDE-predicted mutations were incorporated into the ParLQ DNA sequence. Codons which are generally recognized as the most prevalently found in the *E. coli* genome were selected.

| **Amino Acid** | **Codon** |
|---|---|
| A | GCG |
| C | TGC |
| D | GAT |
| E | GAA |
| F | TTT |
| G | GGC |
| H | CAT |
| I | ATT |
| K | AAA |
| L | CTG |
| M | ATG |
| N | AAC |
| P | CCG |
| Q | CAG |
| R | CGT |
| S | AGC |
| T | ACC |
| V | GTG |
| W | TGG |
| Y | TAT |

**Nomenclature for Variant Naming:**

Single-site mutants are named using the standard nomenclature: (original AA)(site)(new AA). The mutant L59F refers to a variant of PromPgb mutated from leucine to phenylalanine at position 59. Eight-site multi-mutants are named as a string of the eight amino acids to which the positions of interest have been mutated. The variant IYVAIIKQ refers to a variant of PromPgb bearing the mutations F59I, W62Y, F63V, F73A, L86I, S90I, F93K, and S149Q. In this nomenclature, PromPgb is named LWFFLSFS.

**Primer Design for Site-Saturation Mutagenesis (SSM):**

*General Cloning Primers*

**Table S4.** Primers 007 and primers 008 are used to generate amplicons of linearized pET-22b(+)[1] backbone. Primers 005 and 006 were used with primers internal to the protoglobin gene (**Table S5**) to generate mutant protoglobin genes. All primers were ordered from IDT (Coralville, IA).

| Primer Name | Direction | Sequence | Description |
|---|---|---|---|
| 005 | Forward | 5'-gaaataattttgtttaactttaagaaggagatatacatatg-3' | Upstream of N-term, anneals with 007 |
| 006 | Reverse | 5'-gccggatctcagtggtggtggtggtggtgctcgag-3' | Downstream of C-term, anneals with 008 |
| 007 | Reverse | 5'-catatgtatatctccttcttaaagttaaacaaaattatttc-3' | Upstream of N-term, anneals with 005 |
| 008 | Forward | 5'-ctcgagcaccaccaccaccaccactgagatccggc-3' | Downstream of C-term, anneals with 006 |

*Cloning Primers for Site-Saturation Mutagenesis*

**Table S5.** Primers containing degenerate codons at sites of interest for mutagenesis. Primers were used in conjunction with either primer 005 or 006 (**Table S4**) to generate mutant protoglobin genes. All primers were ordered from IDT (Coralville, IA). Abbreviations: SSM – site saturation mutagenesis, dSSM – double site saturation mutagenesis, tSSM – triple site saturation mutagenesis.

| **Primer Name** | **Sequence** | **Description** |
|---|---|---|
| 59_fwd | 5'-attgacttgNNKggtggttggtttgcatcaaatgagc-3' | SSM for site 59 |
| 59_rev | 5'-atttgatgcaaaccaaccaccMNNcaagtcaatgatctcatc-3' | SSM for site 59 |
| 73_fwd | 5'-gcatttgatttattacNNKtccaatccggatacaggagag-3' | SSM for site 73 |
| 73_rev | 5'-tgtatccggattggaMNNgtaataaatcaaatgctcatttg-3' | SSM for site 73 |
| 86_fwd | 5'-ttaaggaatacNNKgaacgtgtaagcgctcgctttg-3' | SSM for site 86 |
| 86_rev | 5'-aagcgagcgcttacacgttcMNNgtattccttaatag-3' | SSM for site 86 |
| 90_fwd | 5'-gcctattaaggaatacctggaacgtgtaNNKgctcgctttggagcctgg-3' | SSM for site 90 |
| 90_rev | 5'-ccaggctccaaagcgagcMNNtacacgttccaggtattccttaataggc-3' | SM for site 90 |
| 93_fwd | 5'-ctggaacgtgtaagcgctcgcNNKggagcctggattctggacac-3' | SSM for site 93 |
| 93_rev | 5'-cagaatccaggctccMNNgcgagcgcttacacgttccaggtattc-3' | SSM for site 93 |
| 149_fwd | 5'-tcgcaggtatctatcctNNKaccgccactatcaagccac-3' | SSM for site 149 |
| 149_rev | 5'-gatagtggcggtMNNaggatagatacctgcgataagataacgaag-3' | SSM for site 149 |
| 59+62_fwd | 5'-atgagatcattgacttgNNKggtggtNNKtttgcatcaaatgagc-3' | dSSM for sites 59 and 62 |
| 59+62_rev | 5'-tttgatgcaaaMNNaccaccMNNcaagtcaatgatctcatcaac-3' | dSSM for sites 59 and 62 |
| 59+63_fwd | 5'-atgagatcattgacttgNNKggtggttggNNKgcatcaaatgagc-3' | dSSM for sites 59 and 63 |
| 59+63_rev | 5'-tttgatgcMNNccaaccaccMNNcaagtcaatgatctcatcaac-3' | dSSM for sites 59 and 63 |

| | | |
|---|---|---|
| 62+63_fwd | 5'-gacttgctgggtggtNNKNNKgcatcaaatgagcatttgatttattacttc-3' | dSSM for sites 62 and 63 |
| 62+63_rev | 5'-aatcaaatgctcatttgatgcMNNMNNaccacccagcaagtcaatgatc-3' | dSSM for sites 62 and 63 |
| 63+73_fwd | 5'-gtggttggNNKgcatcaaatgagcatttgatttattacNNKtccaatccggatacaggag-3' | dSSM for sites 63 and 73 |
| 63+73_rev | 5'-ggattggaMNNgtaataaatcaaatgctcatttgatgcMNNccaaccacccagcaagtc-3' | dSSM for sites 63 and 73 |
| 86+90_fwd | 5'-ctattaaggaatacNNKgaacgtgtaNNKgctcgctttggagcctg-3' | dSSM for sites 86 and 90 |
| 86+90_rev | 5'-caggctccaaagcgagcMNNtacacgttcMNNgtattccttaatag-3' | dSSM for sites 86 and 90 |
| 86+93_fwd | 5'-tattaaggaatacNNKgaacgtgtaagcgctcgcNNKggagcctggattctggac-3' | dSSM for sites 86 and 93 |
| 86+93_rev | 5'-aatccaggctccMNNgcgagcgcttacacgttcMNNgtattccttaataggctc-3' | dSSM for sites 86 and 93 |
| 90+93_fwd | 5'-ggaacgtgtaNNKgctcgcNNKggagcctggattctggacactacctgc-3' | dSSM for sites 90 and 93 |
| 90+93_rev | 5'-ccagaatccaggctccMNNgcgagcMNNtacacgttccaggtattccttaatag-3' | dSSM for sites 90 and 93 |
| 59+62+63_fwd | 5'-caagttgatgagatcattgacttgNNKggtggtNNKNNKgcatcaaatgagcatttg-3' | tSSM for sites 59, 62, and 63 |
| 59+62+63_rev | 5'-gctcatttgatgcMNNMNNaccaccMNNcaagtcaatgatctcatcaacttgatc-3' | tSSM for sites 59, 62, and 63 |

*Cloning Primers for Oligo Libraries*

**Table S6.** Primers used for incorporating adaptors to the ends of oligo fragments for incorporation into the pET-22b(+) vector.

| Primer Name | Sequence | Description |
|---|---|---|
| oligo_fwd | 5'-gttctgaaagatcaagttgatgagatcattgacttg-3' | Anneals to multi-mutant oligo fragments |
| oligo_rev | 5'-ccaaaagtggcttgatagtggcggt-3' | Anneals to multi-mutant oligo fragments |
| pET_BB_fwd | 5'-accgccactatcaagccacttttgg-3' | Anneals to base pairs internal to PromPgb |
| pET_BB_rev | 5'-caagtcaatgatctcatcaacttgatctttcagaac-3' | Anneals to base pairs internal to PromPgb |

### Cloning Protocols and Results

#### Protocols for the Cloning of Random PromPgb Variants:

*Cloning for Site-Saturation Mutagenesis (SSM)*

Chemically competent *Escherichia coli* (*E. coli*) cells (T7 Express Competent *E. coli*) were purchased from New England Biolabs (NEB, Ipswich, MA). Additionally, Phusion polymerase and *Dpn*I were purchased from NEB. SSM experiments were performed using primers bearing degenerate codons (NNK) using a modified QuikChange™ protocol (**Tables S4** and **S5**).[2]

The PCR conditions were as follows (final concentrations): Phusion HF Buffer 1x, 0.2 mM dNTPs each, 0.5 μM of forward primers, 0.5 μM reverse primer, and 0.02 U/μL of Phusion polymerase. The standard Phusion PCR protocol was used.[3] Upon completion of PCRs, the remaining template was digested with *Dpn*I. Gel purification was performed with a Zymoclean Gel DNA Recovery Kit (Zymo Research Corp, Irvine, CA). The purified PCR product was then assembled using the Gibson assembly protocol.[4]

**Table S7.** Primer combinations for the generation of PCR amplicons for the construction of expression plasmids containing mutagenized PromPgb variants. '*site*' refers to the specific site or set of sites being saturated.

| **Fragment Name** | **Forward Primer** | **Reverse Primer** |
|---|---|---|
| SSM_Frag1 | 005 | *site*_rev |
| SSM_Frag2 | *site*_fwd | 006 |
| pET_Backbone | 008 | 007 |

*Construction of Random Multi-Mutant Libraries*

The primers described in **Table S5** were designed to provide access to all 18 of the random multi-mutation variant libraries (**Table S8**). These libraries represent the set of variants used to initially train machine learning models. Libraries requiring multiple mutagenesis steps to construct the final multi-mutants were generated as follows: Site saturation or multi-site saturation mutagenesis was performed on the first set of sites within the defined library according to the above PCR and assembly protocol (*Cloning for Site-Saturation Mutagenesis (SSM)*). The assembly products obtained were used to transform T7 Express Competent *E. coli* (High Efficiency) cells following the protocol recommended by the manufacturer. Upon heat-shock, freshly transformed *E. coli* cells were recovered in 0.4 mL Luria-Bertani medium (LB) (Research Products Int.) at 37 ºC with shaking at 220 rpm for 30 minutes. For libraries requiring only a single mutagenesis step, this transformation mixture was directly applied to the protocol described below for plating on LB-Amp agar plates. For libraries requiring two mutagenesis steps, 50 µL was transferred into 6 mL of LB with 100 µg/mL ampicillin (LB-Amp) in a 15-mL culture tube. This culture was allowed to shake overnight at 37 °C and 220 rpm. The following morning, this library overnight culture was miniprepped using a QIAprep Spin Miniprep Kit (Qiagen, Hilden, Germany). The miniprepped plasmid DNA pool was used as the new template for mutagenesis with the primers for the remaining sites in the random library. T7 Express Competent *E. coli* were transformed with the Gibson products for the new multi-site library using the recommended protocol. Upon heat-shock transformation, the freshly transformed *E. coli* cells were recovered in 0.4 mL LB medium at 37 ºC with shaking at 220 rpm for 30 minutes, yielding a transformation mixture with *E. coli* harboring the final multi-site mutagenesis library.

Transformation mixtures with *E. coli* harboring one of the final desired libraries were plated on LB-agar plates with 100 μg/mL ampicillin (LB-Amp agar plates). The plates were incubated overnight at 37 °C until colony formation was observed. For each of the 18 libraries, single colonies from LB-Amp agar plates were picked with sterilized toothpicks to individually inoculate the wells of a 2-mL 96-well deep-well plate charged with 400 μL of LB-Amp. The plates were incubated at 37 ºC and shaken at 220 rpm for 16–18 hours. 96-Well deep-well plates were shaken in an INFORS HT Multitron Shaker in all instances. The following morning, 50 μL of preculture from each well were added to the wells of a 96-well flat-bottom tissue culture plate (ThermoFisher) preloaded with 50 μL of 50% glycerol solution. These glycerol stocks were stored at -80 °C for future inoculation. Additionally, the sequences of protoglobin genes contained in every well were sequenced using LevSeq sequencing (**Figures S1–S20**).[5]

**Table S8.** Combinatorial libraries tested used to collect training data and the assembly methods used in their generation. Residue distances were calculated from C$\alpha$ positions in a homology model of PromPgb co-folded with heme using AlphaFold3.[6]

| **Library** | **Sites** | **Residue Distance (Å)** | **Construction Steps** |
|---|---|---|---|
| 1 | 59, 62 | 5.0 | single dSSM step with 59+62 primers |
| 2 | 62, 63 | 3.8 | single dSSM step with 62+63 primers |
| 3 | 63, 73 | 9.0 | single dSSM step with 63+73 primers |
| 4 | 73, 86 | 6.3 | sSSM with 73 primers, then sSSM with 86 primers |
| 5 | 86, 90 | 5.9 | single dSSM step with 86+90 primers |
| 6 | 90, 93 | 4.8 | single dSSM step with 90+93 primers |
| 7 | 93,149 | 11.3 | sSSM with 93 primers, then sSSM with 149 primers |
| 8 | 59, 149 | 9.8 | sSSM with 59 primers, then sSSM with 149 primers |
| 9 | 59, 73 | 12.8 | single dSSM step with 59+73 primers |
| 10 | 73, 90 | 11.2 | sSSM with 73 primers, then sSSM with 90 primers |
| 11 | 86, 93 | 10.1 | single dSSM step with 86+93 primers |
| 12 | 59, 93 | 9.0 | sSSM with 59 primers, then sSSM with 93 primers |
| 13 | 62, 63, 86 | 3.8 (62-63), 11.8 (62-86), 9.2 (63-86) | dSSM with 62+63 primers, then sSSM with 86 primers |
| 14 | 59, 62, 63 | 5.0 (59-62), 6.3 (59-63), 3.8 (62-63) | single tSSM step with 59+62+63 primers |
| 15 | 59, 63, 90 | 6.3 (59-63), 7.6 (59-90), 10.7 (63-90) | dSSM with 59+63 primers, then sSSM with 90 primers |
| 16 | 59, 90, 93 | 7.6 (59-90), 9.0 (59-93), 4.8 (90-93) | dSSM with 90+93 primers, then sSSM with 59 primers |
| 17 | 62, 63,149 | 3.8 (62-63), 11.2 (62-149), 12.6 (63-149) | dSSM with 62+63 primers, then sSSM with 149 primers |
| 18 | 86, 90, 149 | 5.9 (86-90), 17.1 (86-149), 13.9 (90-149) | dSSM with 86+90 primers, then sSSM with 149 primers |

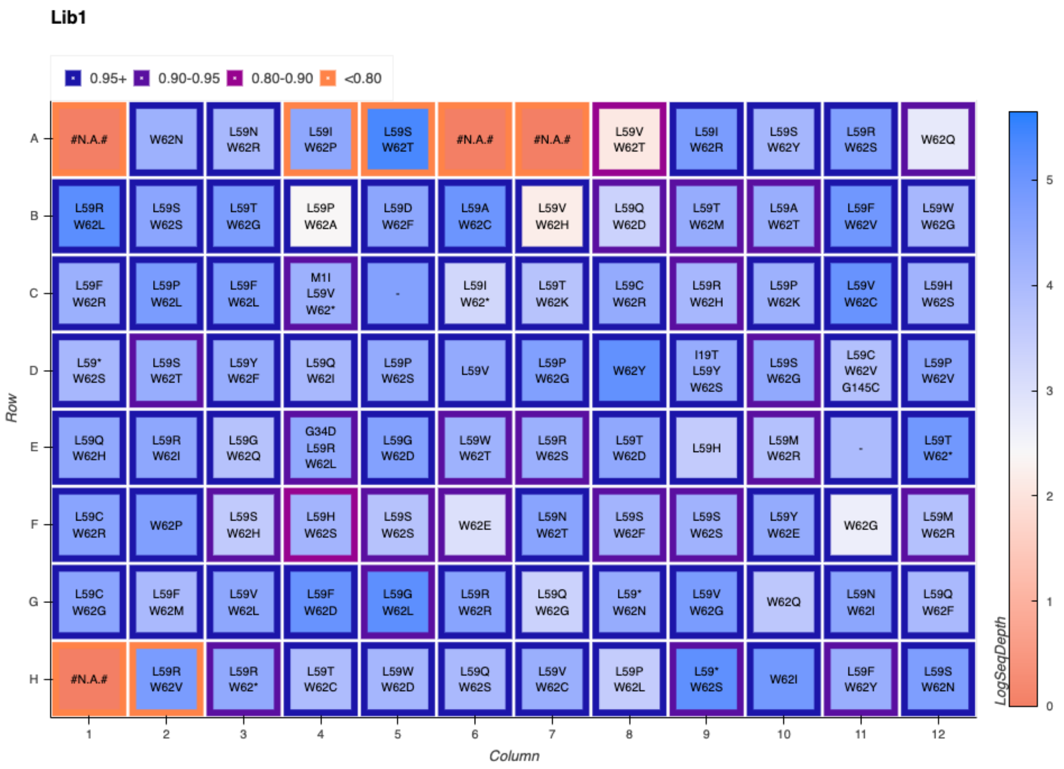


**Figure S1.** LevSeq sequencing plate map for random variants picked from library 1.

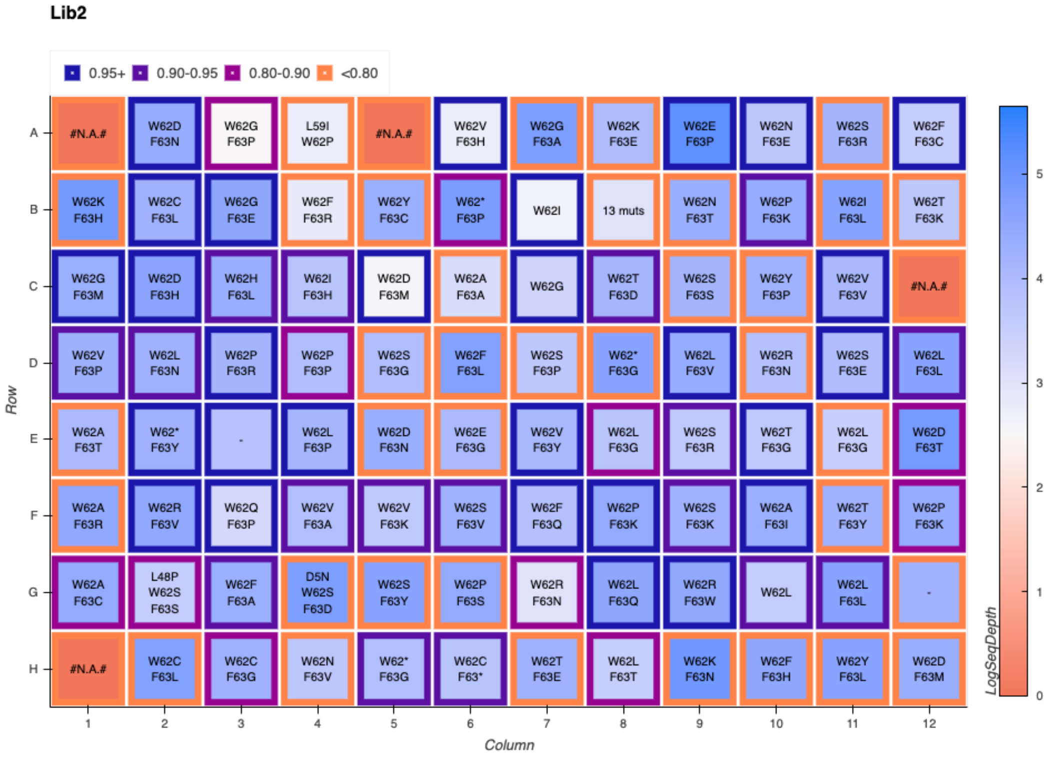


**Figure S2.** LevSeq sequencing plate map for random variants picked from library 2.

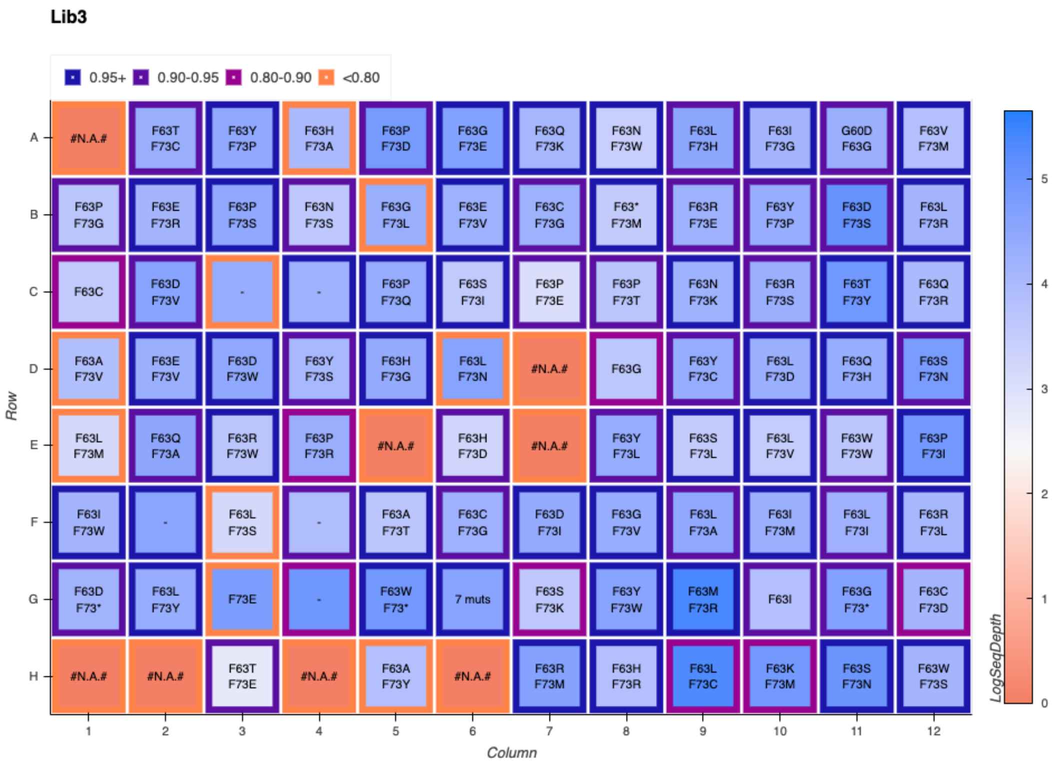


**Figure S3.** LevSeq sequencing plate map for random variants picked from library 3.

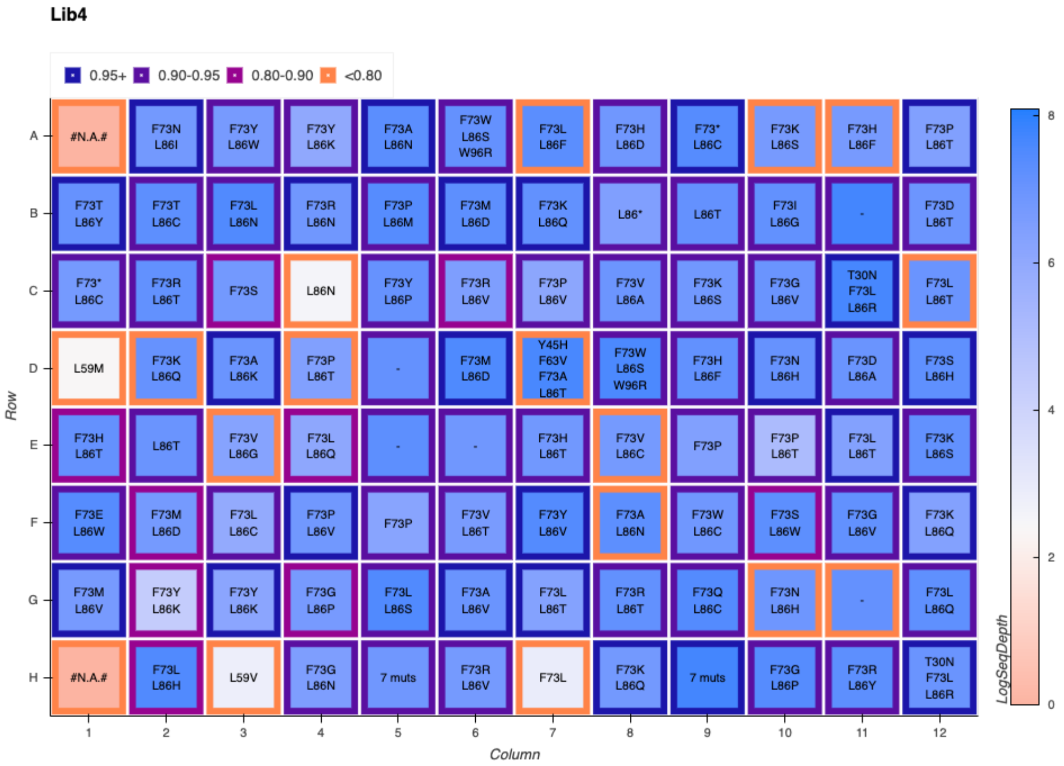


**Figure S4.** LevSeq sequencing plate map for random variants picked from library 4.

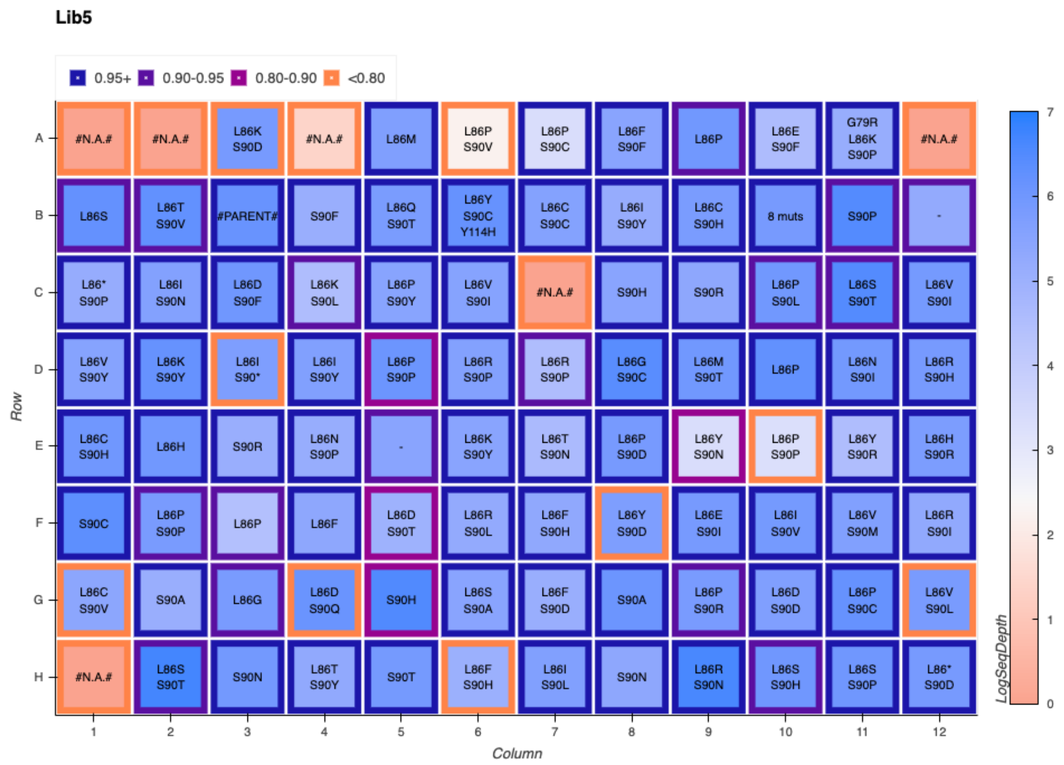


**Figure S5.** LevSeq sequencing plate map for random variants picked from library 5.

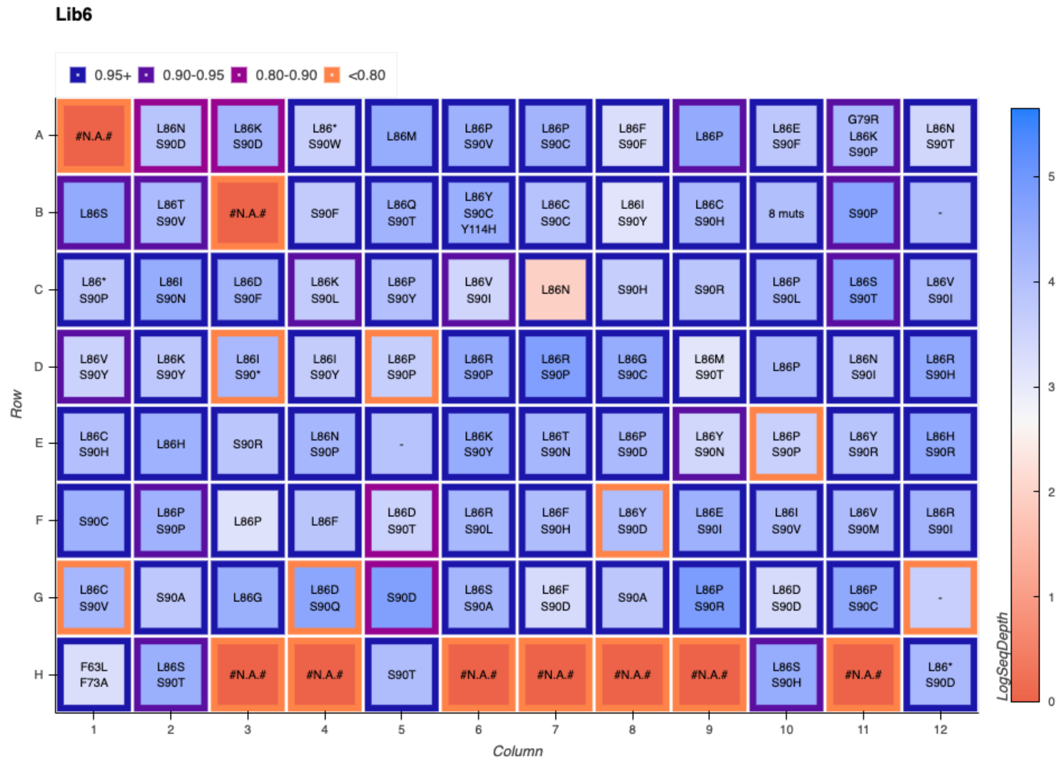


**Figure S6.** LevSeq sequencing plate map for random variants picked from library 6.

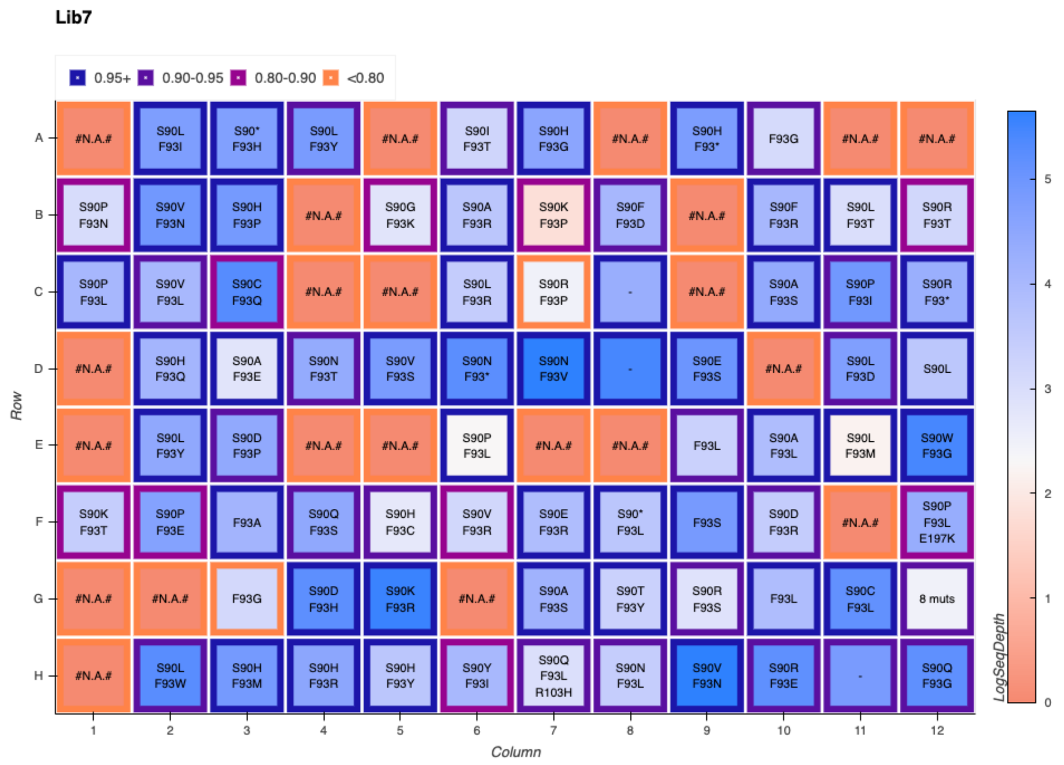


**Figure S7.** LevSeq sequencing plate map for random variants picked from library 7.

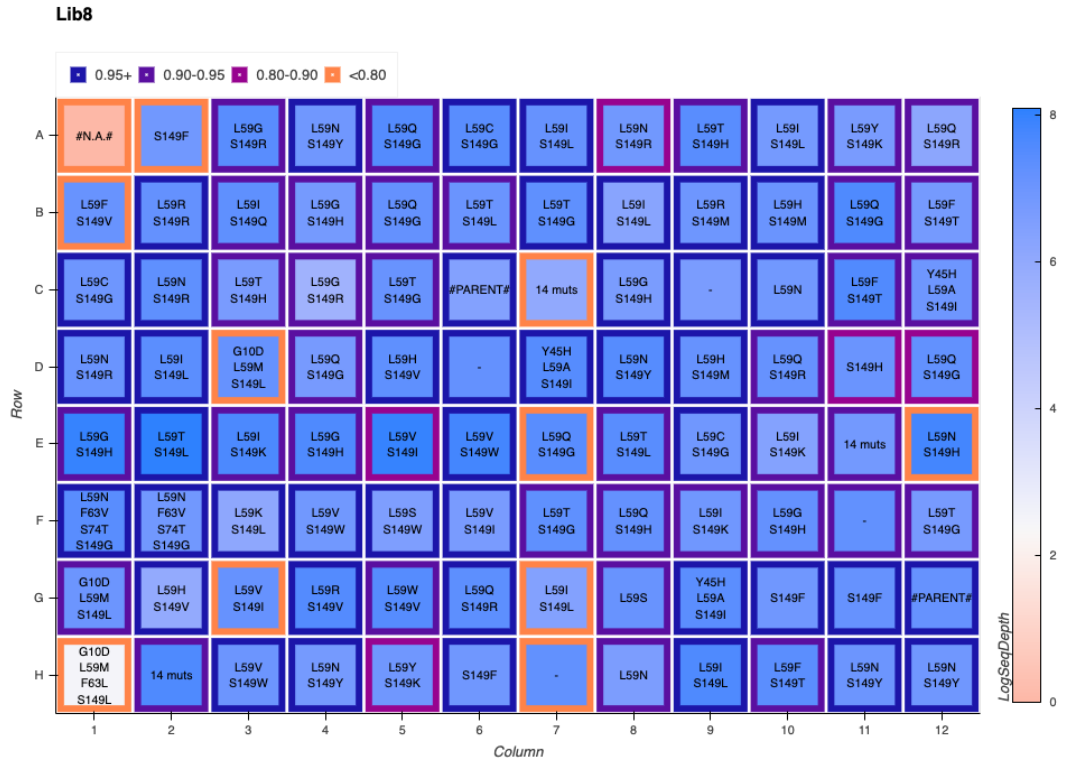


**Figure S8.** LevSeq sequencing plate map for random variants picked from library 8.

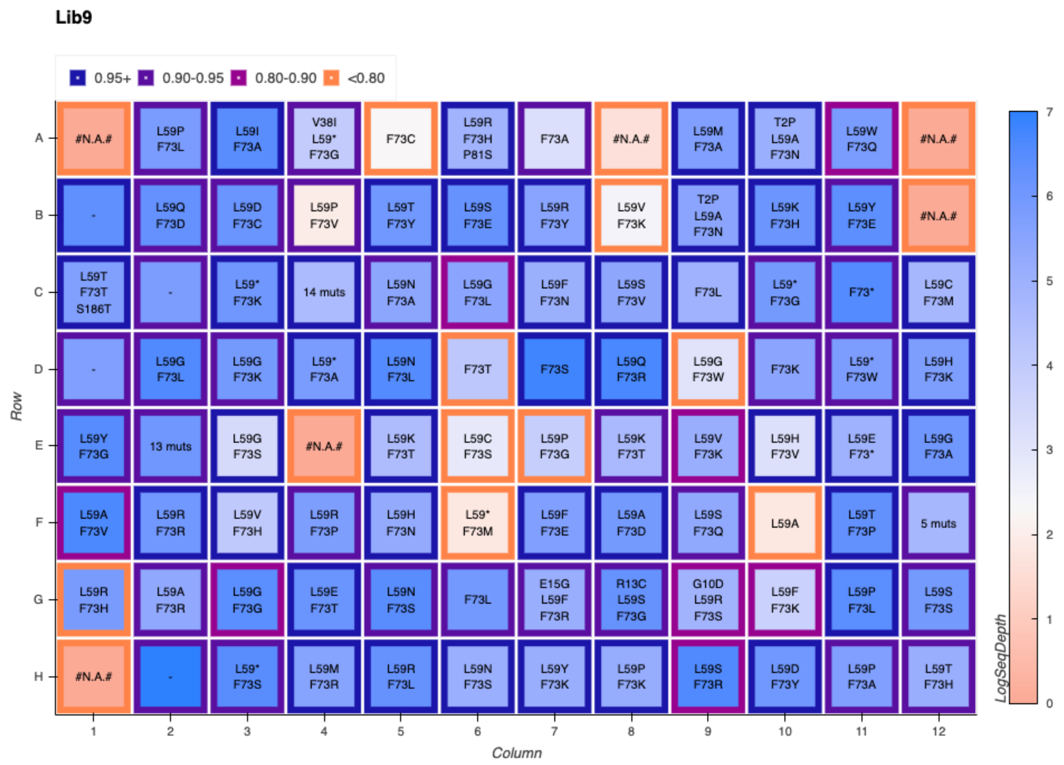


**Figure S9.** LevSeq sequencing plate map for random variants picked from library 9.

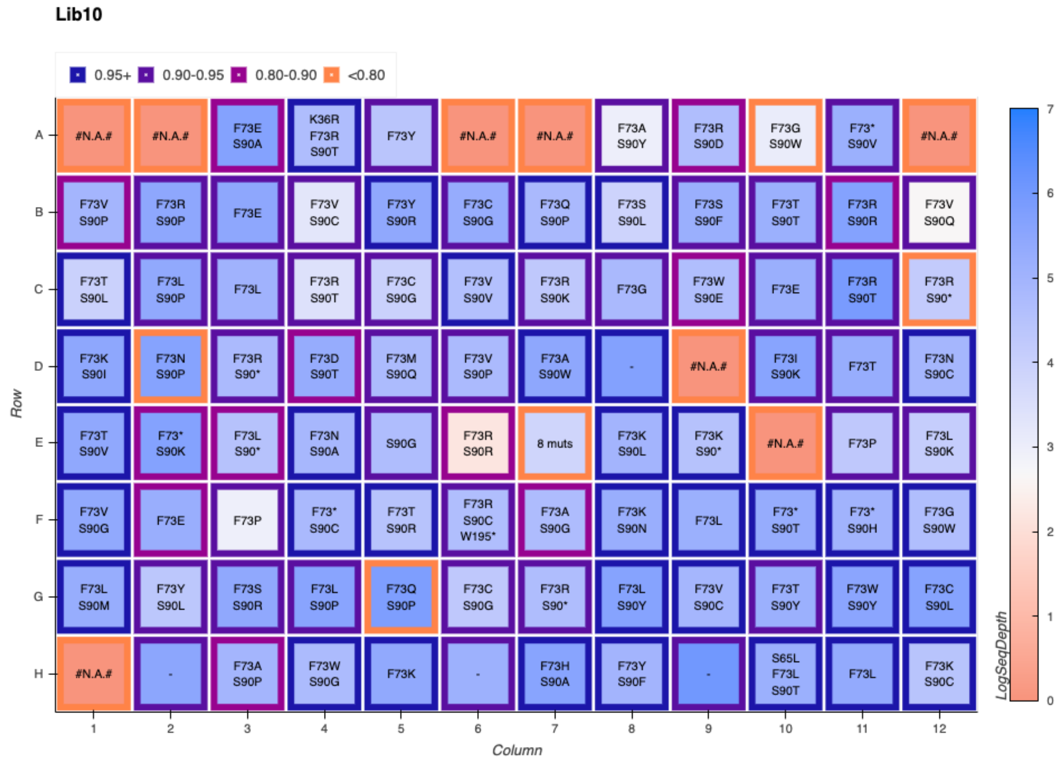


**Figure S10.** LevSeq sequencing plate map for random variants picked from library 10.

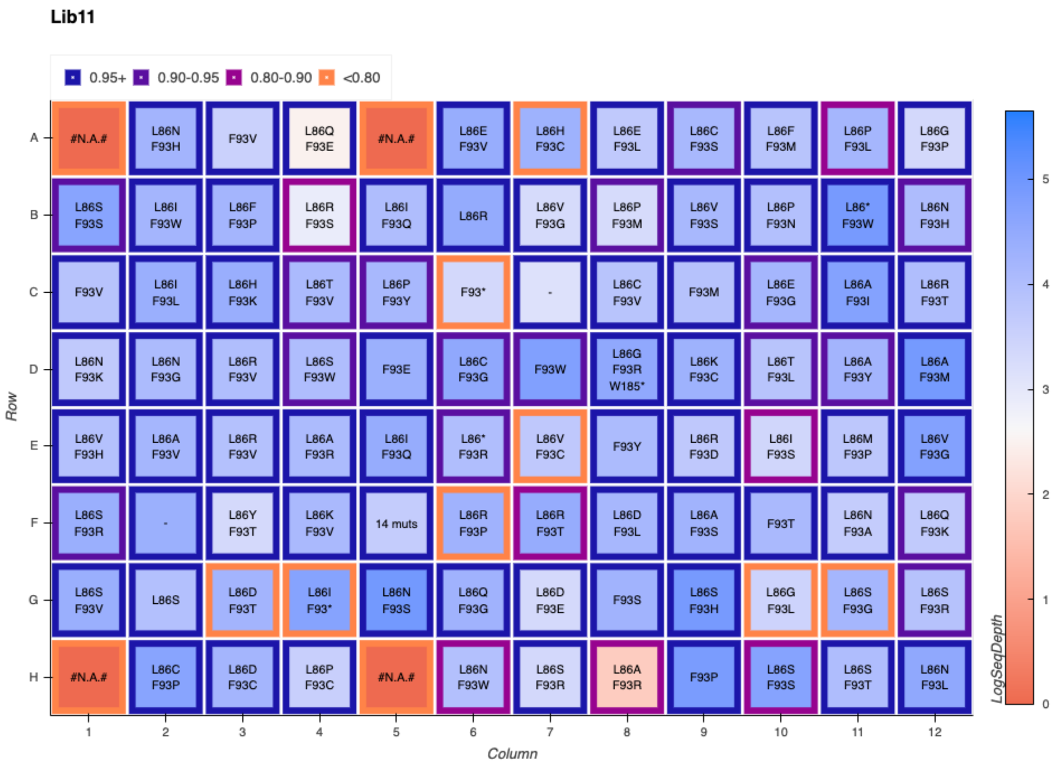


**Figure S11.** LevSeq sequencing plate map for random variants picked from library 11.

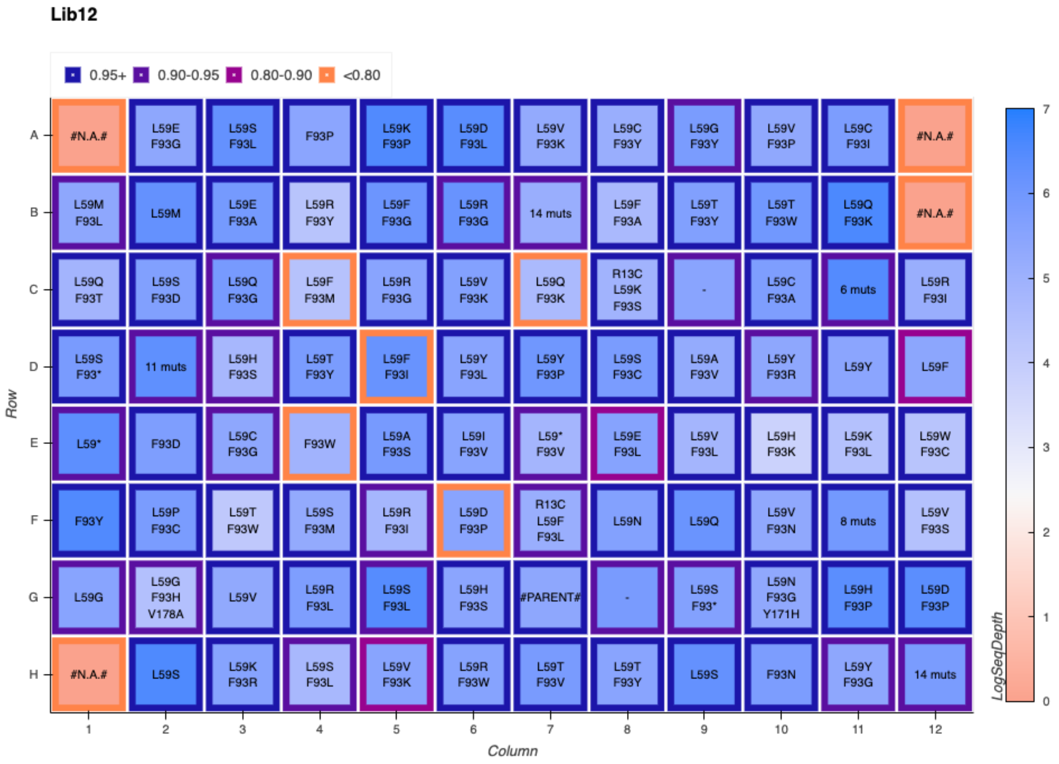


**Figure S12.** LevSeq sequencing plate map for random variants picked from library 12.

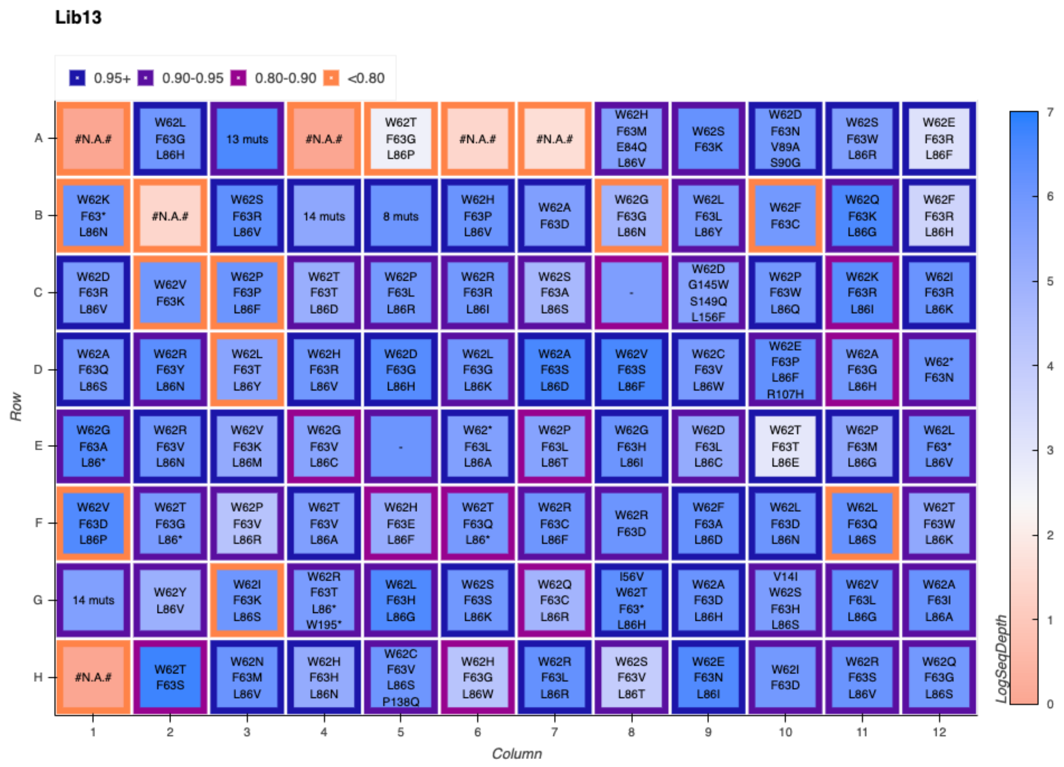


**Figure S13.** LevSeq sequencing plate map for random variants picked from library 13.

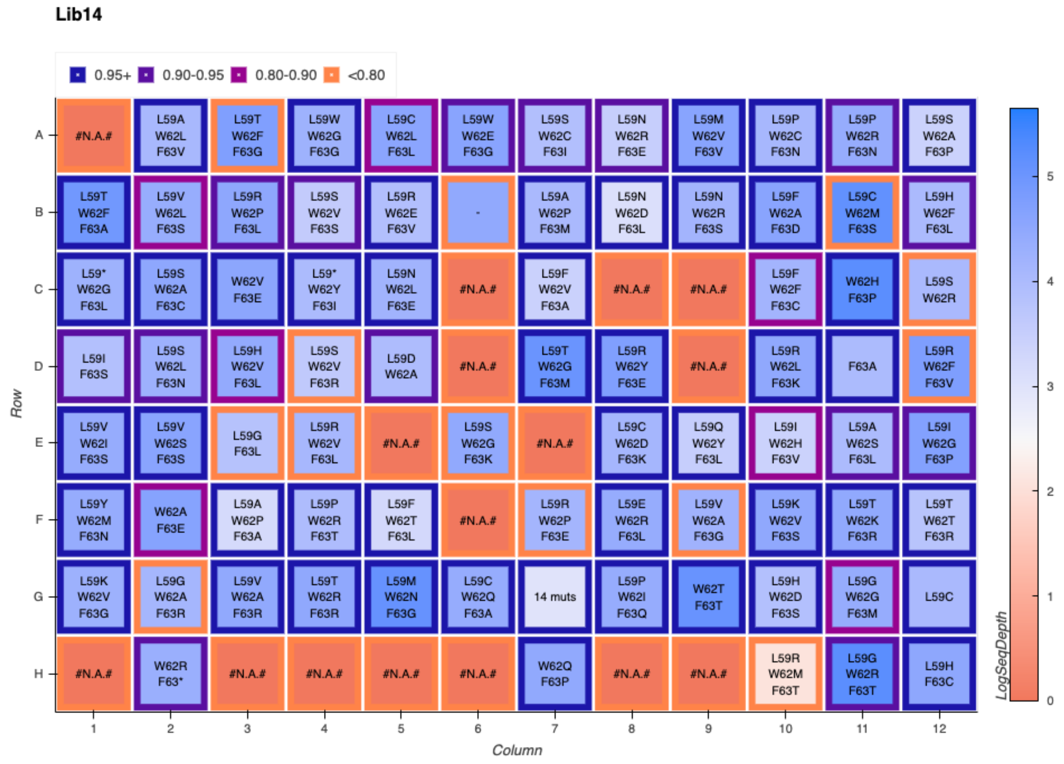


**Figure S14.** LevSeq sequencing plate map for random variants picked from library 14.

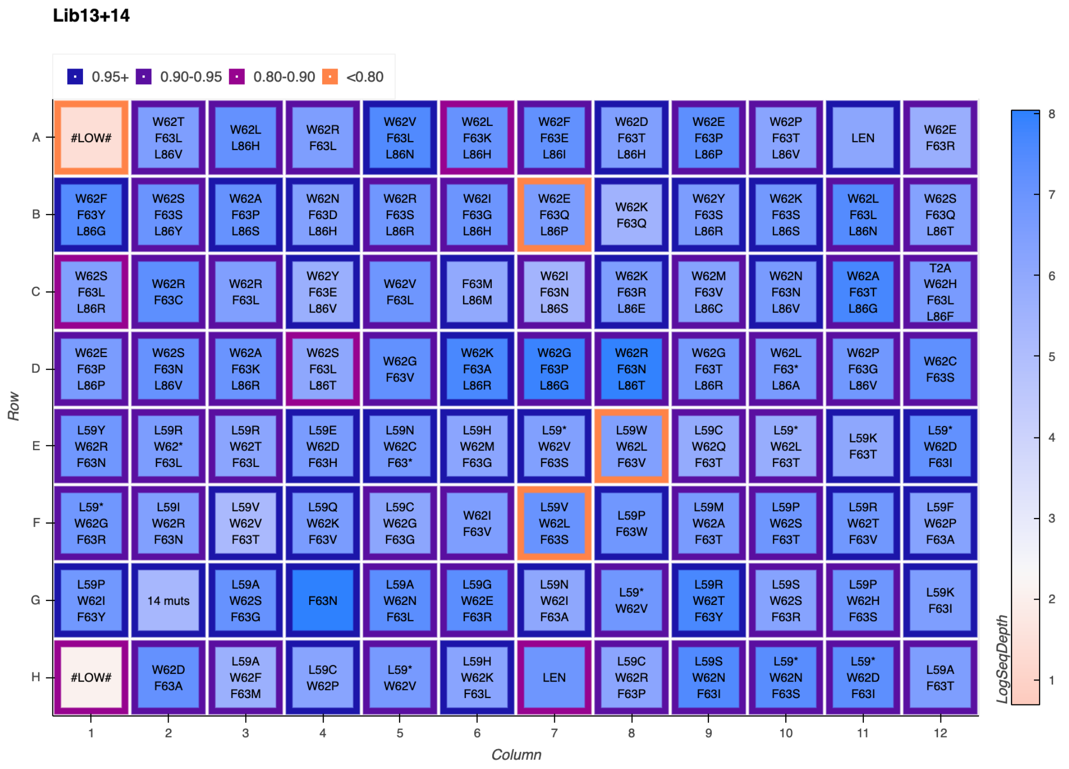


**Figure S15.** LevSeq sequencing plate map for additional random variants picked from libraries 13 and 14.

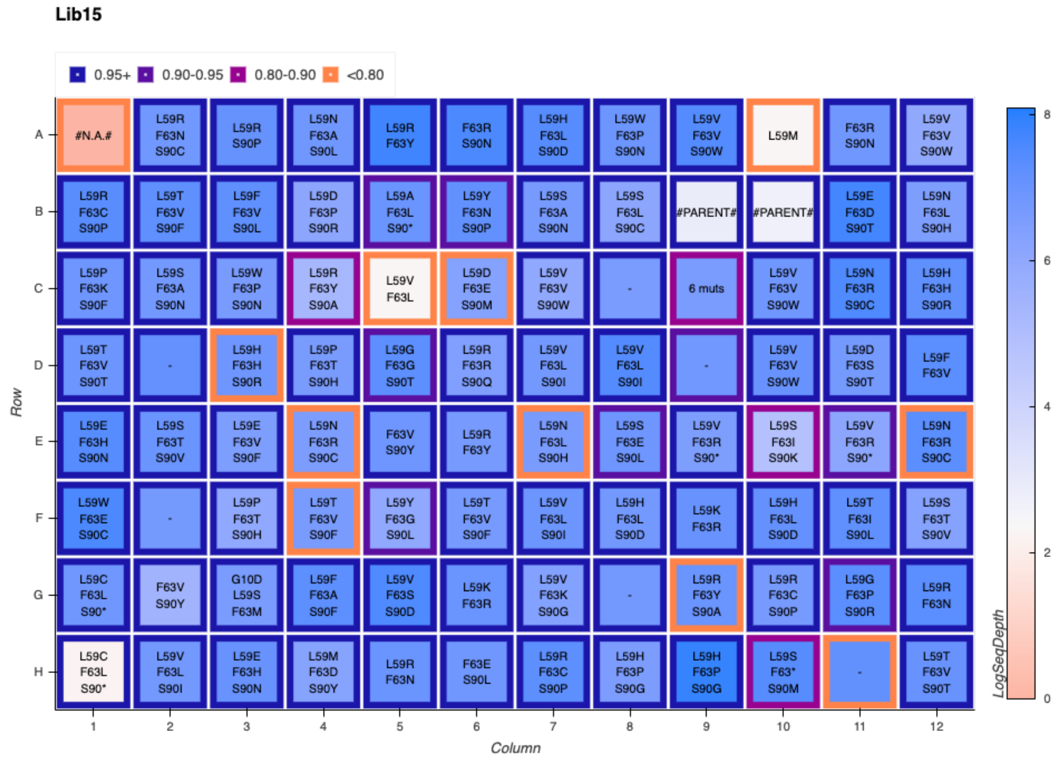


**Figure S16.** LevSeq sequencing plate map for random variants picked from library 15.

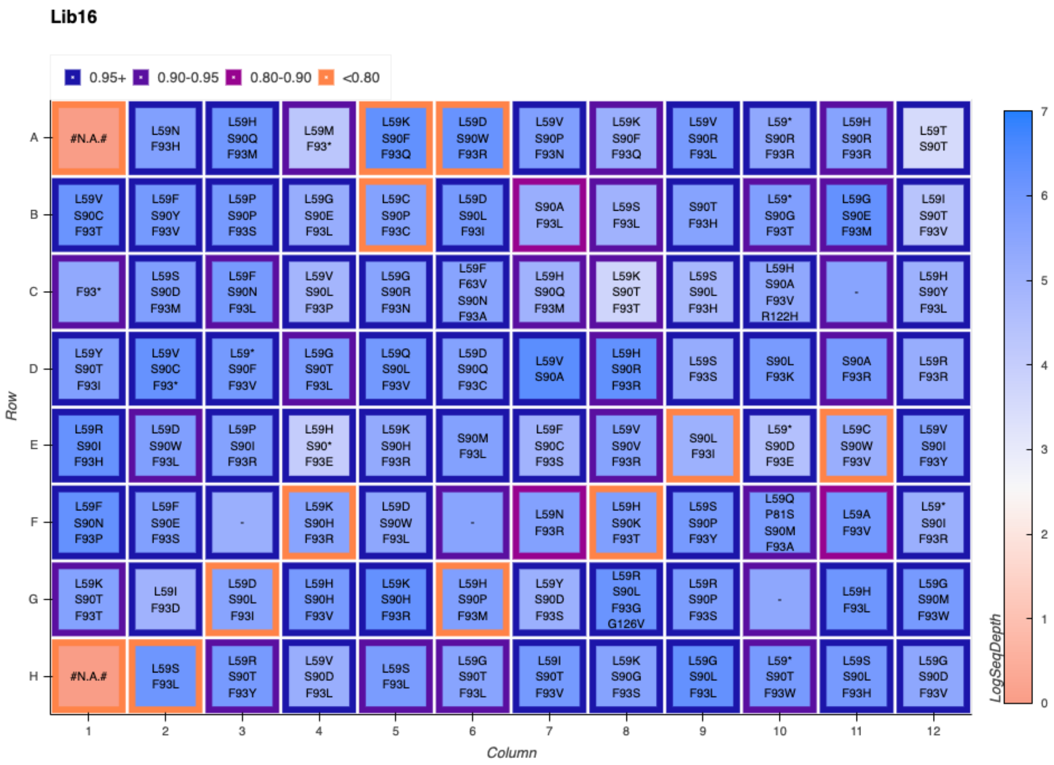


**Figure S17.** LevSeq sequencing plate map for random variants picked from library 16.

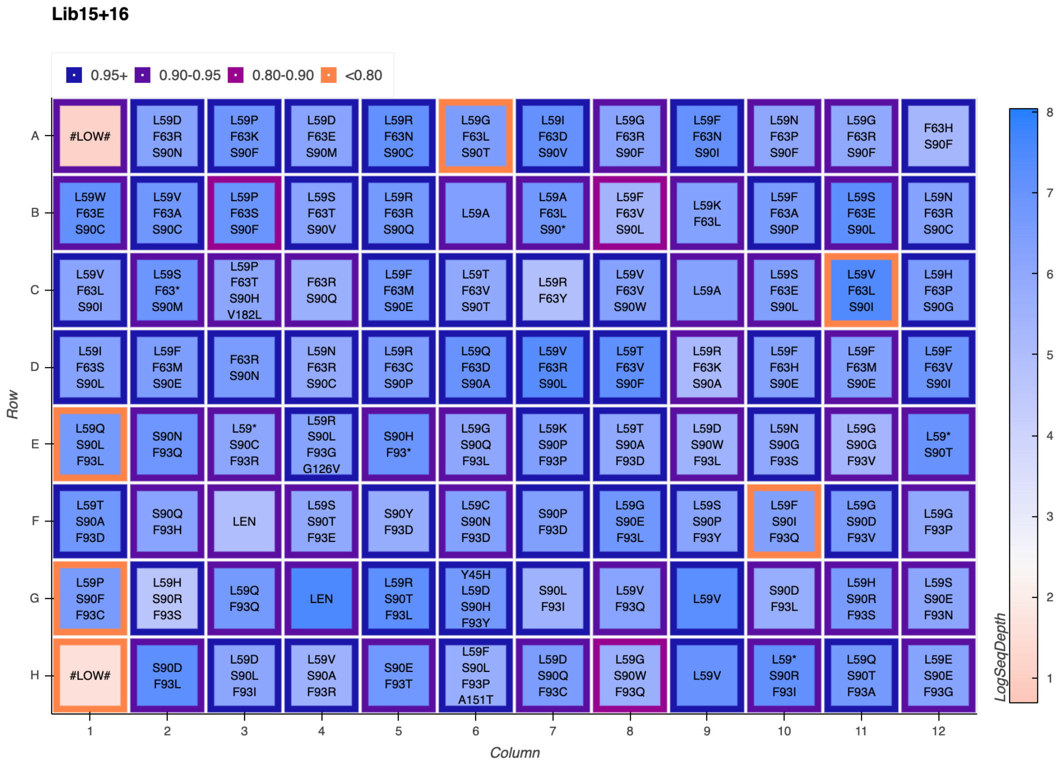


**Figure S18.** LevSeq sequencing plate map for additional random variants picked from libraries 15 and 16.

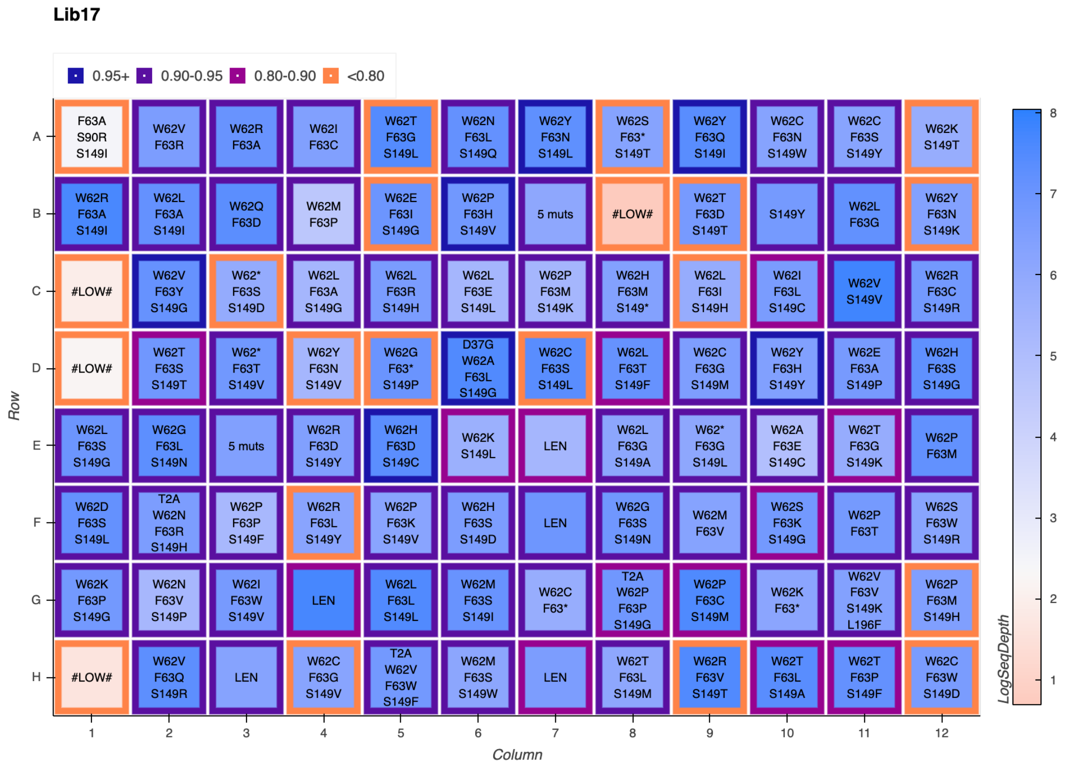


**Figure S19.** LevSeq sequencing plate map for random variants picked from library 17.

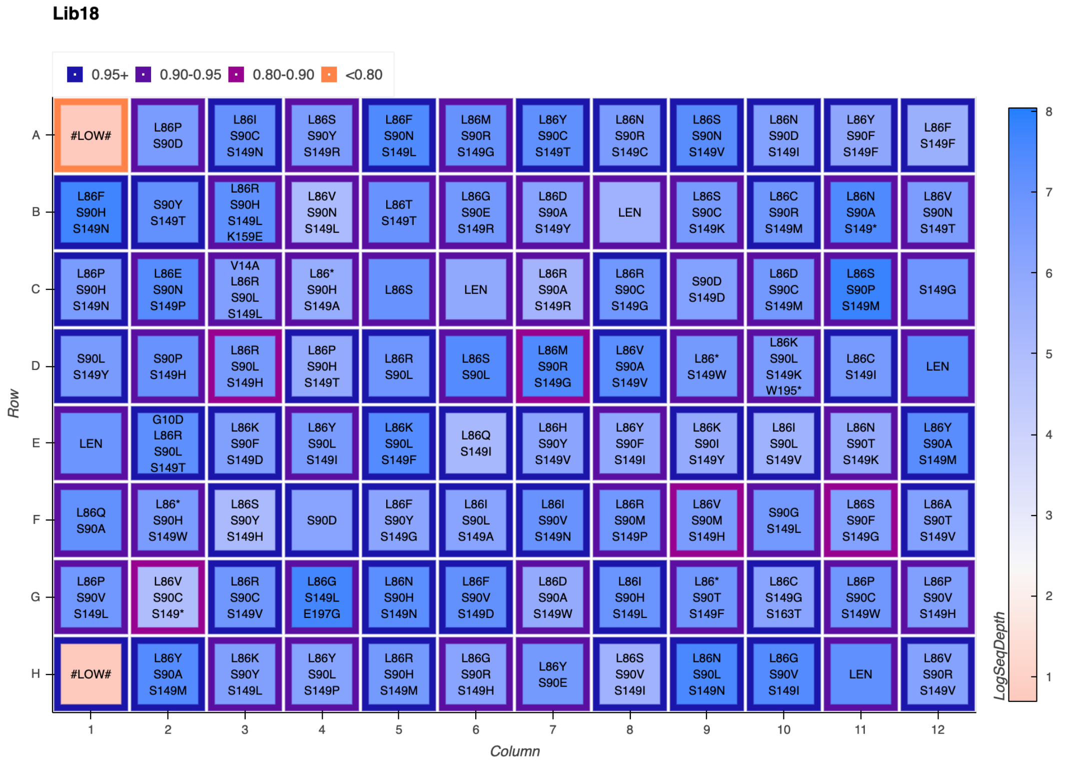


**Figure S20.** LevSeq sequencing plate map for random variants picked from library 18.

*Rearray of Random Mutants*

Multi-mutation variants were rearrayed from the previously described, randomly picked, plates to reduce screening burden for model training data collection. A Python script was used to randomly select 22 variants from each of the 12 double-mutant libraries and 44 variants from each of the six triple-site libraries, yielding 528 unique multi-mutants. These variants were rearrayed over six 96-well deep-well plates in the following manner: The wells of a 2-mL 96-well deep-well plate were filled with 400 µL LB-Amp. Previously generated 96-well plates were removed from -80 °C storage and placed on dry ice. Pipet tips were used to scratch the frozen glycerol stock surface and used to inoculate 88 wells of the aforementioned deep-well plate. Additionally, to each of these deep-well plates, six wells were inoculated with *E. coli* harboring the parent gene, PromPgb, one well was inoculated with *E. coli* harboring a gene encoding a tryptophan synthase variant (TrpB, UniProt: P0A879), and one well was left sterile. These overnight cultures were incubated at 37 ºC and shaken at 220 rpm for 16–18 hours. The following morning, 50 µL of overnight culture from each well were added to the wells of a 96-well flat-bottom tissue culture plate (ThermoFisher) preloaded with 50 µL of 50% glycerol solution. These glycerol stocks were stored at -80°C for future inoculation.

**Table S9.** Variants selected as members of the initial training library for model training. A Python script was used to randomly select sequenced variants generated through multi-NNK cloning.

| Library | Variant |
|---|---|
| 1 | ['L59L', 'W62N'] |
| 1 | ['L59T', 'W62G'] |
| 1 | ['L59A', 'W62T'] |
| 1 | ['L59C', 'W62R'] |
| 1 | ['L59R', 'W62L'] |
| 1 | ['L59G', 'W62D'] |
| 1 | ['L59P', 'W62S'] |

| | |
|---|---|
| 1 | ['L59C', 'W62G'] |
| 1 | ['L59R', 'W62S'] |
| 1 | ['L59P', 'W62K'] |
| 1 | ['L59T', 'W62D'] |
| 1 | ['L59S', 'W62H'] |
| 1 | ['L59N', 'W62R'] |
| 1 | ['L59N', 'W62I'] |
| 1 | ['L59H', 'W62S'] |
| 1 | ['L59P', 'W62G'] |
| 1 | ['L59Q', 'W62H'] |
| 1 | ['L59R', 'W62I'] |
| 1 | ['L59A', 'W62C'] |
| 1 | ['L59F', 'W62M'] |
| 1 | ['L59N', 'W62T'] |
| 1 | ['L59P', 'W62L'] |
| 2 | ['W62P', 'F63R'] |
| 2 | ['W62V', 'F63K'] |
| 2 | ['W62T', 'F63D'] |
| 2 | ['W62L', 'F63L'] |
| 2 | ['W62A', 'F63I'] |
| 2 | ['W62G', 'F63E'] |
| 2 | ['W62F', 'F63C'] |
| 2 | ['W62L', 'F63V'] |
| 2 | ['W62F', 'F63Q'] |
| 2 | ['W62N', 'F63E'] |
| 2 | ['W62I', 'F63H'] |
| 2 | ['W62S', 'F63E'] |
| 2 | ['W62S', 'F63K'] |
| 2 | ['W62R', 'F63W'] |
| 2 | ['W62E', 'F63P'] |
| 2 | ['W62V', 'F63V'] |
| 2 | ['W62V', 'F63A'] |
| 2 | ['W62D', 'F63N'] |

| | |
|---|---|
| 2 | ['W62S', 'F63R'] |
| 3 | ['F63P', 'F73I'] |
| 3 | ['F63R', 'F73M'] |
| 3 | ['F63W', 'F73W'] |
| 3 | ['F63H', 'F73R'] |
| 3 | ['F63Q', 'F73H'] |
| 3 | ['F63Y', 'F73C'] |
| 3 | ['F63N', 'F73S'] |
| 3 | ['F63A', 'F73T'] |
| 3 | ['F63D', 'F73W'] |
| 3 | ['F63P', 'F73T'] |
| 3 | ['F63D', 'F73V'] |
| 3 | ['F63P', 'F73S'] |
| 3 | ['F63T', 'F73Y'] |
| 3 | ['F63D', 'F73S'] |
| 3 | ['F63M', 'F73R'] |
| 3 | ['F63E', 'F73V'] |
| 3 | ['F63T', 'F73C'] |
| 3 | ['F63Y', 'F73P'] |
| 3 | ['F63G', 'F73E'] |
| 3 | ['F63Q', 'F73K'] |
| 3 | ['F63S', 'F73I'] |
| 3 | ['F63L', 'F73Y'] |
| 4 | ['F73G', 'L86P'] |
| 4 | ['F73G', 'L86N'] |
| 4 | ['F73R', 'L86Y'] |
| 4 | ['F73A', 'L86V'] |
| 4 | ['F73V', 'L86A'] |
| 4 | ['F73V', 'L86T'] |
| 4 | ['F73L', 'L86T'] |
| 4 | ['F73P', 'L86T'] |
| 4 | ['F73R', 'L86V'] |
| 4 | ['F73H', 'L86T'] |

| | |
|---|---|
| 4 | ['F73H', 'L86D'] |
| 4 | ['F73K', 'L86S'] |
| 4 | ['F73D', 'L86T'] |
| 4 | ['F73L', 'L86S'] |
| 4 | ['F73N', 'L86I'] |
| 4 | ['F73Y', 'L86K'] |
| 4 | ['F73R', 'L86N'] |
| 4 | ['F73K', 'L86Q'] |
| 4 | ['F73P', 'L86M'] |
| 4 | ['F73D', 'L86A'] |
| 4 | ['F73T', 'L86Y'] |
| 4 | ['F73P', 'L86V'] |
| 5 | ['L86H', 'S90R'] |
| 5 | ['L86I', 'S90L'] |
| 5 | ['L86V', 'S90Y'] |
| 5 | ['L86P', 'S90Y'] |
| 5 | ['L86I', 'S90Y'] |
| 5 | ['L86F', 'S90F'] |
| 5 | ['L86S', 'S90P'] |
| 5 | ['L86K', 'S90L'] |
| 5 | ['L86Q', 'S90T'] |
| 5 | ['L86T', 'S90Y'] |
| 5 | ['L86F', 'S90H'] |
| 5 | ['L86S', 'S90A'] |
| 5 | ['L86C', 'S90H'] |
| 5 | ['L86D', 'S90D'] |
| 5 | ['L86D', 'S90F'] |
| 5 | ['L86C', 'S90C'] |
| 5 | ['L86N', 'S90I'] |
| 5 | ['L86M', 'S90T'] |
| 5 | ['L86F', 'S90D'] |
| 5 | ['L86P', 'S90P'] |
| 5 | ['L86T', 'S90N'] |

| | |
|---|---|
| 5 | ['L86V', 'S90I'] |
| 6 | ['S90L', 'F93D'] |
| 6 | ['S90H', 'F93G'] |
| 6 | ['S90N', 'F93T'] |
| 6 | ['S90H', 'F93R'] |
| 6 | ['S90A', 'F93R'] |
| 6 | ['S90E', 'F93R'] |
| 6 | ['S90H', 'F93P'] |
| 6 | ['S90Q', 'F93S'] |
| 6 | ['S90L', 'F93I'] |
| 6 | ['S90Q', 'F93G'] |
| 6 | ['S90R', 'F93E'] |
| 6 | ['S90C', 'F93L'] |
| 6 | ['S90H', 'F93Y'] |
| 6 | ['S90N', 'F93L'] |
| 6 | ['S90V', 'F93L'] |
| 6 | ['S90L', 'F93W'] |
| 6 | ['S90D', 'F93H'] |
| 6 | ['S90N', 'F93V'] |
| 6 | ['S90F', 'F93R'] |
| 6 | ['S90H', 'F93M'] |
| 6 | ['S90W', 'F93G'] |
| 6 | ['S90P', 'F93I'] |
| 7 | ['F93K', 'S149K'] |
| 7 | ['F93E', 'S149H'] |
| 7 | ['F93T', 'S149T'] |
| 7 | ['F93Q', 'S149V'] |
| 7 | ['F93P', 'S149T'] |
| 7 | ['F93R', 'S149P'] |
| 7 | ['F93V', 'S149Y'] |
| 7 | ['F93C', 'S149H'] |
| 7 | ['F93T', 'S149G'] |
| 7 | ['F93S', 'S149F'] |

| | |
|---|---|
| 7 | ['F93Y', 'S149R'] |
| 7 | ['F93S', 'S149N'] |
| 7 | ['F93W', 'S149I'] |
| 7 | ['F93L', 'S149A'] |
| 7 | ['F93R', 'S149T'] |
| 7 | ['F93Y', 'S149L'] |
| 7 | ['F93D', 'S149G'] |
| 7 | ['F93S', 'S149L'] |
| 7 | ['F93H', 'S149I'] |
| 7 | ['F93T', 'S149Q'] |
| 7 | ['F93M', 'S149V'] |
| 7 | ['F93P', 'S149I'] |
| 8 | ['L59T', 'S149G'] |
| 8 | ['L59Q', 'S149G'] |
| 8 | ['L59V', 'S149W'] |
| 8 | ['L59F', 'S149T'] |
| 8 | ['L59H', 'S149M'] |
| 8 | ['L59K', 'S149L'] |
| 8 | ['L59Y', 'S149K'] |
| 8 | ['L59S', 'S149W'] |
| 8 | ['L59N', 'S149R'] |
| 8 | ['L59T', 'S149L'] |
| 8 | ['L59R', 'S149M'] |
| 8 | ['L59G', 'S149R'] |
| 8 | ['L59Q', 'S149R'] |
| 8 | ['L59T', 'S149H'] |
| 8 | ['L59I', 'S149Q'] |
| 8 | ['L59Q', 'S149H'] |
| 8 | ['L59R', 'S149V'] |
| 8 | ['L59R', 'S149R'] |
| 8 | ['L59W', 'S149V'] |
| 8 | ['L59C', 'S149G'] |
| 8 | ['L59H', 'S149V'] |

| | |
|---|---|
| 8 | ['L59V', 'S149I'] |
| 9 | ['L59M', 'F73R'] |
| 9 | ['L59D', 'F73Y'] |
| 9 | ['L59V', 'F73H'] |
| 9 | ['L59R', 'F73L'] |
| 9 | ['L59G', 'F73K'] |
| 9 | ['L59S', 'F73E'] |
| 9 | ['L59F', 'F73N'] |
| 9 | ['L59S', 'F73V'] |
| 9 | ['L59R', 'F73R'] |
| 9 | ['L59S', 'F73S'] |
| 9 | ['L59P', 'F73A'] |
| 9 | ['L59C', 'F73M'] |
| 9 | ['L59I', 'F73A'] |
| 9 | ['L59H', 'F73N'] |
| 9 | ['L59A', 'F73R'] |
| 9 | ['L59Y', 'F73E'] |
| 9 | ['L59S', 'F73Q'] |
| 9 | ['L59G', 'F73L'] |
| 9 | ['L59T', 'F73H'] |
| 9 | ['L59P', 'F73L'] |
| 9 | ['L59T', 'F73P'] |
| 9 | ['L59Y', 'F73K'] |
| 10 | ['F73L', 'S90M'] |
| 10 | ['F73S', 'S90R'] |
| 10 | ['F73S', 'S90F'] |
| 10 | ['F73T', 'S90T'] |
| 10 | ['F73S', 'S90L'] |
| 10 | ['F73W', 'S90G'] |
| 10 | ['F73L', 'S90Y'] |
| 10 | ['F73V', 'S90G'] |
| 10 | ['F73T', 'S90R'] |
| 10 | ['F73K', 'S90I'] |

| | |
|---|---|
| 10 | ['F73A', 'S90W'] |
| 10 | ['F73Y', 'S90R'] |
| 10 | ['F73M', 'S90Q'] |
| 10 | ['F73N', 'S90C'] |
| 10 | ['F73T', 'S90V'] |
| 10 | ['F73T', 'S90Y'] |
| 10 | ['F73L', 'S90K'] |
| 10 | ['F73V', 'S90P'] |
| 10 | ['F73R', 'S90T'] |
| 10 | ['F73L', 'S90P'] |
| 10 | ['F73Y', 'S90L'] |
| 10 | ['F73N', 'S90A'] |
| 11 | ['L86S', 'F93V'] |
| 11 | ['L86K', 'F93V'] |
| 11 | ['L86K', 'F93C'] |
| 11 | ['L86R', 'F93T'] |
| 11 | ['L86S', 'F93T'] |
| 11 | ['L86P', 'F93Y'] |
| 11 | ['L86S', 'F93W'] |
| 11 | ['L86I', 'F93W'] |
| 11 | ['L86S', 'F93H'] |
| 11 | ['L86A', 'F93R'] |
| 11 | ['L86D', 'F93C'] |
| 11 | ['L86A', 'F93Y'] |
| 11 | ['L86V', 'F93G'] |
| 11 | ['L86A', 'F93I'] |
| 11 | ['L86N', 'F93S'] |
| 11 | ['L86R', 'F93D'] |
| 11 | ['L86A', 'F93M'] |
| 11 | ['L86N', 'F93A'] |
| 11 | ['L86E', 'F93G'] |
| 11 | ['L86N', 'F93L'] |
| 11 | ['L86N', 'F93G'] |

| | |
|---|---|
| 11 | ['L86C', 'F93V'] |
| 12 | ['L59R', 'F93L'] |
| 12 | ['L59S', 'F93M'] |
| 12 | ['L59T', 'F93Y'] |
| 12 | ['L59Q', 'F93G'] |
| 12 | ['L59Y', 'F93G'] |
| 12 | ['L59W', 'F93C'] |
| 12 | ['L59M', 'F93L'] |
| 12 | ['L59S', 'F93D'] |
| 12 | ['L59E', 'F93A'] |
| 12 | ['L59C', 'F93Y'] |
| 12 | ['L59V', 'F93L'] |
| 12 | ['L59Q', 'F93K'] |
| 12 | ['L59V', 'F93P'] |
| 12 | ['L59E', 'F93G'] |
| 12 | ['L59C', 'F93I'] |
| 12 | ['L59I', 'F93V'] |
| 12 | ['L59R', 'F93Y'] |
| 12 | ['L59V', 'F93K'] |
| 12 | ['L59S', 'F93C'] |
| 12 | ['L59H', 'F93P'] |
| 12 | ['L59T', 'F93W'] |
| 12 | ['L59A', 'F93S'] |
| 13 | ['W62D', 'F63G', 'L86H'] |
| 13 | ['W62R', 'F63Y', 'L86N'] |
| 13 | ['W62H', 'F63R', 'L86V'] |
| 13 | ['W62L', 'F63G', 'L86H'] |
| 13 | ['W62V', 'F63S', 'L86F'] |

| | |
|---|---|
| 13 | ['W62Q', 'F63K', 'L86G'] |
| 13 | ['W62V', 'F63K', 'L86M'] |
| 13 | ['W62A', 'F63I', 'L86A'] |
| 13 | ['W62S', 'F63S', 'L86K'] |
| 13 | ['W62S', 'F63V', 'L86T'] |
| 13 | ['W62L', 'F63D', 'L86N'] |
| 13 | ['W62P', 'F63M', 'L86G'] |
| 13 | ['W62H', 'F63P', 'L86V'] |
| 13 | ['W62G', 'F63H', 'L86I'] |
| 13 | ['W62A', 'F63S', 'L86D'] |
| 13 | ['W62P', 'F63L', 'L86R'] |
| 13 | ['W62R', 'F63V', 'L86N'] |
| 13 | ['W62S', 'F63A', 'L86S'] |
| 13 | ['W62P', 'F63V', 'L86R'] |
| 13 | ['W62E', 'F63N', 'L86I'] |
| 13 | ['W62P', 'F63W', 'L86Q'] |

| | |
|---|---|
| 13 | ['W62F', 'F63A', 'L86D'] |
| 13 | ['W62R', 'F63L', 'L86R'] |
| 13 | ['W62L', 'F63G', 'L86K'] |
| 13 | ['W62N', 'F63M', 'L86V'] |
| 13 | ['W62R', 'F63C', 'L86F'] |
| 13 | ['W62Q', 'F63G', 'L86S'] |
| 13 | ['W62L', 'F63H', 'L86G'] |
| 13 | ['W62T', 'F63T', 'L86D'] |
| 13 | ['W62R', 'F63S', 'L86V'] |
| 13 | ['W62A', 'F63D', 'L86H'] |
| 13 | ['W62S', 'F63R', 'L86V'] |
| 13 | ['W62F', 'F63R', 'L86H'] |
| 13 | ['W62I', 'F63R', 'L86K'] |
| 13 | ['W62V', 'F63L', 'L86G'] |
| 13 | ['W62C', 'F63V', 'L86W'] |
| 13 | ['W62F', 'F63Y', 'L86G'] |

| | |
|---|---|
| 13 | ['W62K', 'F63R', 'L86E'] |
| 13 | ['W62Y', 'F63E', 'L86V'] |
| 13 | ['W62K', 'F63S', 'L86S'] |
| 13 | ['W62A', 'F63T','L86G'] |
| 13 | ['W62A', 'F63P', 'L86S'] |
| 13 | ['W62S', 'F63N', 'L86V'] |
| 13 | ['W62S', 'F63S', 'L86Y'] |
| 14 | ['L59S', 'W62C', 'F63I'] |
| 14 | ['L59V', 'W62I', 'F63S'] |
| 14 | ['L59S', 'W62V', 'F63S'] |
| 14 | ['L59K', 'W62V', 'F63S'] |
| 14 | ['L59P', 'W62C', 'F63N'] |
| 14 | ['L59C', 'W62Q', 'F63A'] |
| 14 | ['L59N', 'W62R', 'F63E'] |
| 14 | ['L59T', 'W62F', 'F63A'] |
| 14 | ['L59H', 'W62D', 'F63S'] |
| 14 | ['L59C', 'W62D', 'F63K'] |

| | |
|---|---|
| 14 | ['L59N', 'W62R', 'F63S'] |
| 14 | ['L59G', 'W62R', 'F63T'] |
| 14 | ['L59R', 'W62E', 'F63V'] |
| 14 | ['L59W', 'W62G', 'F63G'] |
| 14 | ['L59H', 'W62F', 'F63L'] |
| 14 | ['L59Y', 'W62M', 'F63N'] |
| 14 | ['L59Q', 'W62Y', 'F63L'] |
| 14 | ['L59T', 'W62K', 'F63R'] |
| 14 | ['L59P', 'W62R', 'F63T'] |
| 14 | ['L59M', 'W62N', 'F63G'] |
| 14 | ['L59T', 'W62G', 'F63M'] |
| 14 | ['L59V', 'W62A', 'F63R'] |
| 14 | ['L59P', 'W62I', 'F63Q'] |
| 14 | ['L59R', 'W62L', 'F63K'] |
| 14 | ['L59R', 'W62Y', 'F63E'] |
| 14 | ['L59N', 'W62L', 'F63E'] |
| 14 | ['L59W', 'W62E', 'F63G'] |

| | |
|---|---|
| 14 | ['L59E', 'W62R', 'F63L'] |
| 14 | ['L59M', 'W62V', 'F63V'] |
| 14 | ['L59A', 'W62L', 'F63V'] |
| 14 | ['L59F', 'W62V', 'F63A'] |
| 14 | ['L59V', 'W62S', 'F63S'] |
| 14 | ['L59S', 'W62A', 'F63C'] |
| 14 | ['L59K', 'W62V', 'F63G'] |
| 14 | ['L59R', 'W62P', 'F63L'] |
| 14 | ['L59P', 'W62R', 'F63N'] |
| 14 | ['L59I', 'W62R', 'F63N'] |
| 14 | ['L59Q', 'W62K', 'F63V'] |
| 14 | ['L59A', 'W62S', 'F63G'] |
| 14 | ['L59P', 'W62H', 'F63S'] |
| 14 | ['L59F', 'W62P', 'F63A'] |
| 14 | ['L59H', 'W62M', 'F63G'] |
| 14 | ['L59C', 'W62Q', 'F63T'] |

| | |
|---|---|
| 14 | ['L59C', 'W62G', 'F63G'] |
| 15 | ['L59R', 'F63N', 'S90C'] |
| 15 | ['L59P', 'F63K', 'S90F'] |
| 15 | ['L59H', 'F63P', 'S90G'] |
| 15 | ['L59T', 'F63I', 'S90L'] |
| 15 | ['L59Y', 'F63G', 'S90L'] |
| 15 | ['L59S', 'F63L', 'S90C'] |
| 15 | ['L59Y', 'F63N', 'S90P'] |
| 15 | ['L59T', 'F63V', 'S90T'] |
| 15 | ['L59S', 'F63T', 'S90V'] |
| 15 | ['L59N', 'F63R', 'S90C'] |
| 15 | ['L59N', 'F63L', 'S90H'] |
| 15 | ['L59F', 'F63A', 'S90F'] |
| 15 | ['L59F', 'F63V', 'S90L'] |
| 15 | ['L59E', 'F63H', 'S90N'] |
| 15 | ['L59D', 'F63S', 'S90T'] |
| 15 | ['L59N', 'F63A', 'S90L'] |
| 15 | ['L59P', 'F63T', 'S90H'] |
| 15 | ['L59G', 'F63G', 'S90T'] |
| 15 | ['L59V', 'F63V', 'S90W'] |
| 15 | ['L59S', 'F63A', 'S90N'] |
| 15 | ['L59W', 'F63P', 'S90N'] |
| 15 | ['L59G', 'F63P', 'S90R'] |

| | |
|---|---|
| 15 | ['L59V', 'F63K', 'S90G'] |
| 15 | ['L59D', 'F63P', 'S90R'] |
| 15 | ['L59E', 'F63D', 'S90T'] |
| 15 | ['L59R', 'F63R', 'S90Q'] |
| 15 | ['L59V', 'F63L', 'S90I'] |
| 15 | ['L59V', 'F63S', 'S90D'] |
| 15 | ['L59T', 'F63V', 'S90F'] |
| 15 | ['L59M', 'F63D', 'S90Y'] |
| 15 | ['L59R', 'F63C', 'S90P'] |
| 15 | ['L59S', 'F63E', 'S90L'] |
| 15 | ['L59H', 'F63L', 'S90D'] |
| 15 | ['L59H', 'F63H', 'S90R'] |
| 15 | ['L59E', 'F63V', 'S90F'] |
| 15 | ['L59W', 'F63E', 'S90C'] |
| 15 | ['L59V', 'F63A', 'S90C'] |
| 15 | ['L59F', 'F63M', 'S90E'] |
| 15 | ['L59F', 'F63A', 'S90P'] |
| 15 | ['L59I', 'F63S', 'S90L'] |
| 15 | ['L59Q', 'F63D', 'S90A'] |
| 15 | ['L59G', 'F63R', 'S90F'] |
| 15 | ['L59I', 'F63D', 'S90V'] |
| 15 | ['L59D', 'F63E', 'S90M'] |
| 16 | ['L59V', 'S90L', 'F93P'] |

| | |
|---|---|
| 16 | ['L59H', 'S90Y', 'F93L'] |
| 16 | ['L59K', 'S90G', 'F93S'] |
| 16 | ['L59P', 'S90P', 'F93S'] |
| 16 | ['L59Y', 'S90T', 'F93I'] |
| 16 | ['L59K', 'S90H', 'F93R'] |
| 16 | ['L59R', 'S90T', 'F93Y'] |
| 16 | ['L59V', 'S90P', 'F93N'] |
| 16 | ['L59I', 'S90T', 'F93V'] |
| 16 | ['L59G', 'S90E', 'F93L'] |
| 16 | ['L59Q', 'S90L', 'F93V'] |
| 16 | ['L59F', 'S90N', 'F93L'] |
| 16 | ['L59R', 'S90P', 'F93S'] |
| 16 | ['L59R', 'S90I', 'F93H'] |
| 16 | ['L59D', 'S90L', 'F93I'] |
| 16 | ['L59F', 'S90N', 'F93P'] |
| 16 | ['L59G', 'S90R', 'F93N'] |
| 16 | ['L59S', 'S90D', 'F93M'] |
| 16 | ['L59V', 'S90I', 'F93Y'] |
| 16 | ['L59D', 'S90Q', 'F93C'] |
| 16 | ['L59H', 'S90R', 'F93R'] |
| 16 | ['L59F', 'S90Y', 'F93V'] |
| 16 | ['L59S', 'S90P', 'F93Y'] |
| 16 | ['L59G', 'S90M', 'F93W'] |
| 16 | ['L59G', 'S90E', 'F93M'] |

| | |
|---|---|
| 16 | ['L59P', 'S90I', 'F93R'] |
| 16 | ['L59D', 'S90W', 'F93L'] |
| 16 | ['L59G', 'S90T', 'F93L'] |
| 16 | ['L59K', 'S90F', 'F93Q'] |
| 16 | ['L59H', 'S90Q', 'F93M'] |
| 16 | ['L59H', 'S90H', 'F93V'] |
| 16 | ['L59F', 'S90E', 'F93S'] |
| 16 | ['L59G', 'S90L', 'F93L'] |
| 16 | ['L59V', 'S90D', 'F93L'] |
| 16 | ['L59S', 'S90L', 'F93H'] |
| 16 | ['L59V', 'S90R', 'F93L'] |
| 16 | ['L59K', 'S90P', 'F93P'] |
| 16 | ['L59G', 'S90Q', 'F93L'] |
| 16 | ['L59Q', 'S90T', 'F93A'] |
| 16 | ['L59H', 'S90R', 'F93S'] |
| 16 | ['L59G', 'S90G', 'F93V'] |
| 16 | ['L59T', 'S90A', 'F93D'] |
| 16 | ['L59C', 'S90N', 'F93D'] |
| 16 | ['L59N', 'S90G', 'F93S'] |
| 17 | ['W62C', 'F63G', 'S149M'] |
| 17 | ['W62N', 'F63L', 'S149Q'] |

| | |
|---|---|
| 17 | ['W62L', 'F63E', 'S149L'] |
| 17 | ['W62I', 'F63L', 'S149C'] |
| 17 | ['W62S', 'F63K', 'S149G'] |
| 17 | ['W62T', 'F63G', 'S149K'] |
| 17 | ['W62P', 'F63C', 'S149M'] |
| 17 | ['W62M', 'F63S', 'S149W'] |
| 17 | ['W62L', 'F63S', 'S149G'] |
| 17 | ['W62L', 'F63R', 'S149H'] |
| 17 | ['W62P', 'F63K', 'S149V'] |
| 17 | ['W62H', 'F63D', 'S149C'] |
| 17 | ['W62P', 'F63H', 'S149V'] |
| 17 | ['W62G', 'F63L', 'S149N'] |
| 17 | ['W62L', 'F63T', 'S149F'] |
| 17 | ['W62R', 'F63D', 'S149Y'] |
| 17 | ['W62N', 'F63V', 'S149P'] |
| 17 | ['W62H', 'F63S', 'S149D'] |

| | |
|---|---|
| 17 | ['W62G', 'F63S', 'S149N'] |
| 17 | ['W62P', 'F63P', 'S149F'] |
| 17 | ['W62Y', 'F63H', 'S149Y'] |
| 17 | ['W62E', 'F63A', 'S149P'] |
| 17 | ['W62C', 'F63N', 'S149W'] |
| 17 | ['W62T', 'F63P', 'S149F'] |
| 17 | ['W62L', 'F63L', 'S149L'] |
| 17 | ['W62L', 'F63A', 'S149I'] |
| 17 | ['W62T', 'F63L', 'S149M'] |
| 17 | ['W62L', 'F63A', 'S149G'] |
| 17 | ['W62H', 'F63S', 'S149G'] |
| 17 | ['W62R', 'F63C', 'S149R'] |
| 17 | ['W62I', 'F63W', 'S149V'] |
| 17 | ['W62V', 'F63Q', 'S149R'] |
| 17 | ['W62C', 'F63S', 'S149Y'] |
| 17 | ['W62T', 'F63S', 'S149T'] |

| | |
|---|---|
| 17 | ['W62M', 'F63S', 'S149I'] |
| 17 | ['W62K', 'F63P', 'S149G'] |
| 17 | ['W62R', 'F63A', 'S149I'] |
| 17 | ['W62V', 'F63Y', 'S149G'] |
| 17 | ['W62D', 'F63S', 'S149L'] |
| 17 | ['W62Y', 'F63N', 'S149L'] |
| 17 | ['W62A', 'F63E', 'S149C'] |
| 17 | ['W62S', 'F63W', 'S149R'] |
| 17 | ['W62L', 'F63G', 'S149A'] |
| 17 | ['W62P', 'F63M', 'S149K'] |
| 18 | ['L86V', 'S90N', 'S149T'] |
| 18 | ['L86E', 'S90N', 'S149P'] |
| 18 | ['L86S', 'S90Y', 'S149H'] |
| 18 | ['L86D', 'S90A', 'S149Y'] |
| 18 | ['L86S', 'S90C', 'S149K'] |
| 18 | ['L86V', 'S90M', 'S149H'] |

| | |
|---|---|
| 18 | ['L86K', 'S90I', 'S149Y'] |
| 18 | ['L86I', 'S90V', 'S149N'] |
| 18 | ['L86V', 'S90R', 'S149V'] |
| 18 | ['L86F', 'S90H', 'S149N'] |
| 18 | ['L86P', 'S90H', 'S149N'] |
| 18 | ['L86G', 'S90R', 'S149H'] |
| 18 | ['L86R', 'S90A', 'S149R'] |
| 18 | ['L86G', 'S90V', 'S149I'] |
| 18 | ['L86K', 'S90F', 'S149D'] |
| 18 | ['L86I', 'S90L', 'S149V'] |
| 18 | ['L86R', 'S90H', 'S149M'] |
| 18 | ['L86V', 'S90A', 'S149V'] |
| 18 | ['L86P', 'S90V', 'S149H'] |
| 18 | ['L86I', 'S90C', 'S149N'] |
| 18 | ['L86S', 'S90P', 'S149M'] |
| 18 | ['L86Y', 'S90A', 'S149M'] |

| | |
|---|---|
| 18 | ['L86M', 'S90R', 'S149G'] |
| 18 | ['L86Y', 'S90L', 'S149I'] |
| 18 | ['L86F', 'S90V', 'S149D'] |
| 18 | ['L86Y', 'S90F', 'S149F'] |
| 18 | ['L86C', 'S90R', 'S149M'] |
| 18 | ['L86R', 'S90C', 'S149G'] |
| 18 | ['L86N', 'S90T', 'S149K'] |
| 18 | ['L86R', 'S90M', 'S149P'] |
| 18 | ['L86Y', 'S90C', 'S149T'] |
| 18 | ['L86A', 'S90T', 'S149V'] |
| 18 | ['L86I', 'S90H', 'S149L'] |
| 18 | ['L86S', 'S90N', 'S149V'] |
| 18 | ['L86I', 'S90L', 'S149A'] |
| 18 | ['L86G', 'S90E', 'S149R'] |
| 18 | ['L86P', 'S90H', 'S149T'] |
| 18 | ['L86R', 'S90C', 'S149V'] |

| | |
|---|---|
| 18 | ['L86V', 'S90N', 'S149L'] |
| 18 | ['L86S', 'S90V', 'S149I'] |
| 18 | ['L86D', 'S90A', 'S149W'] |
| 18 | ['L86Y', 'S90L', 'S149P'] |
| 18 | ['L86S', 'S90Y', 'S149R'] |
| 18 | ['L86P', 'S90C', 'S149W'] |

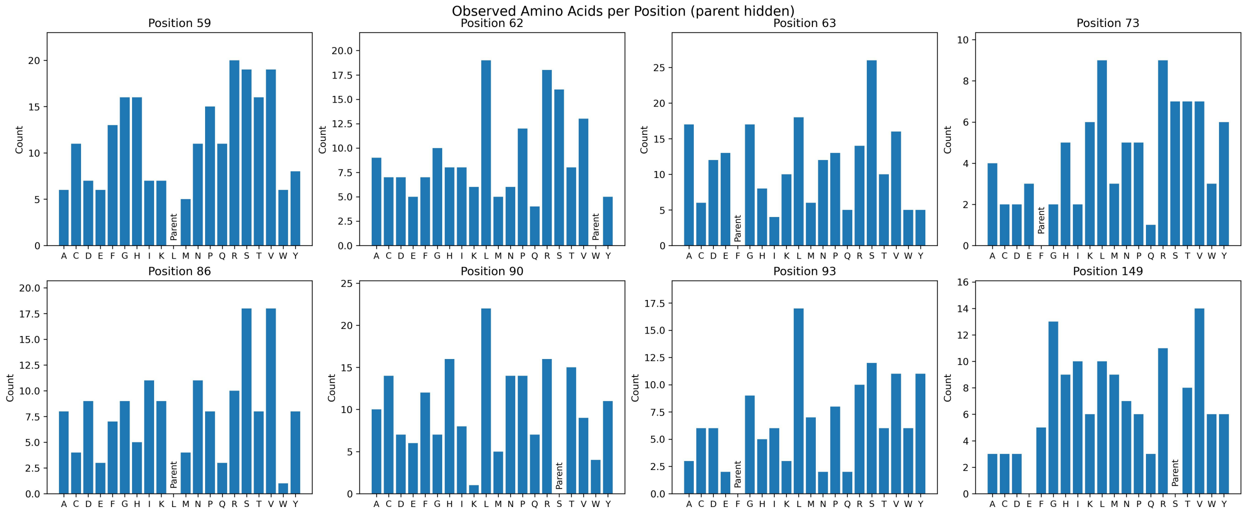


**Figure S21.** Counts of amino acid identities observed at each position in variants selected for the initial training library. The only amino-acid substitution not observed in training was S149E.

#### Protocols for the Cloning of MLDE Predicted Sequences:

##### *Assembly of Plasmids Bearing Predicted Mutant Genes*

Ninety-six multi-mutation variants were generated at each round of predictions. For each round, the DNA sequences for these mutants were achieved through mutagenesis of the target codons in the parent DNA sequence, and an oligo sequence (300 bp) was generated for each variant, containing mutations at up to all eight positions of interest (**Table S2**). All 96 oligo sequences were synthesized and delivered by Twist Bioscience (South San Francisco, CA) as a single pooled sample for each round. Oligo pools were received as dry residues which were reconstituted in 10 mM Tris–HCl (pH=8.0) to a final oligo concentration of 10 ng/µL. Oligo pools were then amplified by PCR using the primers described in **Table S6**. This fragment library was assembled with a pET-22b(+) backbone with overhangs designed for Gibson ligation to generate fully encoded protoglobin sequences bearing exact sets of desired mutations. The assembly products obtained were used to transform T7 Express Competent *E. coli* (High Efficency) cells. Upon heat-shock, freshly transformed *E. coli* cells were recovered in 0.4 mL Luria-Bertani medium (LB) (Research Products Int.) at 37 ºC with shaking at 220 rpm for 30 minutes. This transformation mixture was directly plated on LB-Amp agar plates. The plates were incubated overnight at 37 °C until colony formation was observed. For each round of predictions between 350–550 colonies from LB-Amp agar plates were picked with sterilized toothpicks to individually inoculate the wells of 2-mL 96-well deep-well plates charged with 400 µL of LB-Amp. The plates were incubated at 37 ºC and shaken at 220 rpm for 16–18 hours. The following morning, 50 µL of preculture from each well were added to the wells of a 96-well flat-bottom tissue culture plate (ThermoFisher) preloaded with 50 µL of 50% glycerol solution. These glycerol stocks were stored at -80 °C for future inoculation. Additionally, the sequences of protoglobin genes contained in every well were sequenced using LevSeq sequencing.[5] In total, 63/96 predicted variants were assembled for the first round of oligo pool assembly, and 64/96 were found for achieved in the second round (**Table S10**). The target variants from each round were rearrayed into the wells of a 96-well plate, along with 4-5 parent wells (PromPgb), one TrpB well, and one sterile well. These plates were stored as glycerol stocks at -80 °C for future inoculation.

**Table S10**. PromPgb mutants which were predicted using MLDE to be able to access a broader range of new-to-nature reactions and were successfully cloned using oligo pools.

| **Round** | **Mutant** |
|---|---|
| Predictions Round 1 | CYVFMIFQ |
| Predictions Round 1 | YAIFIIFQ |
| Predictions Round 1 | YALFIVFQ |
| Predictions Round 1 | YYIFMIFQ |
| Predictions Round 1 | YYLFIIFQ |
| Predictions Round 1 | TAVLMGKQ |
| Predictions Round 1 | YYVFMCKQ |
| Predictions Round 1 | QYVFMCFQ |
| Predictions Round 1 | LYLFICKQ |
| Predictions Round 1 | VALFMCVQ |
| Predictions Round 1 | FNIFMAFQ |
| Predictions Round 1 | FYFFMCFQ |
| Predictions Round 1 | FYIFMMFQ |
| Predictions Round 1 | LYIFIMKQ |
| Predictions Round 1 | YAIFMIFQ |
| Predictions Round 1 | FNIFMAKQ |
| Predictions Round 1 | FVIFMVFQ |
| Predictions Round 1 | YAVAIIKQ |
| Predictions Round 1 | TDVFITFQ |
| Predictions Round 1 | FYIFKMKQ |
| Predictions Round 1 | YAVAIVKQ |
| Predictions Round 1 | YAVFIVFQ |
| Predictions Round 1 | VYIAIVFQ |
| Predictions Round 1 | LYLFMMKQ |
| Predictions Round 1 | YYIAIVFQ |
| Predictions Round 1 | FYVFMMFQ |
| Predictions Round 1 | LYIFIVKQ |
| Predictions Round 1 | YYIAMIFQ |
| Predictions Round 1 | YAVVMTFQ |
| Predictions Round 1 | YAIFIVFQ |
| Predictions Round 1 | YYVAICFQ |
| Predictions Round 1 | VYIFMVFQ |
| Predictions Round 1 | YYIFIMFQ |
| Predictions Round 1 | YAVFMTFQ |
| Predictions Round 1 | YYLFICFQ |
| Predictions Round 1 | YYVFMVKQ |

| | |
|---|---|
| Predictions Round 1 | FYVFITFQ |
| Predictions Round 1 | IYVAIIKQ |
| Predictions Round 1 | LYLFIIKQ |
| Predictions Round 1 | LYLFIVKQ |
| Predictions Round 1 | YYVFMIFQ |
| Predictions Round 1 | LYVFIVFQ |
| Predictions Round 1 | VALFMAFQ |
| Predictions Round 1 | TAVLMIKQ |
| Predictions Round 1 | TNVFITFQ |
| Predictions Round 1 | TYVFICVQ |
| Predictions Round 1 | TYVFITFQ |
| Predictions Round 1 | YYVAICKQ |
| Predictions Round 1 | YYVFIVFQ |
| Predictions Round 1 | YYVAMIFQ |
| Predictions Round 1 | YYVAIVKQ |
| Predictions Round 1 | YYVFICFQ |
| Predictions Round 1 | VALAMCVQ |
| Predictions Round 1 | YAIFIIFA |
| Predictions Round 1 | TYVFMIFQ |
| Predictions Round 1 | TYVFMCVQ |
| Predictions Round 1 | TYVFMCFQ |
| Predictions Round 1 | TYVFICFQ |
| Predictions Round 1 | TDVFTCFQ |
| Predictions Round 1 | TAVFMCVQ |
| Predictions Round 1 | LYVAIVFQ |
| Predictions Round 1 | YALFMCVQ |
| Predictions Round 1 | YYVLMIFQ |
| Predictions Round 2 | IAFFMAFQ |
| Predictions Round 2 | IAFFMVFN |
| Predictions Round 2 | VPFFMIFN |
| Predictions Round 2 | LAFFIAFN |
| Predictions Round 2 | VGFFMVFQ |
| Predictions Round 2 | LAFFMAFQ |
| Predictions Round 2 | LYFFMAFN |
| Predictions Round 2 | VAFFIAFN |
| Predictions Round 2 | TAFFMAFN |
| Predictions Round 2 | TNVFMIFN |
| Predictions Round 2 | TAFFIAFN |
| Predictions Round 2 | VEFFMAFN |

| | |
|---|---|
| Predictions Round 2 | VGFFMAFN |
| Predictions Round 2 | LAFFIVFN |
| Predictions Round 2 | VPFFMAFQ |
| Predictions Round 2 | LGFFMAVN |
| Predictions Round 2 | LYFFMKFN |
| Predictions Round 2 | VAVFIAFN |
| Predictions Round 2 | LAFFMVFQ |
| Predictions Round 2 | VAFFMAFQ |
| Predictions Round 2 | TGFFMVFQ |
| Predictions Round 2 | TNFFMVFQ |
| Predictions Round 2 | LYFFMAFQ |
| Predictions Round 2 | TGFFMVFN |
| Predictions Round 2 | LYFFMVFN |
| Predictions Round 2 | TNFFMKFN |
| Predictions Round 2 | TNFFMVFN |
| Predictions Round 2 | TGFFMVYQ |
| Predictions Round 2 | VVFFIAFN |
| Predictions Round 2 | TAFFMAVN |
| Predictions Round 2 | TAFFMVVN |
| Predictions Round 2 | TAFFMIFN |
| Predictions Round 2 | LAFFQVFN |
| Predictions Round 2 | LAFFQAFN |
| Predictions Round 2 | LAFFMAYN |
| Predictions Round 2 | LNFFMVFQ |
| Predictions Round 2 | LYFFMVFQ |
| Predictions Round 2 | LAFFIAFQ |
| Predictions Round 2 | VYFFMAFQ |
| Predictions Round 2 | VYFFMVFN |
| Predictions Round 2 | TAIFMVFQ |
| Predictions Round 2 | VVFFMAFN |
| Predictions Round 2 | VVFFMAFQ |
| Predictions Round 2 | VAFFMVFN |
| Predictions Round 2 | TVFFMVFN |
| Predictions Round 2 | IAQFMAFN |
| Predictions Round 2 | IAFFMAFN |
| Predictions Round 2 | TAFFMVFN |
| Predictions Round 2 | TAFFMAFQ |
| Predictions Round 2 | TNFFMMFN |
| Predictions Round 2 | LNFFIVFN |

| | |
|---|---|
| Predictions Round 2 | LGFFMAFQ |
| Predictions Round 2 | LAVFIAFN |
| Predictions Round 2 | VAFFMAFN |
| Predictions Round 2 | VAFFMVFQ |
| Predictions Round 2 | LAFFMVFN |
| Predictions Round 2 | LAFFMAFN |
| Predictions Round 2 | VAFFIAFQ |
| Predictions Round 2 | TNAFMIFN |
| Predictions Round 2 | VYFFMAFN |
| Predictions Round 2 | TAFFMVFQ |
| Predictions Round 2 | TAQFMAFN |
| Predictions Round 2 | LAFFMKFN |
| Predictions Round 2 | VYFFMVFQ |

### Protoglobin Library Screening Protocols

**Protocols for the Screening of Protoglobin Variants in 96-well Plate Format:**

*96-Well Plate Library Expression*

The wells of a 2-mL 96-well deep-well plate were filled with 400 µL LB-Amp. Previously generated 96-well plate glycerol stocks were removed from -80 °C storage and placed on dry ice. Multichannel pipet tips were used to scratch the frozen glycerol stock surface and used to inoculate the aforementioned deep-well plate. These overnight cultures were incubated at 37 ºC and shaken at 220 rpm for 16–18 hours. For expression cultures, the following morning 50 µL of the precultures were used to inoculate 900 µL TB-amp per well in 96-well deep-well plates. The expression cultures were initially incubated at 37 ºC and 220 rpm for 2.5 hours, at which point they were allowed to sit at room temperature for 30 minutes. Expression of proteins was induced with IPTG, and cellular heme production was increased with ALA. An induction mixture containing IPTG and ALA in TB-amp (50 µL) was added to each well such that the final concentrations of IPTG and ALA were 0.5 mM and 1.0 mM respectively. The total culture volumes were 1 mL per well. The plates were then incubated at 22 ºC and 220 rpm overnight.

*96-Well Plate Library Reaction Preparation*

Expression cultures containing *E. coli* expressing hemoproteins of interest were centrifuged at 4,000 × g for 10 minutes at 4 ºC. The supernatant was discarded, and nitrogen-free M9 minimal media (M9-N, 380 µL for carbene transfer reactions, 365 µL for nitrene transfer reactions) were added to each well. For nitrene transfer reactions, 5 µL of a solution of D-glucose dissolved in M9-N were added such that the final concentration of D-glucose in reactions was 20 mM. The pellets were resuspended in the medium via shaking at room temperature for 30 minutes. The plates were then pumped into a vinyl Coy anaerobic chamber (0–30 ppm $O_2$). For carbene transfer reactions, to each well were added 20 µL of a MeCN solution containing the reaction substrate and ethyl diazoacetate (EDA). The final reaction volume was 400 µL. For nitrene transfer reactions, to each well were added 20 µL of an EtOH solution containing the reaction substrate followed by 10 µL of an aqueous solution containing the desired nitrene precursor. The final reaction volume was 400 µL. The plates were then sealed carefully with a foil cover and shaken at room temperature for 16 hours in the Coy chamber.

*96-Well Plate Library Reaction Preparation for Analysis – Carbene Transfer*

Once the reactions were complete, plates were worked up for processing by adding 600 µL of a 1:1 solution of ethyl acetate:cyclohexane containing 1,3,5-trimethoxybenzene as an internal standard (1.0 mM concentration). A silicone sealing mat (AWSM1003S, ArcticWhite) was used to cover each of the plates, and the two layers were thoroughly mixed by rapid inversion of the plates. The plates were then centrifuged (5,000 × g for 5 minutes at room temperature) to separate the phases.

For reactions **C1**–**C4**, which were always run in parallel with one another for library screening, 50 µL from each reaction plate were mixed in a GC vial insert in a GC vial, and the pooled samples were analyzed by GC-FID. For all other carbene transfer reactions, a 200-µL aliquot of the organic layer was transferred to a GC vial insert in a GC vial, and the samples were assayed by GC-FID or GC-MS.

*96-Well Plate Library Reaction Preparation for Analysis – Nitrene Transfer*

Once the reactions were complete, plates were worked up for processing by adding 400 µL of MeCN. The plates were then sealed with foil and stored at -20 °C for 1 hour. The plates were then centrifuged (5,000 × g for 5 minutes at room temperature) to clarify the processed solution.

Afterwards, a 200-μL aliquot of the combined layers was transferred to microtiter plate which was subsequently sealed by heat-sealing foil. These solutions were assayed by LC-MS.

### Library Screening Details and Data

The following section contains descriptions on how data was generated for each reaction system analyzed in this work. **Table S11** describes the manner in which data were collected for each reaction. Note that in some cases the data used to train our learning model in either training round in this work differs from the final data collected for that reaction. Both the datasets used for model training in this work and the final collected activity data are made available at https://github.com/fhalab/mrALDE. For future applications of these data to MLDE model training and validation studies we recommend using the final collected activity data. Detailed descriptions of the data collection procedures for each individual reaction system are provided below.

**Table S11.** Overview of data collection methods for reactions investigated in this work. In some cases, data collected from the first round of predictions for model updating differed from a final round of data collection. A value of N/A in the "Data Collection Method – Final" column indicates that the same data were used for model training as are made available online.

| Reaction Name | Data Collection Method – Training | Data Collection Method – Final | Notes |
|---|---|---|---|
| C1 | GC-FID, pooled with reactions C2, C3, and C4 | N/A | |
| C2 | GC-FID, pooled with reactions C1, C3, and C4 | N/A | |
| C3 | GC-FID, pooled with reactions C1, C2, and C4 | N/A | |
| C4 | GC-FID, pooled with reactions C1, C2, and C3 | N/A | |
| C5 | GC-FID | N/A | |
| C6 | GC-FID | N/A | |
| C7 | GC-MS | N/A | |
| C8 | GC-FID | N/A | |
| C9 | GC-MS | N/A | |
| C10 | LC-MS | N/A | |
| C11 | GC-MS | N/A | |
| C12 | GC-FID | N/A | Data for this reaction was not used to update |

| | | | |
|---|---|---|---|
| | | | the model after Round 1 Predictions |
| C13 | GC-MS, pooled with reactions C14, C15, C16, and C17 | GC-FID, unpooled | |
| C14 | GC-MS, pooled with reactions C13, C15, C16, and C17 | GC-MS, unpooled | |
| C15 | GC-MS, pooled with reactions C13, C14, C16, and C17 | GC-MS, unpooled | |
| C16 | GC-MS, pooled with reactions C13, C14, C15, and C17 | GC-MS, unpooled | |
| C17 | GC-MS, pooled with reactions C13, C14, C15, and C16 | GC-MS, unpooled | |
| N1 | LC-MS | N/A | |
| N2 | LC-MS | N/A | |
| N3 | LC-MS | N/A | |
| N4 | LC-MS | N/A | |
| N5 | LC-MS | N/A | |
| N6 | LC-MS | N/A | |
| N7 | LC-MS | N/A | |
| N8 | LC-MS | | The exact product structure was not determined; activity was quantified based on an LC–MS peak corresponding to the expected [M+H]+ ion. |
| N9 | LC-MS | | During Round 1 data collection, several variants were initially assigned activity for reaction N9, and these measurements were incorporated into the model update. Subsequent reanalysis of these samples did not confirm product formation under the reported conditions. |

#### Reaction C1:

**Scheme S1.** Reaction conditions for reaction **C1** and expected products.

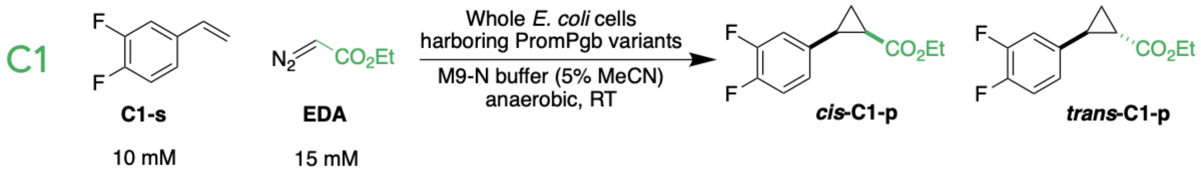


For library screening, reaction **C1** was analyzed by GC-FID using an equal part mixture of reaction extracts from reactions **C1**, **C2**, **C3**, and **C4**. Authentic standards for this reaction were synthesized and characterized in our laboratory. Reaction **C1** was tested with the initial training library, round 1 of predictions, and round 2 of predictions.

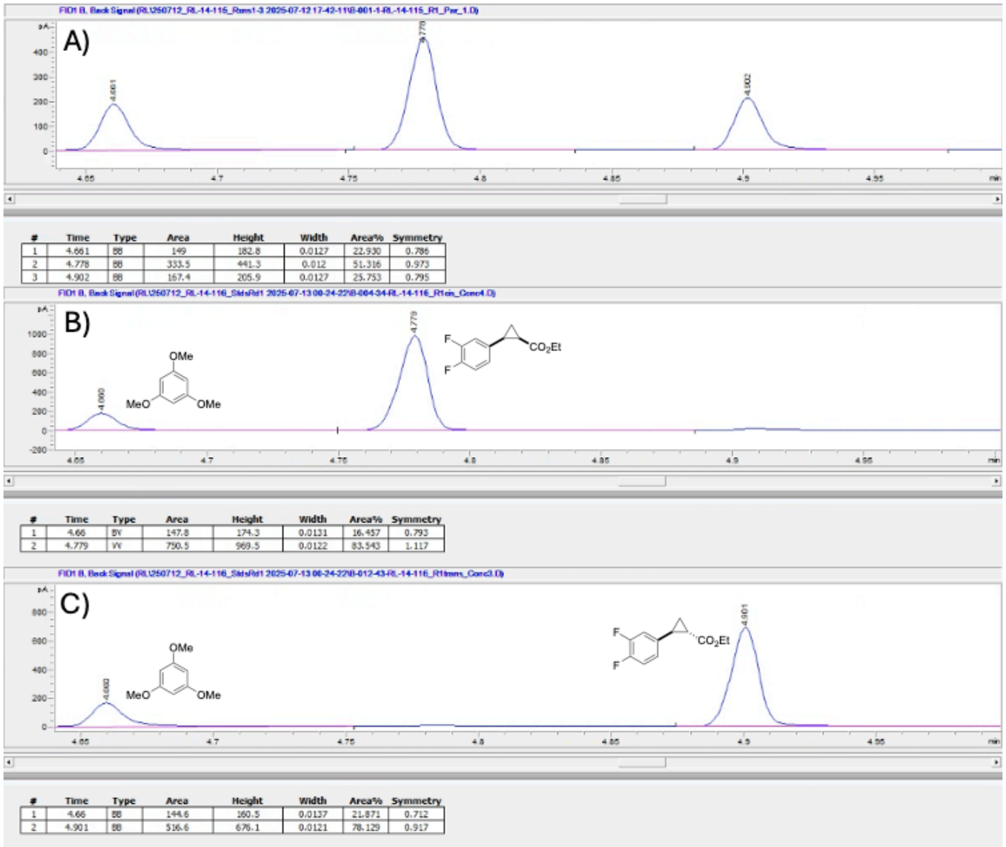


**Figure S22. (A)** Representative GC-FID trace for reaction **C1** with PromPgb. **(B)** GC-FID trace for a sample of the authentic standard of ***cis*-C1-p** with 1,3,5-trimethoxybenzene as an authentic standard. **(C)** GC-FID trace for a sample of the authentic standard of ***trans*-C1-p** with 1,3,5-trimethoxybenzene as an authentic standard.

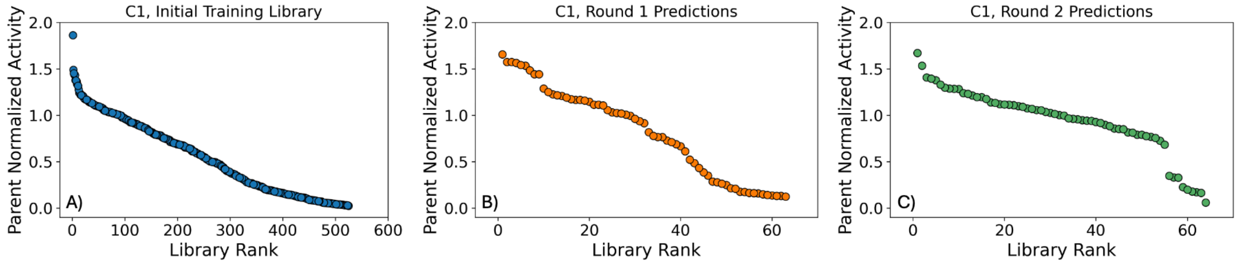


**Figure S23.** Retention of function plots for reaction **C1** activity for variants in the **(A)** initial training library, **(B)** the first round of predictions, and **(C)** the second round of predictions. Parent normalized activities are computed from the total formation of cyclopropane product for each variant relative to PromPgb.

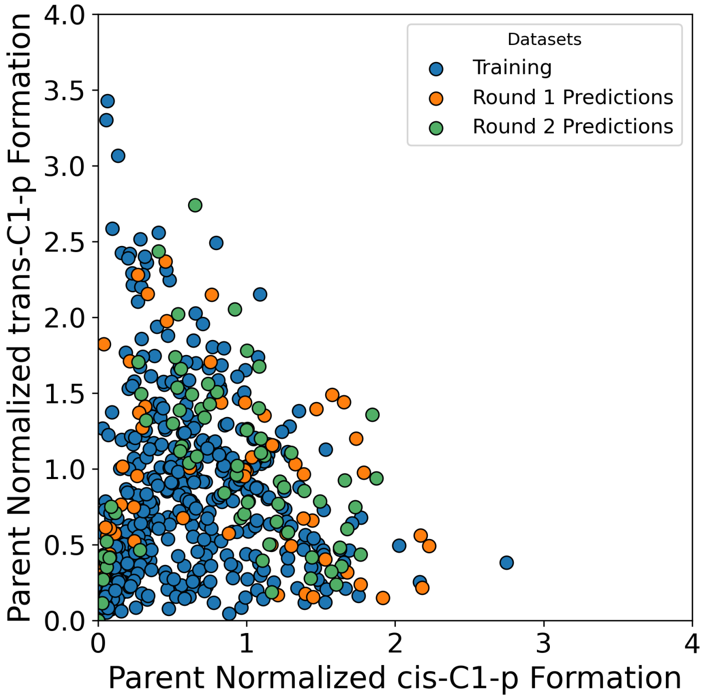


**Figure S24.** Changes in the formation of ***cis*-C1-p** and ***trans*-C1-p** for tested libraries. Parent normalized activities are computed from the formation of each cyclopropane diastereomer relative to PromPgb.

#### Reaction C2:

**Scheme S2.** Reaction conditions for reaction **C2** and expected products.

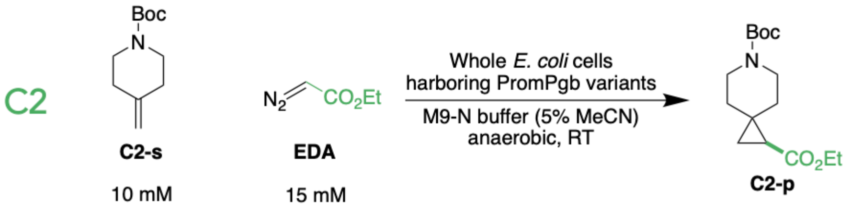

For library screening, reaction **C2** was analyzed by GC-FID in an equal part mixture of reaction extracts from reactions **C1**, **C2**, **C3**, and **C4**. The authentic standard of **C2-p** was kindly provided by Dr. Jae L. Kennemur, who previously synthesized and characterized this compound in our laboratory. Full characterization data are reported in reference 7. Reaction **C2** was tested with the initial training library, round 1 of predictions, and round 2 of predictions.

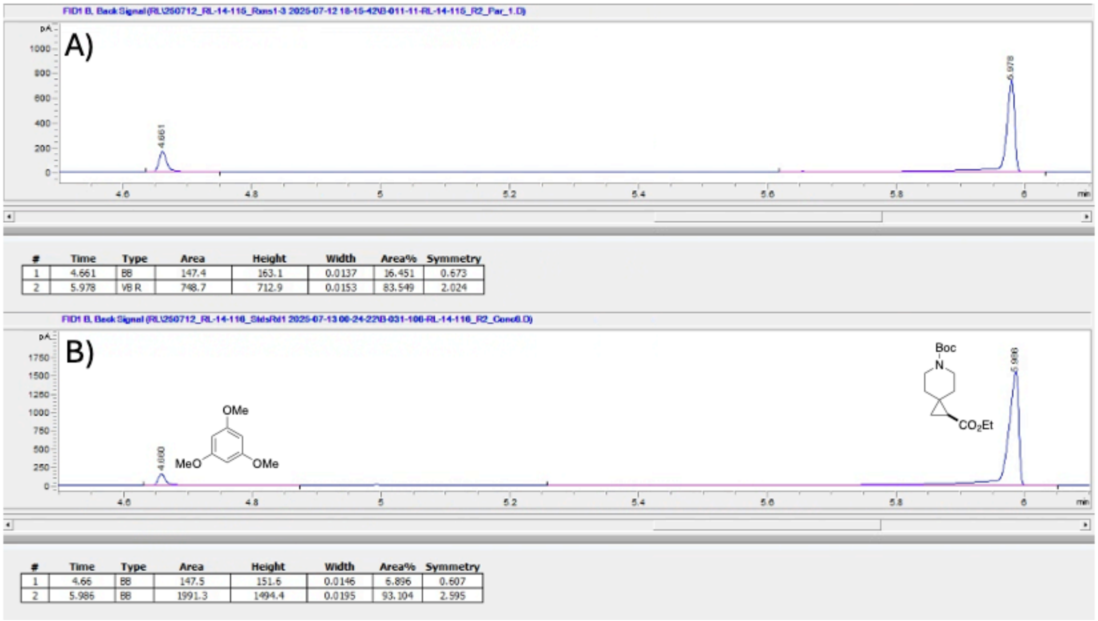


**Figure S24. (A)** Representative GC-FID trace for reaction **C2** with PromPgb. **(B)** GC-FID trace for a sample of the authentic standard of **C2-p** with 1,3,5-trimethoxybenzene as an authentic standard.

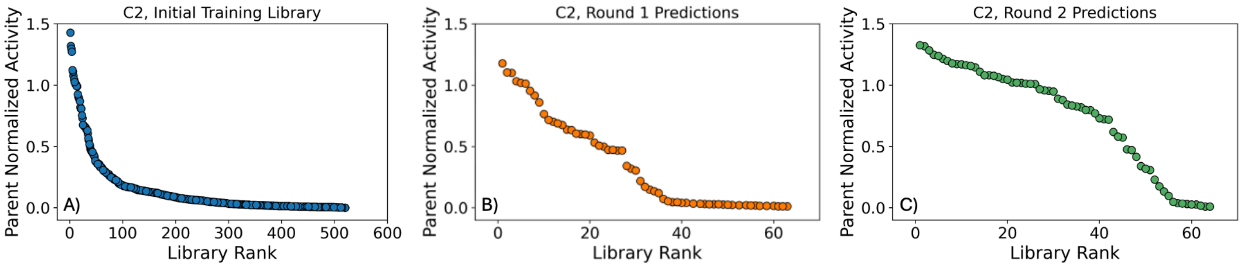


**Figure S25.** Retention of function plots for reaction **C2** activity for variants in the **(A)** initial training library, **(B)** round 1 of predictions and **(C)** round 2 of predictions. Parent normalized activities are computed from the total formation of cyclopropane product for each variant relative to PromPgb.

#### Reaction C3:

**Scheme S3.** Reaction conditions for reaction **C3** and expected products.

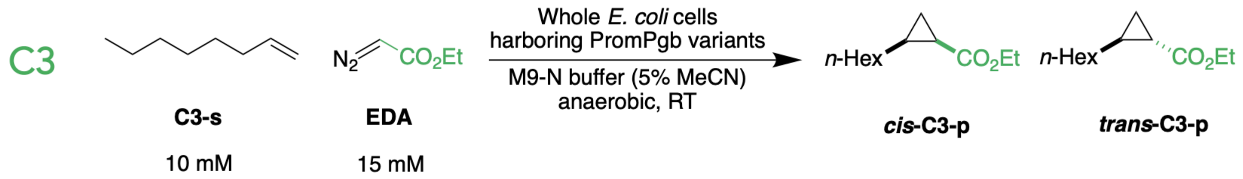


For library screening, reaction **C3** was analyzed by GC-FID in an equal part mixture of reaction extracts from reactions **C1**, **C2**, **C3**, and **C4**. Authentic standards for this reaction were synthesized and characterized in our laboratory. Reaction **C3** was tested with the initial training library, round 1 of predictions, and round 2 of predictions.

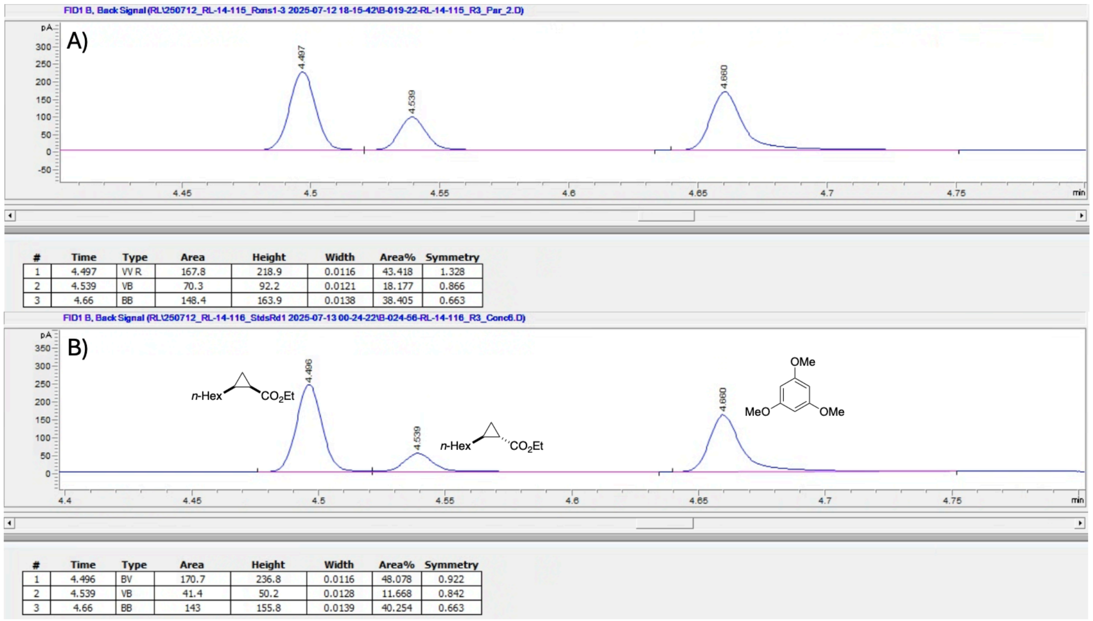


| # | Time | Type | Area | Height | Width | Area% | Symmetry |
|---|---|---|---|---|---|---|---|
| 1 | 4.497 | VV R | 167.8 | 218.9 | 0.0116 | 43.418 | 1.328 |
| 2 | 4.539 | VB | 70.3 | 92.2 | 0.0121 | 18.177 | 0.866 |
| 3 | 4.66 | BB | 148.4 | 163.9 | 0.0138 | 38.405 | 0.663 |

| # | Time | Type | Area | Height | Width | Area% | Symmetry |
|---|---|---|---|---|---|---|---|
| 1 | 4.496 | BV | 170.7 | 236.8 | 0.0116 | 48.078 | 0.922 |
| 2 | 4.539 | VB | 41.4 | 50.2 | 0.0128 | 11.668 | 0.842 |
| 3 | 4.66 | BB | 143 | 155.8 | 0.0139 | 40.254 | 0.663 |

**Figure S26. (A)** Representative GC-FID trace for reaction **C3** with PromPgb. **(B)** GC-FID trace for a sample of mixture of the authentic standards of ***cis*-C3-p** and ***trans*-C3-p** with 1,3,5-trimethoxybenzene as an authentic standard.

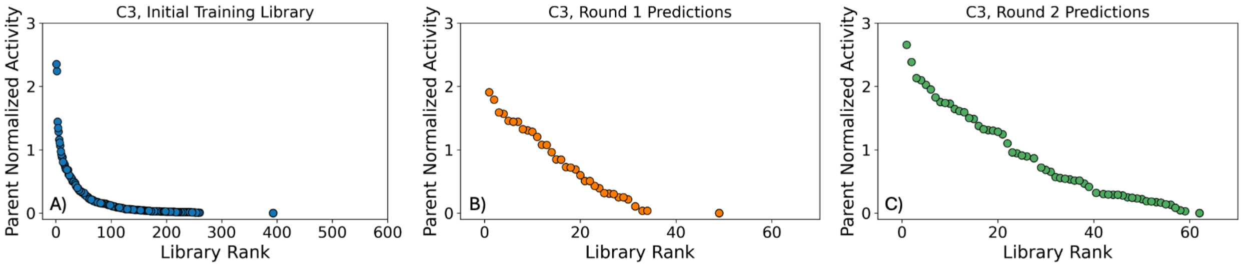


**Figure S27.** Retention of function plots for reaction **C3** activity for variants in the **(A)** initial training library, **(B)** the first round of predictions, and **(C)** the second round of predictions. Parent normalized activities are computed from the total formation of cyclopropane product for each variant relative to PromPgb.

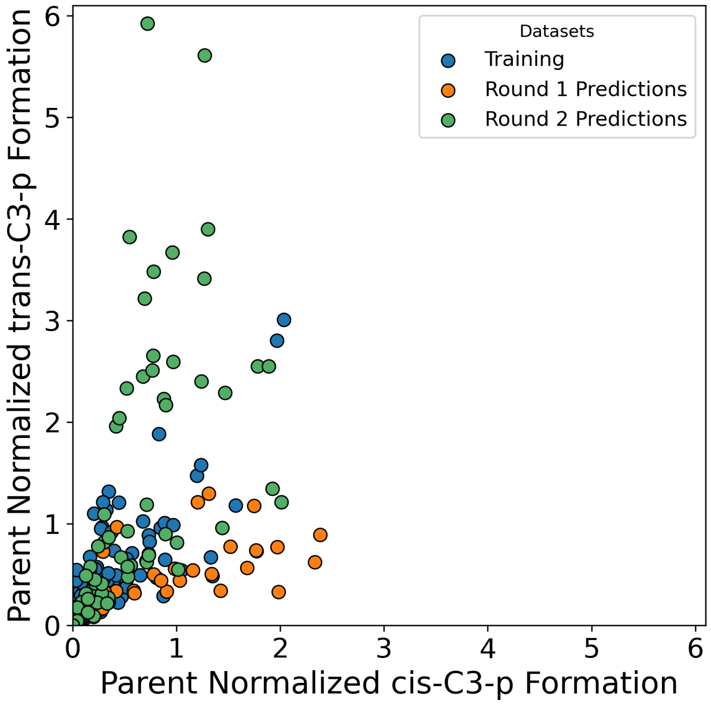


**Figure S28.** Changes in the formation of ***cis*-C3-p** and ***trans*-C3-p** for tested libraries. Parent normalized activities are computed from the formation of each cyclopropane diastereomer relative to PromPgb.

#### Reaction C4:

**Scheme S4.** Reaction conditions for reaction **C4** and expected products.

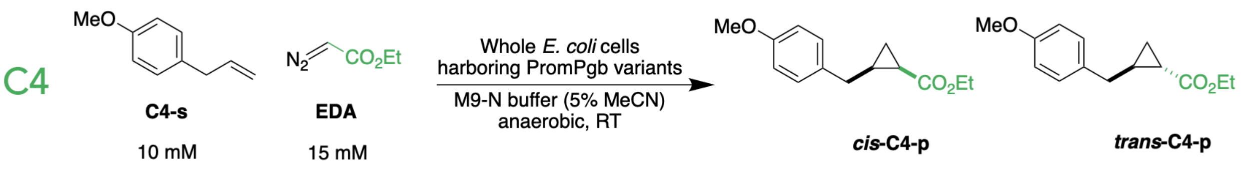


For library screening, reaction **C4** was analyzed by GC-FID in an equal part mixture of reaction extracts from reactions **C1**, **C2**, **C3**, and **C4**. Authentic standards for this reaction were synthesized and characterized in our laboratory. Reaction **C4** was tested with the initial training library, round 1 of predictions, and round 2 of predictions.

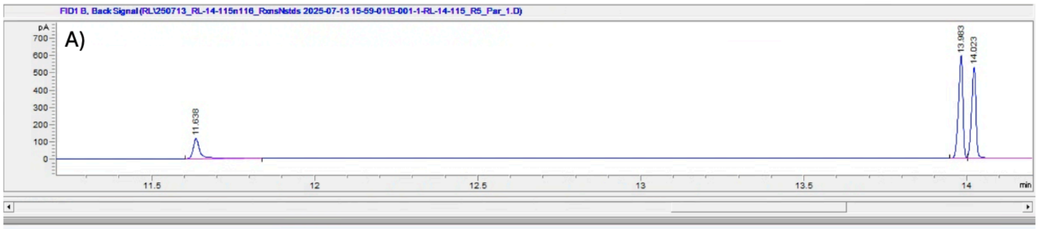


| # | Time | Type | Area | Height | Width | Area% | Symmetry |
|---|---|---|---|---|---|---|---|
| 1 | 11.638 | BB | 162.9 | 112.9 | 0.0216 | 13.266 | 0.616 |
| 2 | 13.983 | BV | 558 | 584.9 | 0.0151 | 45.425 | 1.357 |
| 3 | 14.023 | VB | 507.4 | 520.3 | 0.0154 | 41.310 | 1.136 |

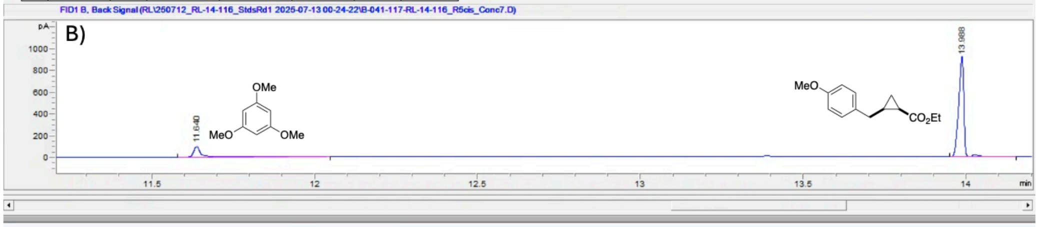


| # | Time | Type | Area | Height | Width | Area% | Symmetry |
|---|---|---|---|---|---|---|---|
| 1 | 11.64 | BB | 146.7 | 96.3 | 0.0226 | 12.388 | 0.567 |
| 2 | 13.988 | BV R | 1037.3 | 910.2 | 0.0168 | 87.612 | 1.583 |

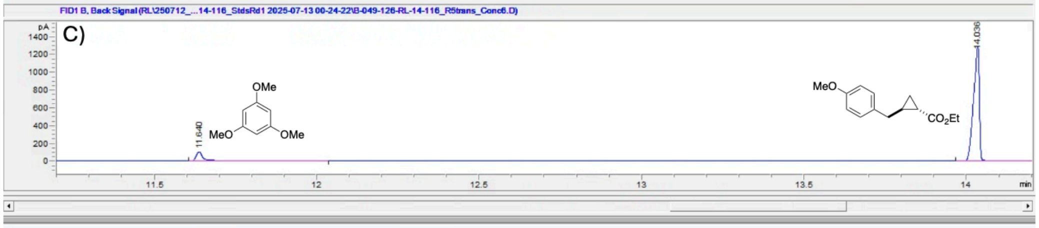


| # | Time | Type | Area | Height | Width | Area% | Symmetry |
|---|---|---|---|---|---|---|---|
| 1 | 11.64 | BB | 150.1 | 98.7 | 0.0225 | 8.610 | 0.566 |
| 2 | 14.036 | BB | 1593.3 | 1273.1 | 0.0186 | 91.390 | 2.014 |

**Figure S29. (A)** Representative GC-FID trace for reaction **C4** with PromPgb. **(B)** GC-FID trace for a sample of the authentic standard of ***cis*-C4-p** with 1,3,5-trimethoxybenzene as an authentic standard. **(C)** GC-FID trace for a sample of the authentic standard of ***trans*-C4-p** with 1,3,5-trimethoxybenzene as an authentic standard.

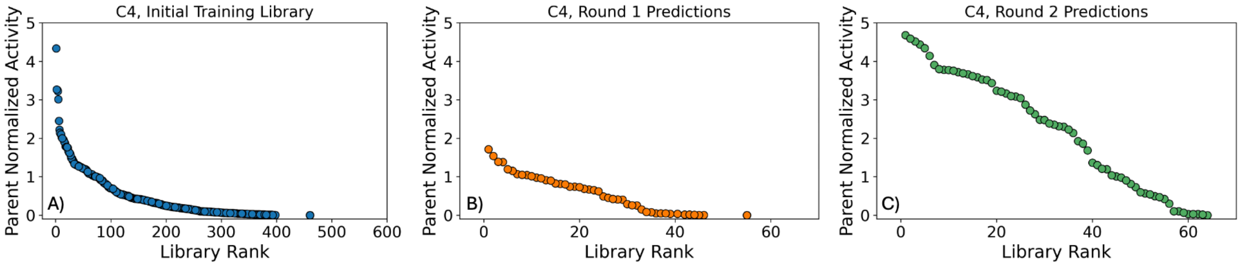


**Figure S30.** Retention of function plots for reaction **C4** activity for variants in the **(A)** initial training library, **(B)** the first round of predictions, and **(C)** the second round of predictions. Parent normalized activities are computed from the total formation of cyclopropane product for each variant relative to PromPgb.

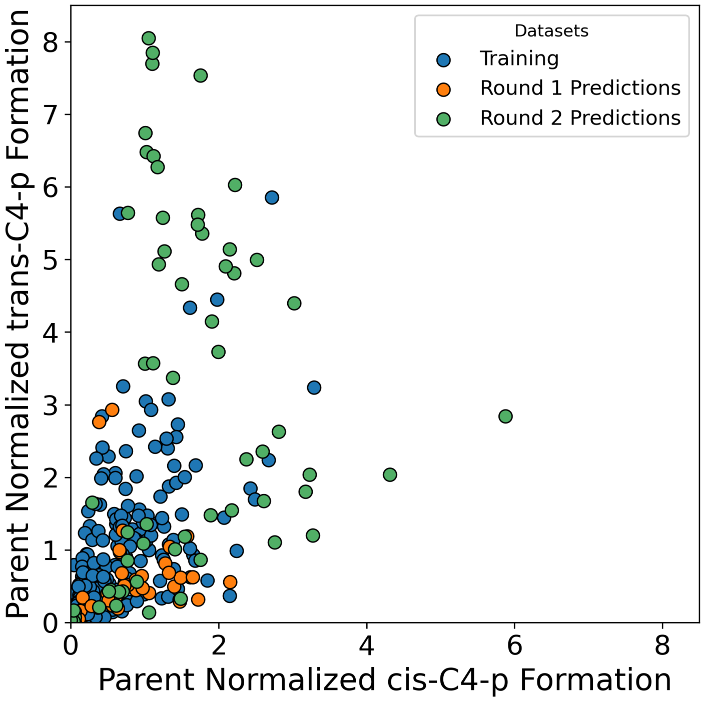


**Figure S31.** Changes in the formation of ***cis*-C4-p** and ***trans*-C4-p** for tested libraries. Parent normalized activities are computed from the formation of each cyclopropane diastereomer relative to PromPgb.

#### Reaction C5:

**Scheme S5.** Reaction conditions for reaction **C5** and expected products.

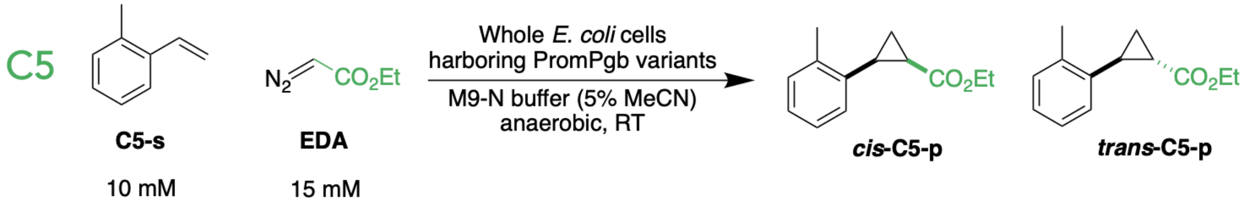


For library screening, reaction **C5** was analyzed by GC-FID. Authentic standards for this reaction were synthesized and characterized in our laboratory in a previous investigation. Full characterization data are reported in reference 8. Reaction **C5** was tested with round 2 of predictions.

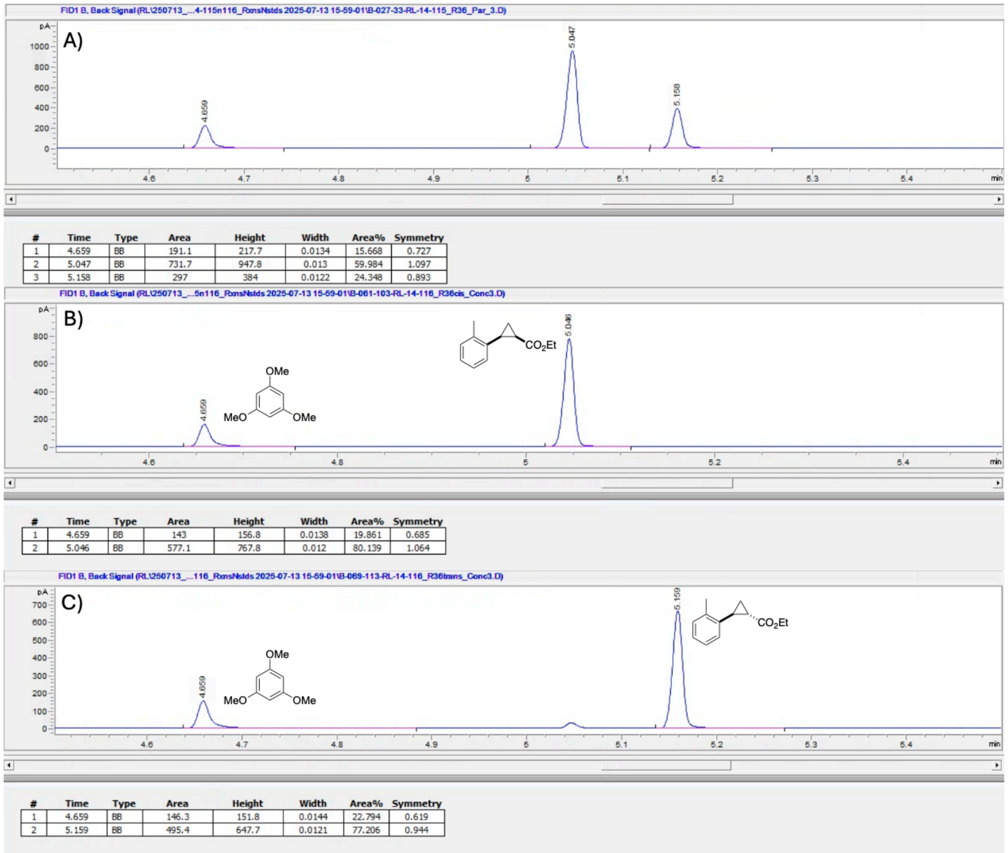


| # | Time | Type | Area | Height | Width | Area% | Symmetry |
|---|---|---|---|---|---|---|---|
| 1 | 4.659 | BB | 191.1 | 217.7 | 0.0134 | 15.668 | 0.727 |
| 2 | 5.047 | BB | 731.7 | 947.8 | 0.013 | 59.984 | 1.097 |
| 3 | 5.158 | BB | 297 | 384 | 0.0122 | 24.348 | 0.893 |

| # | Time | Type | Area | Height | Width | Area% | Symmetry |
|---|---|---|---|---|---|---|---|
| 1 | 4.659 | BB | 143 | 156.8 | 0.0138 | 19.861 | 0.685 |
| 2 | 5.046 | BB | 577.1 | 767.8 | 0.012 | 80.139 | 1.064 |

| # | Time | Type | Area | Height | Width | Area% | Symmetry |
|---|---|---|---|---|---|---|---|
| 1 | 4.659 | BB | 146.3 | 151.8 | 0.0144 | 22.794 | 0.619 |
| 2 | 5.159 | BB | 495.4 | 647.7 | 0.0121 | 77.206 | 0.944 |

**Figure S32. (A)** Representative GC-FID trace for reaction **C5** with PromPgb. **(B)** GC-FID trace for a sample of the authentic standard of ***cis*-C5-p** with 1,3,5-trimethoxybenzene as an authentic standard. **(C)** GC-FID trace for a sample of the authentic standard of ***trans*-C5-p** with 1,3,5-trimethoxybenzene as an authentic standard.

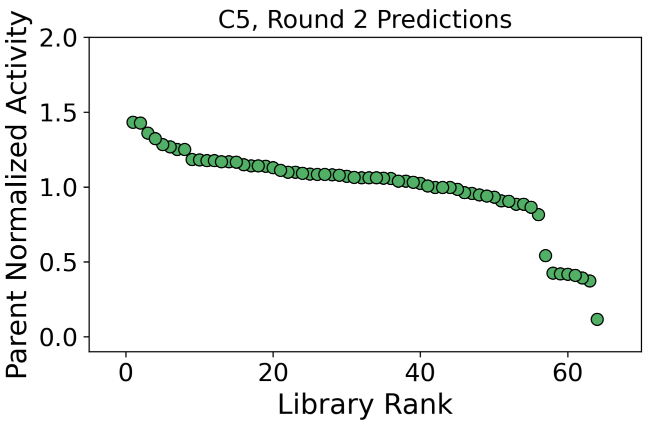


**Figure S33.** Retention of function plot for reaction **C5** activity for variants in the second round of predictions. Parent normalized activities are computed from the total formation of cyclopropane product for each variant relative to PromPgb.

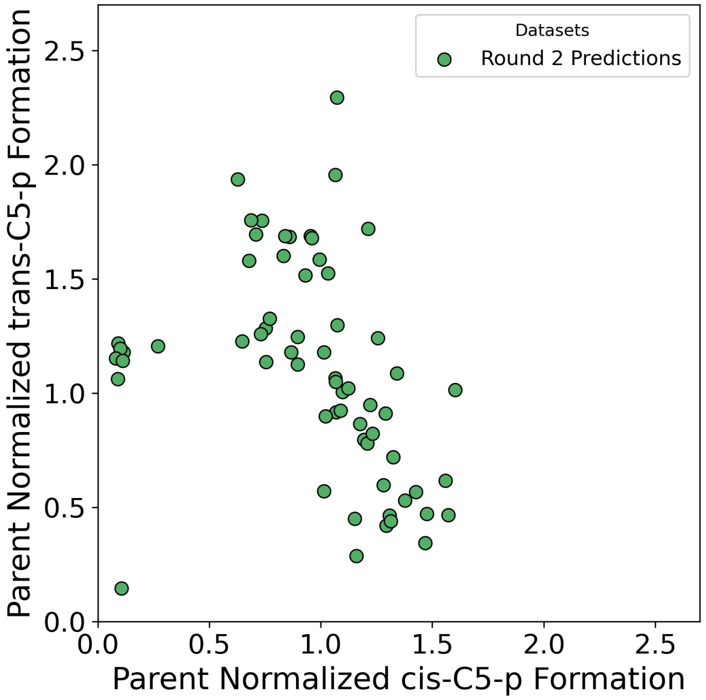


**Figure S34.** Changes in the formation of ***cis*-C4-p** and ***trans*-C4-p** for tested libraries. Parent normalized activities are computed from the formation of each cyclopropane diastereomer relative to PromPgb.

#### Reaction C6:

**Scheme S6.** Reaction conditions for reaction **C6** and expected products.

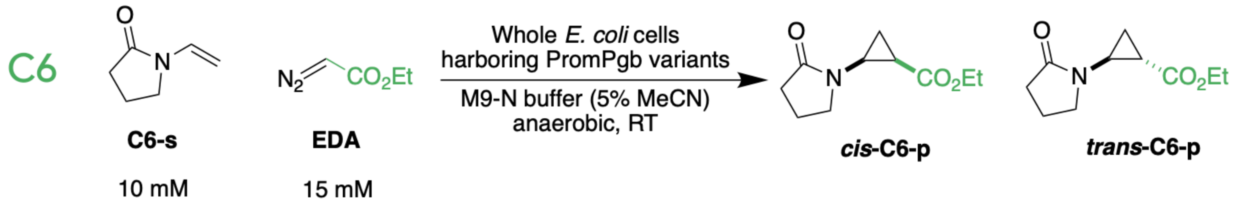


For library screening, reaction **C6** was analyzed by GC-FID. Authentic standards for this reaction were synthesized and characterized in our laboratory. Reaction **C6** was tested with round 2 of predictions.

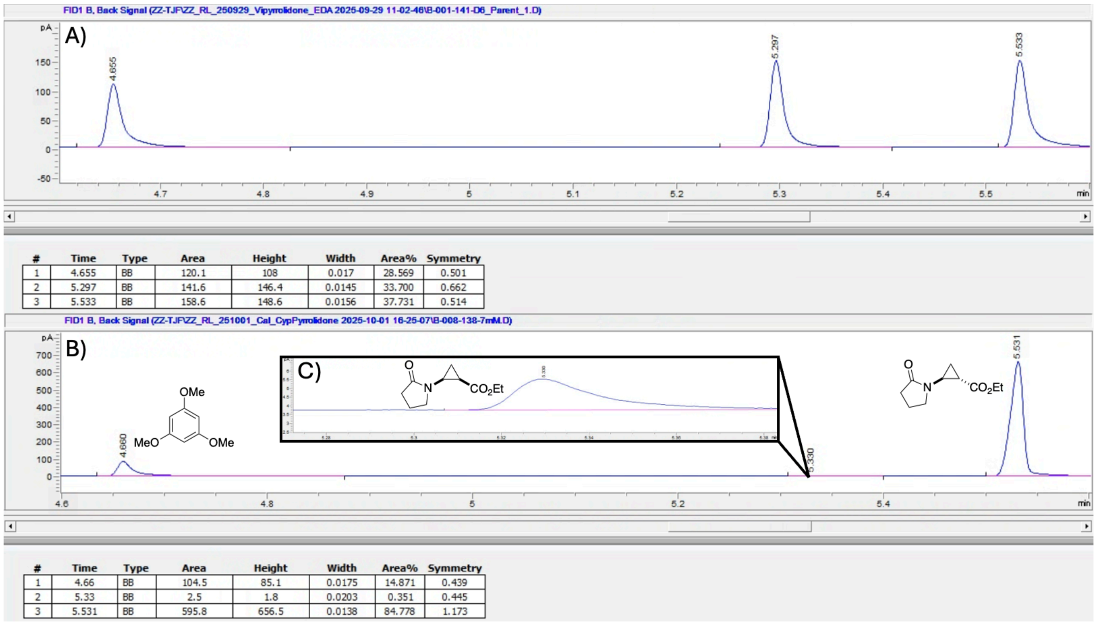


| # | Time | Type | Area | Height | Width | Area% | Symmetry |
|---|---|---|---|---|---|---|---|
| 1 | 4.655 | BB | 120.1 | 108 | 0.017 | 28.569 | 0.501 |
| 2 | 5.297 | BB | 141.6 | 146.4 | 0.0145 | 33.700 | 0.662 |
| 3 | 5.533 | BB | 158.6 | 148.6 | 0.0156 | 37.731 | 0.514 |

| # | Time | Type | Area | Height | Width | Area% | Symmetry |
|---|---|---|---|---|---|---|---|
| 1 | 4.66 | BB | 104.5 | 85.1 | 0.0175 | 14.871 | 0.439 |
| 2 | 5.33 | BB | 2.5 | 1.8 | 0.0203 | 0.351 | 0.445 |
| 3 | 5.531 | BB | 595.8 | 656.5 | 0.0138 | 84.778 | 1.173 |

**Figure S35. (A)** Representative GC-FID trace for reaction **C5** with PromPgb. **(B)** GC-FID trace for a sample of the authentic standard of ***cis*-C6-p** and ***trans*-C6-p** with 1,3,5-trimethoxybenzene as an authentic standard. **(C)** The authentic standard for ***cis*-C6-p** was isolated in trace quantities during synthesis.

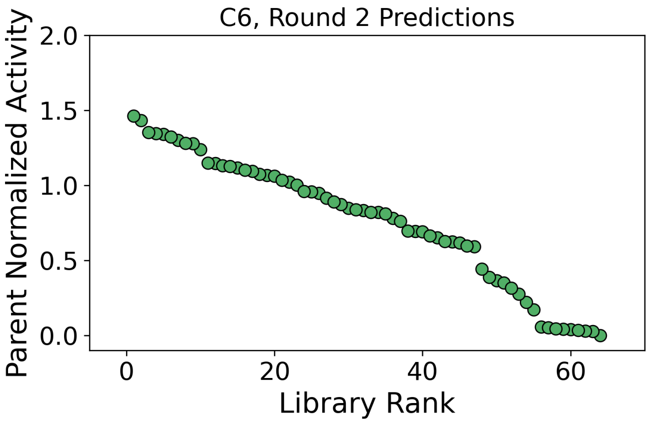


**Figure S36.** Retention of function plot for reaction **C6** activity for variants in the second round of predictions. Parent normalized activities are computed from the total formation of cyclopropane product for each variant relative to PromPgb.

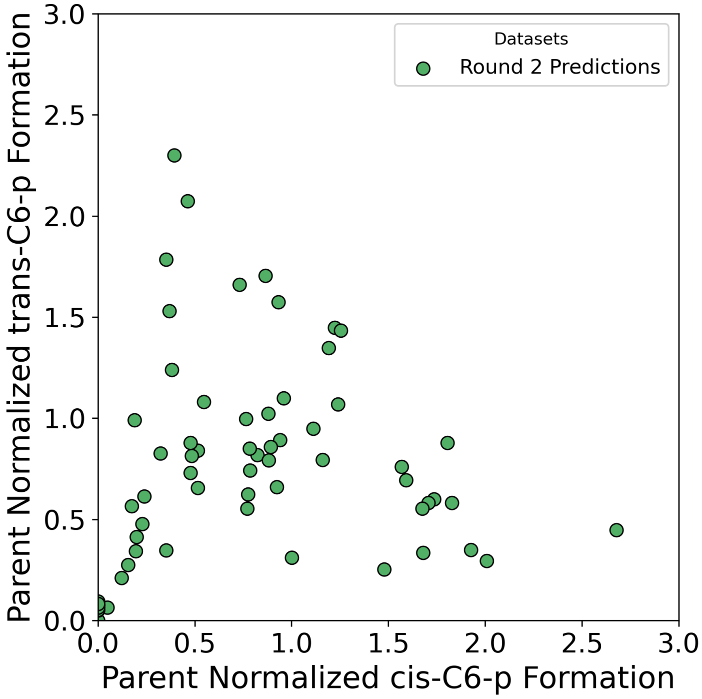


**Figure S37.** Changes in the formation of ***cis*-C6-p** and ***trans*-C6-p** for tested libraries. Parent normalized activities are computed from the formation of each cyclopropane diastereomer relative to PromPgb.

#### Reaction C7:

**Scheme S7.** Reaction conditions for reaction **C7** and expected products.

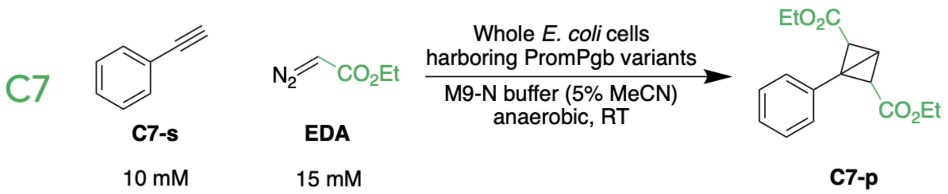


For library screening, reaction **C7** was analyzed by GC-MS. The authentic standard was obtained by scaling up the enzymatic reaction PromPgb, followed by purification and characterization. Reaction **C7** was tested with round 1 and round 2 of predictions. The bicyclobutane products obtained in reaction **C7** did not provide well-resolved peaks under the applied GC conditions; therefore, reaction yields were not quantified. Instead, for model updating, data from this reaction were treated as binary, where a value of 0 denoted no reaction, and a value of 1 indicated detectable product formation.

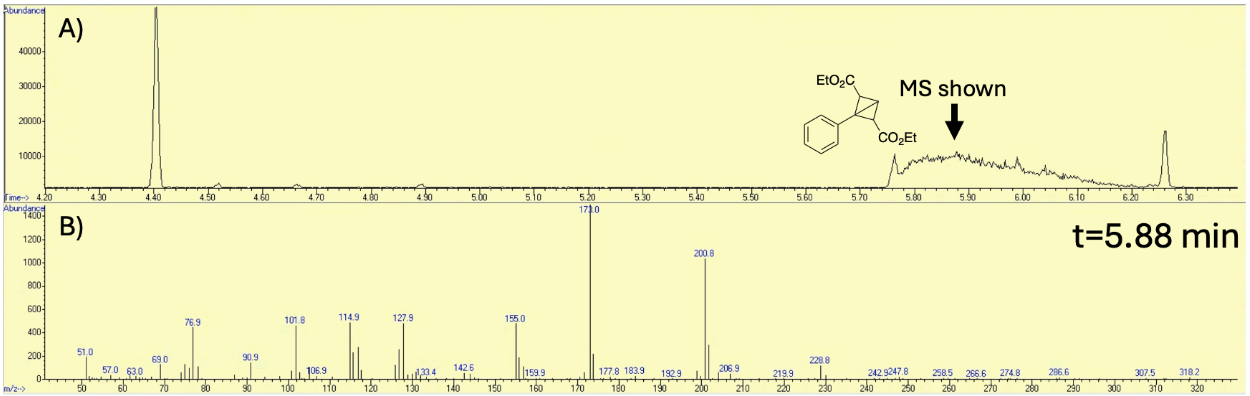


**Figure S38. (A)** Representative GC-MS trace for reaction **C7** with PromPgb. This trace was measured from an aliquot taken from the scaled-up reaction of PromPgb under conditions for **C7** to isolate **C7-p**. The bicyclobutane product does not provide a well-resolved peak. **(B)** Selected mass spectrum from t = 5.88 min, showing representative mass peaks for the bicyclobutane **C7-p**.

**Reaction C8:**

**Scheme S8.** Reaction conditions for reaction **C8** and expected products.

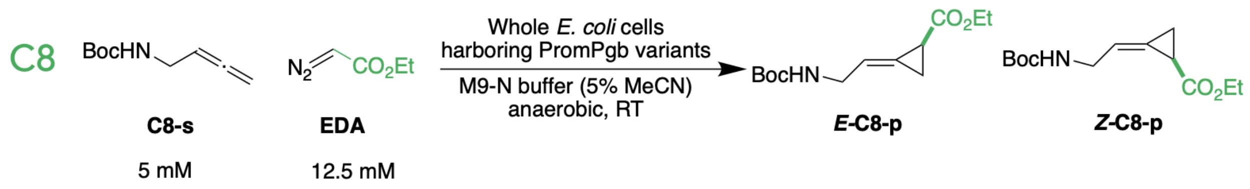


For library screening, reaction **C8** was analyzed by GC-FID. The authentic standards were obtained by scaling up the enzymatic reaction with variants VYIFMVFQ (***E*-C8-p**) and TAIFMVFQ (***Z*-C8-p**), followed by purification and characterization. Reaction **C8** was tested with round 1 and round 2 of predictions. The configuration of **C8-p** was assigned by analogy to reaction **C9**. NMR characterization of the product of reaction **C9** with PromPgb confirmed exclusive formation of ***E*-C9-p**. Given that PromPgb predominantly produces a single alkylidenecyclopropane isomer in both reactions **C8** and **C9** (**Figures S39A** and **Figure S42A**), the product for reaction **C8** with PromPgb was assigned as the *E*- configuration. The product isolated from variant TAIFMVFQ was thus assigned the *Z*- configuration (**Figure S39B**).

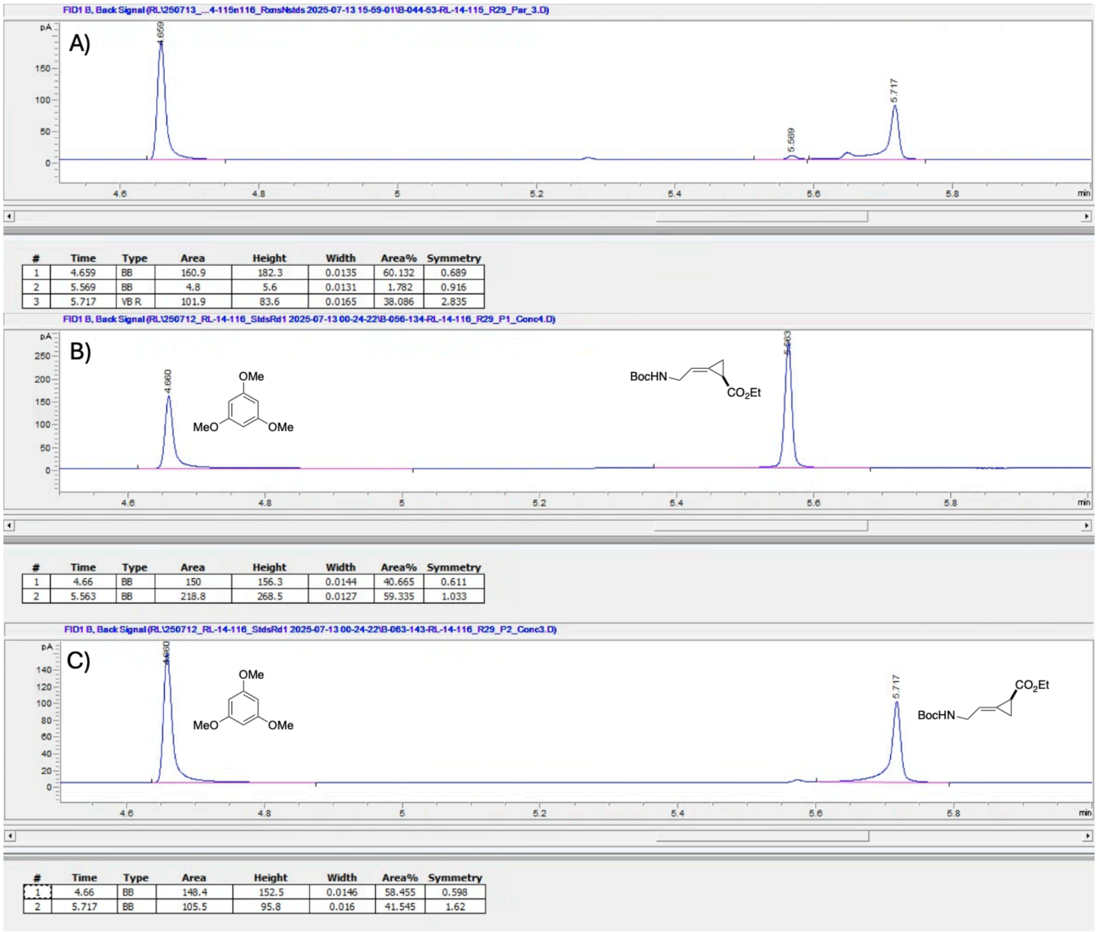


**Figure S39. (A)** Representative GC-FID trace for reaction **C8** with PromPgb. **(B)** GC-FID trace for a sample of the authentic standard of ***Z*-C8-p** with 1,3,5-trimethoxybenzene as an authentic standard. The authentic standard was isolated from a scaled-up reaction with variant TAIFMVFQ. **(C)** GC-FID trace for a sample of the authentic standard of ***E*-C8-p** with 1,3,5-trimethoxybenzene as an authentic standard. The authentic standard was isolated from a scaled-up reaction with variant VYIFMVFQ.

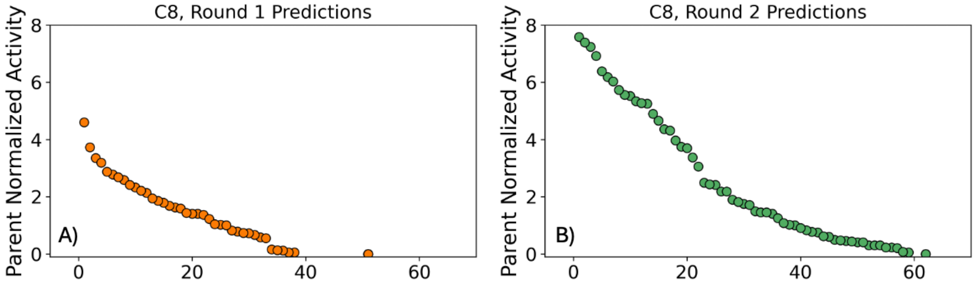


**Figure S40.** Retention of function plots for reaction **C8** activity for variants in the **(A)** first round of predictions and **(B)** the second round of predictions. Parent normalized activities are computed from the total formation of alkylidene cyclopropane product for each variant relative to PromPgb.

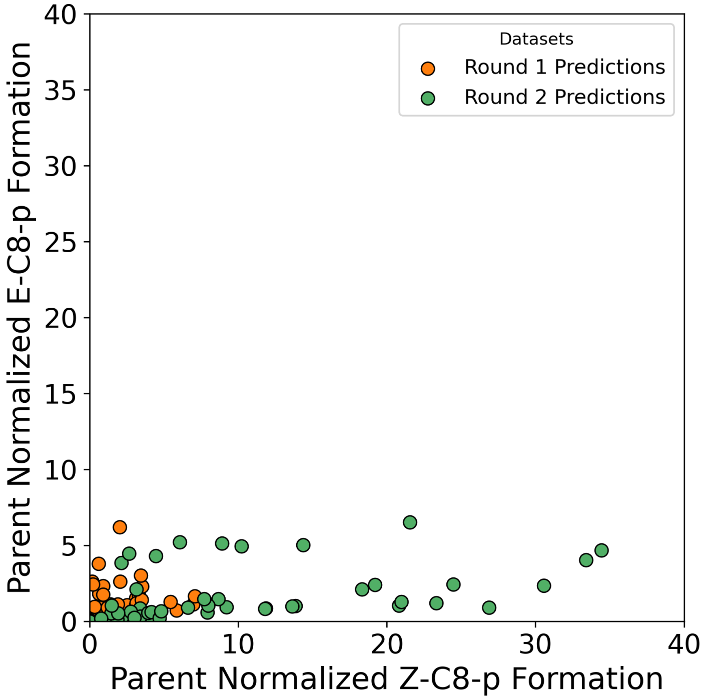


**Figure S41.** Changes in the formation of ***Z*-C8-p** and ***E*-C8-p** for tested libraries. Parent normalized activities are computed from the formation of each cyclopropane diastereomer relative to PromPgb.

#### Reaction C9:

**Scheme S9.** Reaction conditions for reaction **C9** and expected products.

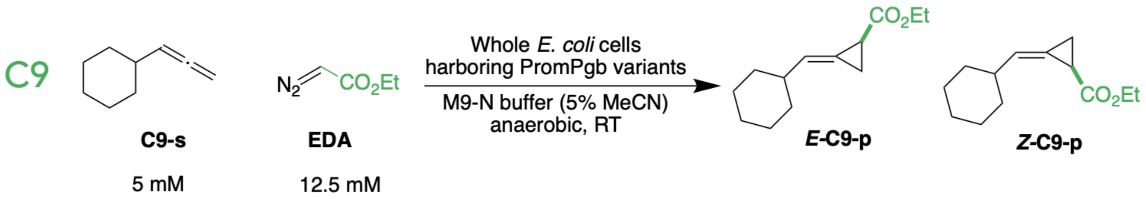


For library screening, reaction **C9** was analyzed by GC-FID. The authentic standards were obtained by scaling up the enzymatic reaction with variant LAFFIAFN, followed by purification and characterization. Reaction **C9** was tested with round 2 of predictions. In the course of library screening, only ***E*-C9-p** formation could be found under enzymatic conditions, as confirmed by NMR.[9] For model updating, data from this reaction were treated as binary, where a value of 0 denoted no reaction, and a value of 1 indicated detectable product formation.

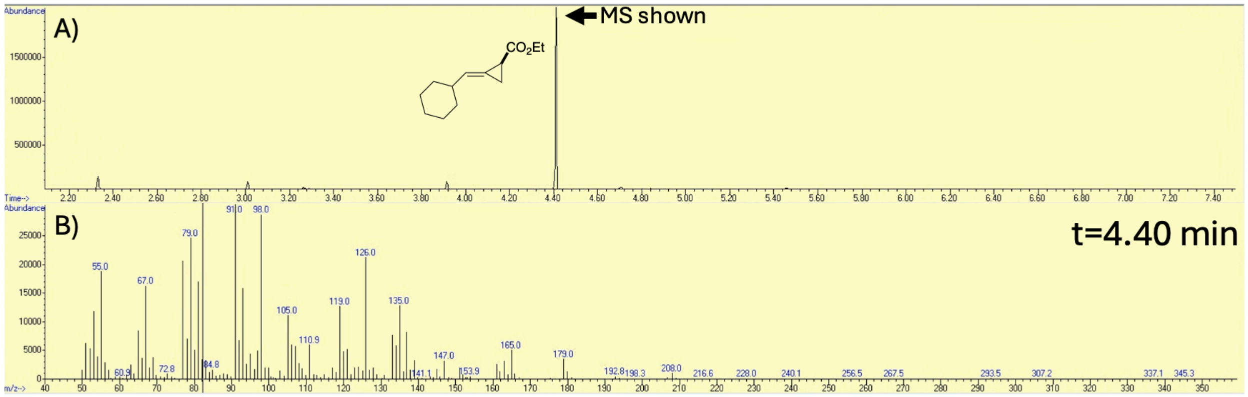


**Figure S42. (A)** Representative GC-MS trace for reaction **C9** with variant LAFFIAFN. This trace was measured from an aliquot taken from the scaled-up reaction of LAFFIAFN under conditions for **C9** to isolate **C9-p** products. Only ***E*-C9-p** could be isolated. **(B)** Selected mass spectrum from t = 4.40 min, showing representative mass peaks for the alkylidene cyclopropane ***E*-C9-p**.

#### Reaction C10:

**Scheme S10.** Reaction conditions for reaction **C10** and expected products.

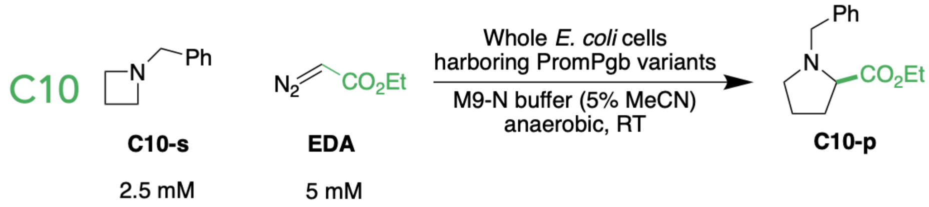


For library screening, reaction **C10** was analyzed by LC-MS. Authentic standards for this reaction were synthesized and characterized in our laboratory. Reaction **C10** was tested with round 1 and round 2 of predictions. PromPgb was not capable of performing this reaction, and no variants were found in either round of predictions which were capable of performing this transformation with detectable yield.

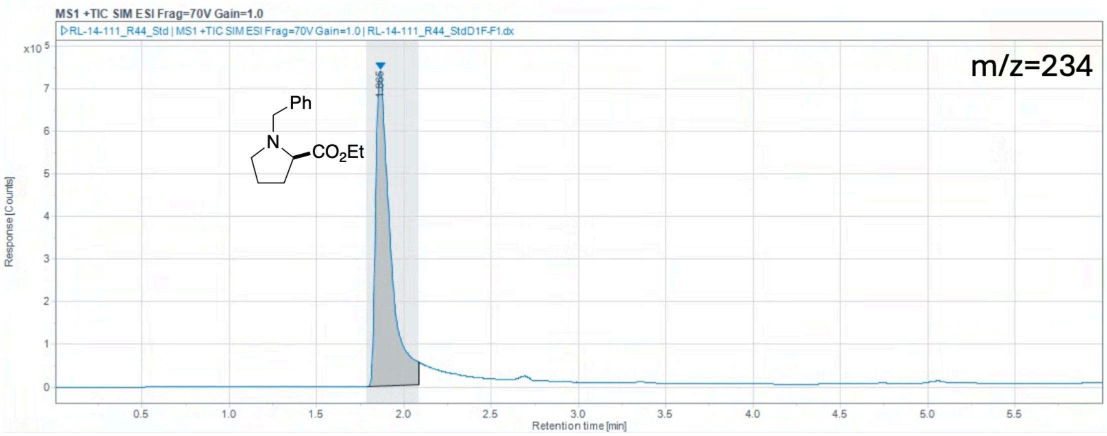


**Figure S43.** LC-MS trace for a sample of the authentic standard of **C10-p**. The Selective Ion Monitoring channel for the [M+H]+ ion of **C10-p** (m/z = 234) is shown.

#### Reaction C11:

**Scheme S11.** Reaction conditions for reaction **C11** and expected products.

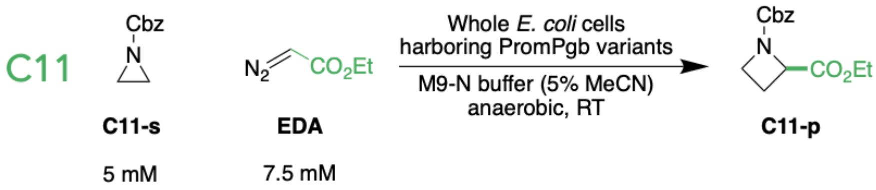


For library screening, reaction **C11** was analyzed by GC-MS. Authentic standards for this reaction were synthesized and characterized in our laboratory in a previous investigation. Full characterization data are reported in reference 10. Reaction **C11** was tested with round 1 and round 2 of predictions. PromPgb was not capable of performing this reaction, and no variants were found in either round of predictions which were capable of performing this transformation with detectable yield.

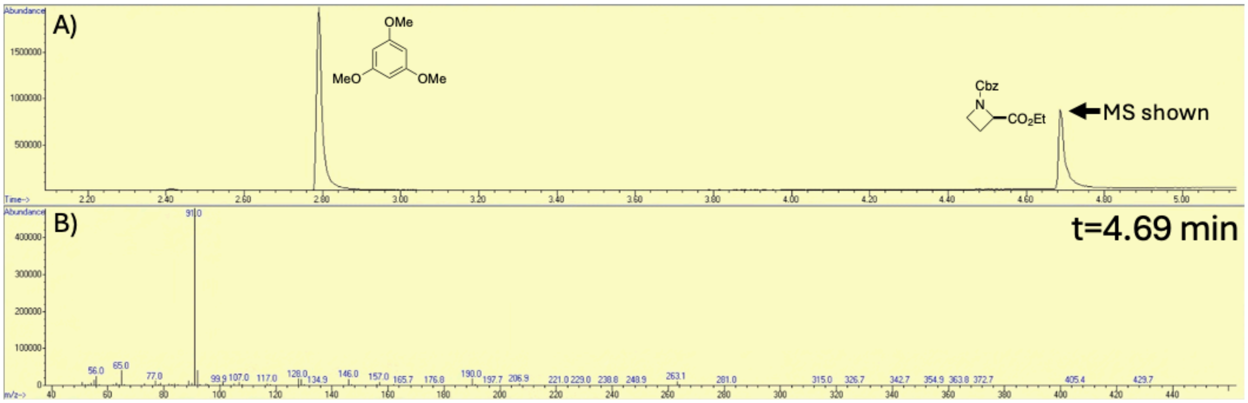


**Figure S44. (A)** GC-MS trace for an authentic standard of **C11-p**. **(B)** Selected mass spectrum from the authentic standard peak (t = 4.69 min), showing representative mass peaks for the ring expansion product **C11-p**.

#### Reaction C12:

**Scheme S12.** Reaction conditions for reaction **C12** and expected products.

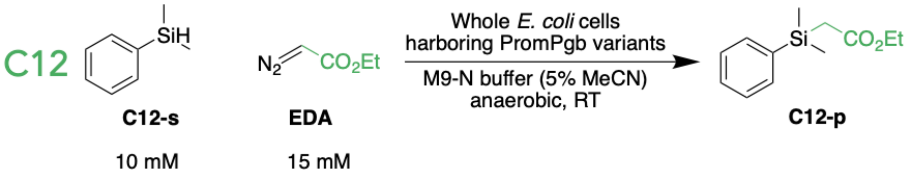


For library screening, reaction **C12** was analyzed by GC-FID. Authentic standards for this reaction were synthesized and characterized in our laboratory. Reaction **C12** was tested with round 1 and round 2 of predictions. Activity data for reaction **C12** with round 1 of predictions were not used to update the model before generating the second round of predictions in this work.

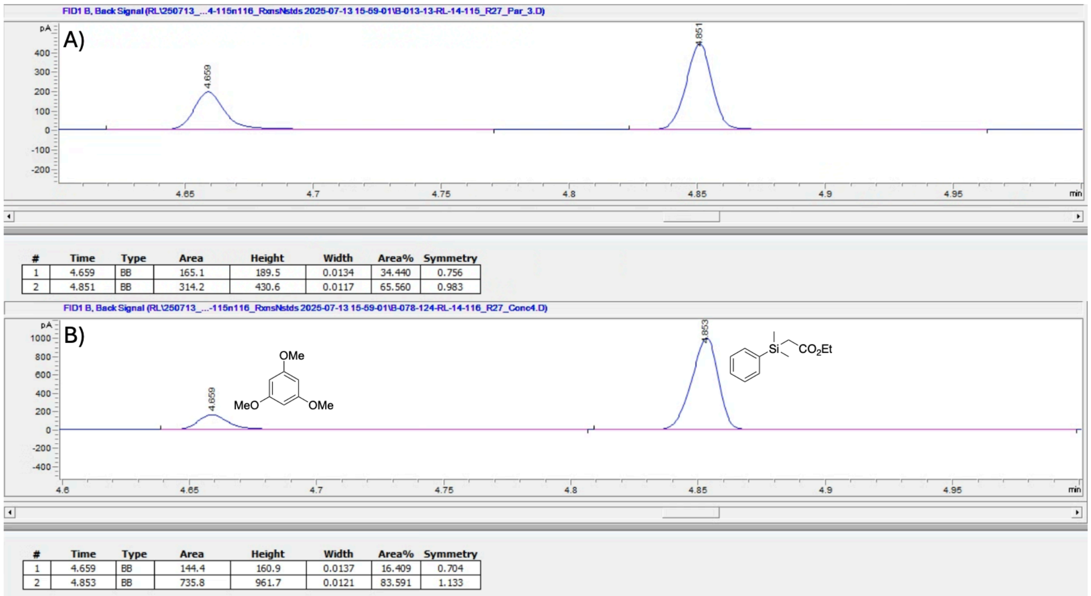


| # | Time | Type | Area | Height | Width | Area% | Symmetry |
|---|---|---|---|---|---|---|---|
| 1 | 4.659 | BB | 165.1 | 189.5 | 0.0134 | 34.440 | 0.756 |
| 2 | 4.851 | BB | 314.2 | 430.6 | 0.0117 | 65.560 | 0.983 |

| # | Time | Type | Area | Height | Width | Area% | Symmetry |
|---|---|---|---|---|---|---|---|
| 1 | 4.659 | BB | 144.4 | 160.9 | 0.0137 | 16.409 | 0.704 |
| 2 | 4.853 | BB | 735.8 | 961.7 | 0.0121 | 83.591 | 1.133 |

**Figure S45. (A)** Representative GC-FID trace for reaction **C12** with PromPgb. **(B)** GC-FID trace for a sample of the authentic standard of **C12-p** with 1,3,5-trimethoxybenzene as an authentic standard.

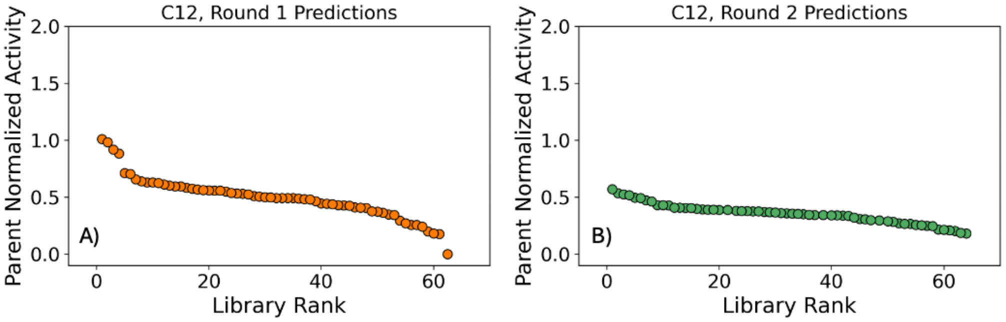


**Figure S46.** Retention of function plots for reaction **C12** activity for variants in the **(A)** first round of predictions and **(B)** the second round of predictions. Parent normalized activities are computed from the total formation of alkylation product for each variant relative to PromPgb.

**Reaction C13:**

**Scheme S13.** Reaction conditions for reaction **C13** and expected products.

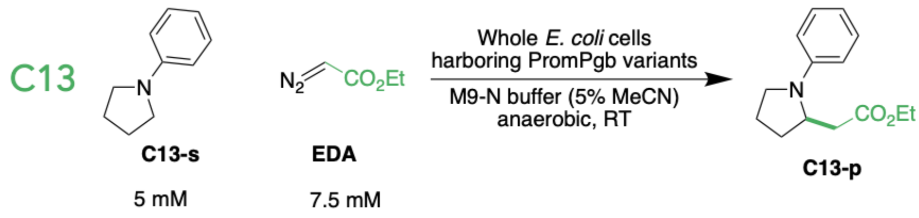


For library screening, reaction **C13** was analyzed by GC-FID. Authentic standards for this reaction were synthesized and characterized in our laboratory. Reaction **C13** was tested with round 1 and round 2 of predictions. While collecting data with round 1 predictions for model updating, a pooled GC-MS analysis was used for reaction **C13**, yielding qualitative reaction data. Reaction extracts from reactions **C13**, **C14**, **C15**, **C16**, and **C17** were pooled and analyzed by GC-MS. As a result, low-performing variants may have been over diluted and analytes may have been brought below the limit of detection. For model updating, data from this reaction were treated as binary, where a value of 0 denoted no reaction, and a value of 1 indicated detectable product formation. Thirty-seven variants were found to have activity for reaction **C13** for model updating in this work. Later, reaction **C13** was evaluated with round 1 and round 2 of predictions using a quantitative GC-FID method, yielding the data shown in **Figure S48**.

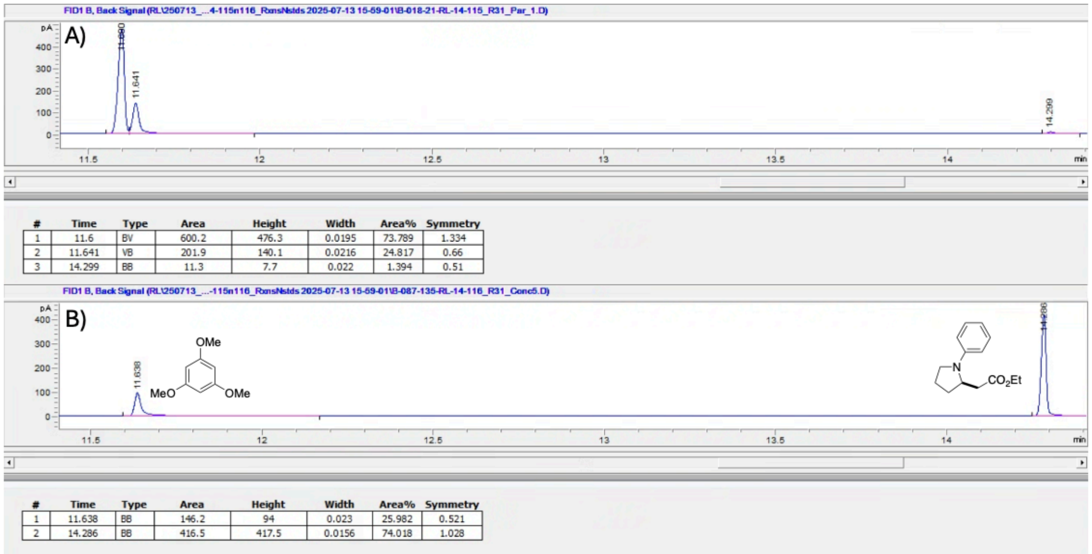


| # | Time | Type | Area | Height | Width | Area% | Symmetry |
|---|---|---|---|---|---|---|---|
| 1 | 11.6 | BV | 600.2 | 476.3 | 0.0195 | 73.789 | 1.334 |
| 2 | 11.641 | VB | 201.9 | 140.1 | 0.0216 | 24.817 | 0.66 |
| 3 | 14.299 | BB | 11.3 | 7.7 | 0.022 | 1.394 | 0.51 |

| # | Time | Type | Area | Height | Width | Area% | Symmetry |
|---|---|---|---|---|---|---|---|
| 1 | 11.638 | BB | 146.2 | 94 | 0.023 | 25.982 | 0.521 |
| 2 | 14.286 | BB | 416.5 | 417.5 | 0.0156 | 74.018 | 1.028 |

**Figure S47. (A)** Representative GC-FID trace for reaction **C13** with PromPgb. **(B)** GC-FID trace for a sample of the authentic standard of **C13-p** with 1,3,5-trimethoxybenzene as an authentic standard.

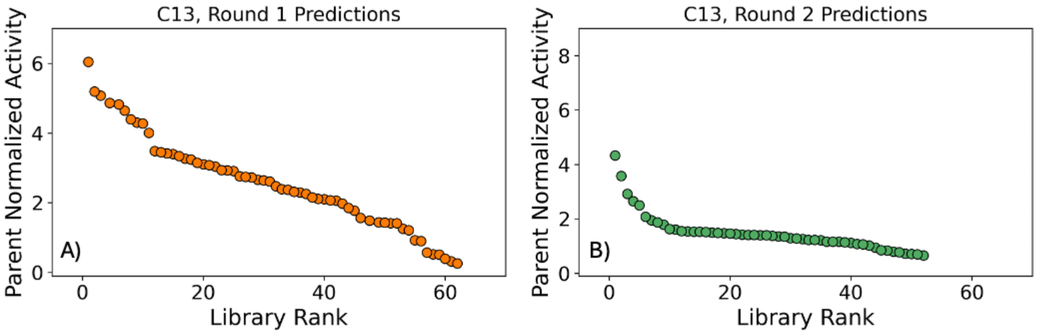


**Figure S48.** Retention of function plots for reaction **C13** activity for variants in the **(A)** first round of predictions and **(B)** the second round of predictions. Parent normalized activities are computed from the total formation of alkylation product for each variant relative to PromPgb.

**Reaction C14:**

**Scheme S14.** Reaction conditions for reaction **C14** and expected products.

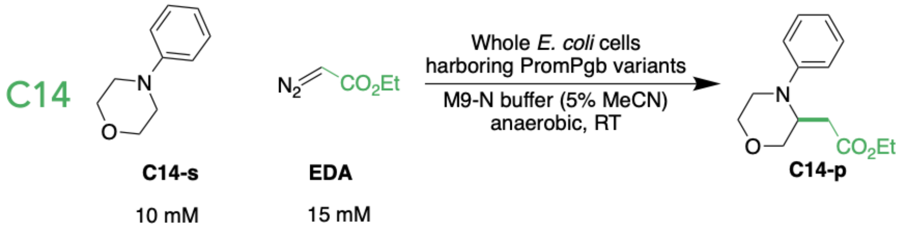


For library screening, reaction **C14** was analyzed by GC-MS. Authentic standards for this reaction were synthesized and characterized in our laboratory. Reaction **C14** was tested with round 1 and round 2 of predictions. PromPgb was incapable of catalyzing reaction **C14** under these conditions. While collecting data with round 1 predictions for model updating, a pooled GC-MS analysis was used for reaction **C14**, yielding qualitative reaction data. Reaction extracts from reactions **C13**, **C14**, **C15**, **C16**, and **C17** were pooled and analyzed by GC-MS. As a result, low-performing variants may have been over diluted and analytes may have been brought below the limit of detection. For model updating, data from this reaction were treated as binary, where a value of 0 denoted no reaction, and a value of 1 indicated detectable product formation. Six variants were found to have activity for reaction **C14** for model updating in this work. Later, reaction **C14** was evaluated with round 1 and round 2 of predictions using an un-pooled GC-MS strategy, uncovering 42 variants which could perform this transformation.

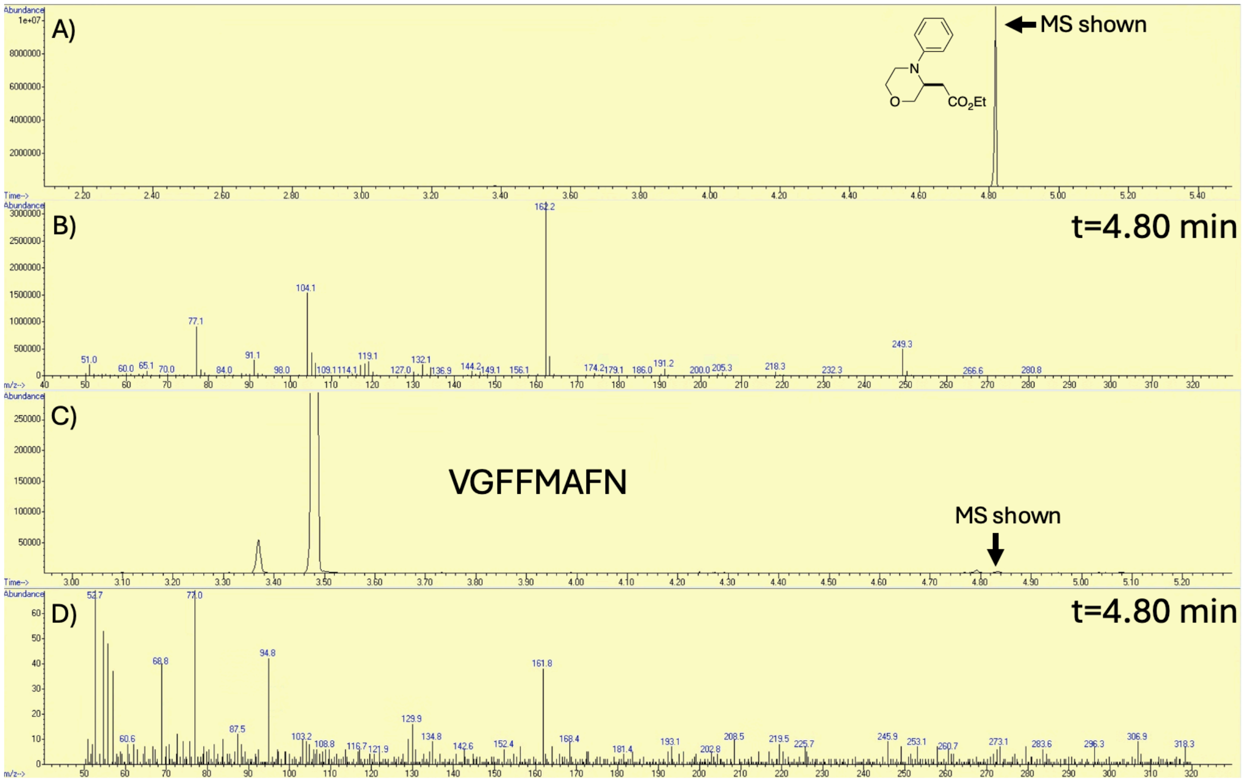


**Figure S49.** **(A)** GC-MS trace for an authentic standard of **C14-p**. **(B)** Selected mass spectrum from the authentic standard peak (t = 4.80 min), showing representative mass peaks for the alkylation product **C14-p**. **(C)** GC-MS trace for reaction **C14** run with variant VFGGMAFN. **C14-p** can be observed in trace quantities. **(D)** Selected mass spectrum from the observed trace product (t = 4.80 min) showing the characteristic m/z = 162 mass for **C14-p**.

#### Reaction C15:

**Scheme S15.** Reaction conditions for reaction **C15** and expected products.

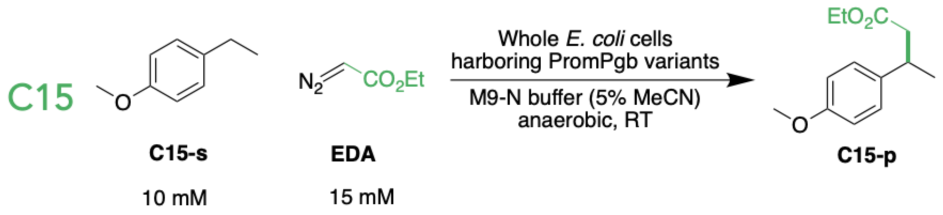


For library screening, reaction **C15** was analyzed by GC-MS. Authentic standards for this reaction were purchased from a commercial source. Reaction **C15** was tested with round 1 and round 2 of predictions. In the course of collecting data with round 1 predictions for model updating, a pooled GC-MS analysis was used for reaction **C15**, yielding qualitative reaction data. Reaction extracts from reactions **C13**, **C14**, **C15**, **C16**, and **C17** were pooled and analyzed by GC-MS. As a result, low-performing variants may have been over diluted and analytes may have been brought below the limit of detection. For model updating, data from this reaction were treated as binary, where a value of 0 denoted no reaction, and a value of 1 indicated detectable product formation. No variants were found to have activity for reaction **C15** for model updating in this work. Later, reaction **C15** was evaluated with round 1 and round 2 of predictions using an un-pooled GC-MS strategy, after which there were still no variants found to catalyze this transformation.

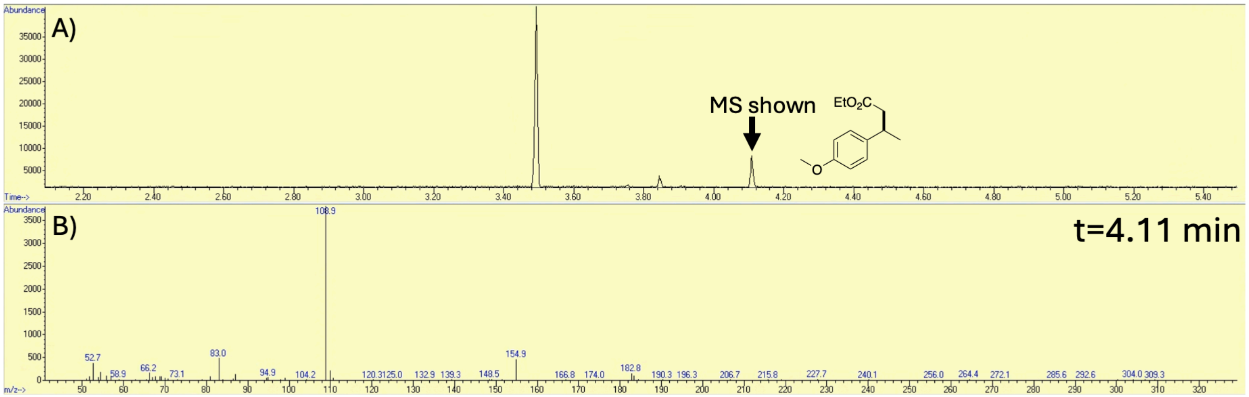

**Figure S50. (A)** GC-MS trace for an authentic standard of **C15-p**. **(B)** Selected mass spectrum from the authentic standard peak (t = 4.11 min), showing representative mass peaks for the alkylation product **C15-p**.

#### Reaction C16:

**Scheme S16.** Reaction conditions for reaction **C16** and expected products.

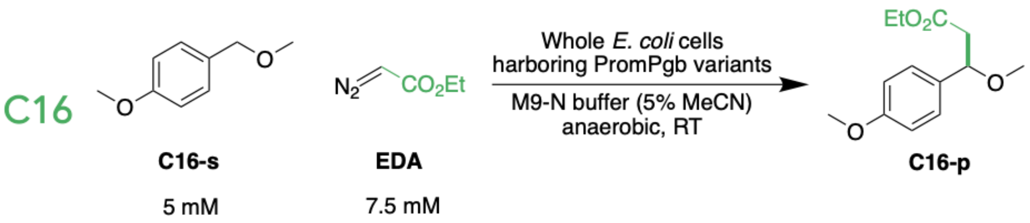


For library screening, reaction **C16** was analyzed by GC-MS. Authentic standards for this reaction were purchased from a commercial source. Reaction **C16** was tested with round 1 and round 2 of predictions. PromPgb was incapable of catalyzing reaction **C16** under these conditions. In the course of collecting data with round 1 predictions for model updating, a pooled GC-MS analysis was used for reaction **C16**, yielding qualitative reaction data. Reaction extracts from reactions **C13**, **C14**, **C15**, **C16**, and **C17** were pooled and analyzed by GC-MS. As a result, low-performing variants may have been over diluted and analytes may have been brought below the limit of detection. For model updating, data from this reaction were treated as binary, where a value of 0 denoted no reaction, and a value of 1 indicated detectable product formation. No variants were found to have activity for reaction **C16** for model updating in this work. Later, reaction **C16** was evaluated with round 1 and round 2 of predictions using an un-pooled GC-MS strategy, uncovering 54 variants which could perform this transformation.

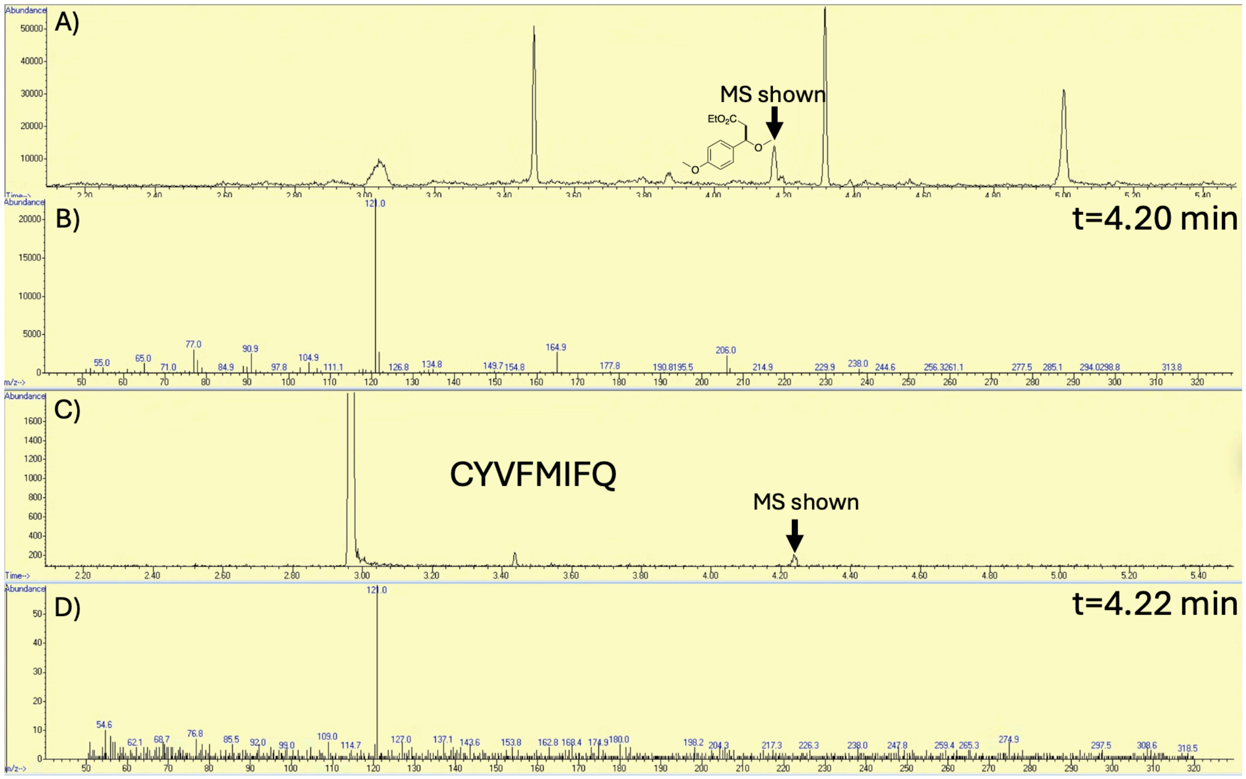


**Figure S51.** **(A)** GC-MS trace for an authentic standard of **C16-p**. **(B)** Selected mass spectrum from the authentic standard peak (t = 4.20 min), showing representative mass peaks for the alkylation product **C16-p**. **(C)** GC-MS trace for reaction **C16** run with variant CYVFMIFQ. **C16-p** can be observed in trace quantities. **(D)** Selected mass spectrum from the observed trace product (t = 4.22 min) showing the characteristic m/z = 121 mass for **C16-p**.

**Reaction C17:**

**Scheme S17.** Reaction conditions for reaction **C17** and expected products.

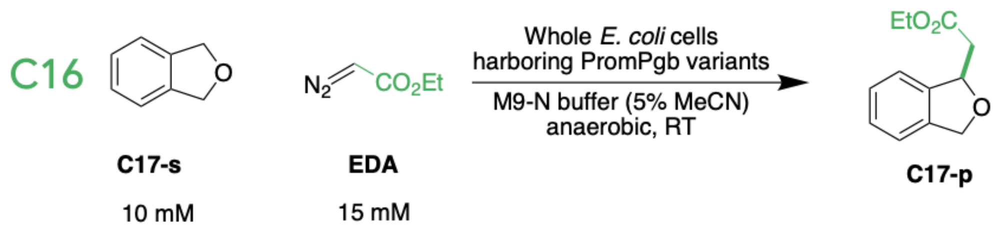


For library screening, reaction **C17** was analyzed by GC-MS. Authentic standards for this reaction were synthesized and characterized in our laboratory in a previous investigation. Full characterization data are reported in reference 11. In the course of collecting data with round 1 predictions for model updating, a pooled GC-MS analysis was used for reaction **C17**, yielding qualitative reaction data. Reaction extracts from reactions **C13**, **C14**, **C15**, **C16**, and **C17** were pooled and analyzed by GC-MS. As a result, low-performing variants may have been over diluted and analytes may have been brought below the limit of detection. For model updating, data from this reaction were treated as binary, where a value of 0 denoted no reaction, and a value of 1 indicated detectable product formation. No variants were found to have activity for reaction **C17** for model updating in this work. Later, reaction **C17** was evaluated with round 1 and round 2 of predictions using an un-pooled GC-MS strategy, after which there were still no variants found to catalyze this transformation.

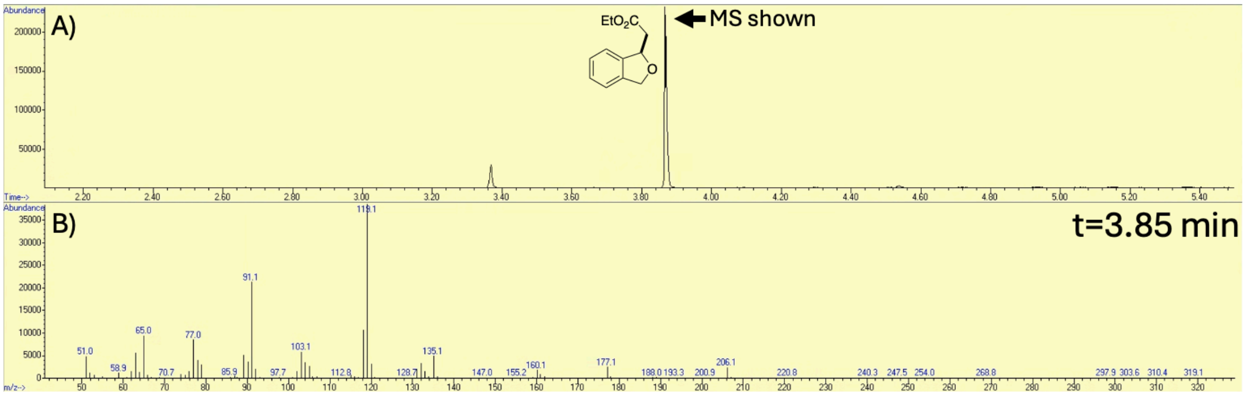


**Figure S52. (A)** GC-MS trace for an authentic standard of **C17-p**. **(B)** Selected mass spectrum from the authentic standard peak (t = 3.85 min), showing representative mass peaks for the alkylation product **C17-p**.

#### Reaction N1:

**Scheme S18.** Reaction conditions for reaction **N1** and expected products.

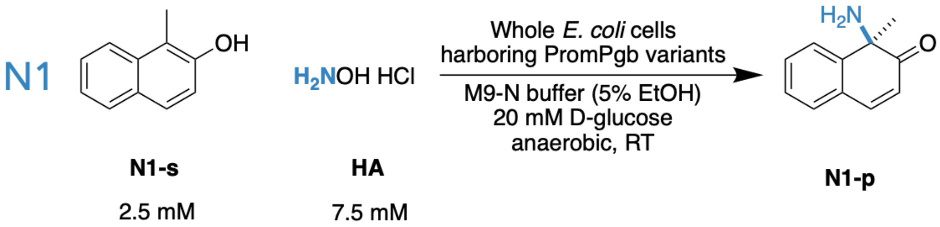


For library screening, reaction **N1** was analyzed by LC-MS. The authentic standard of **N1-p** was kindly provided by Dr. Kathleen M. Sicinski, who previously synthesized and characterized this compound in our laboratory. Full characterization data are reported in reference 12. Reaction **N1** was tested with the initial training library, round 1 of predictions, and round 2 of predictions.

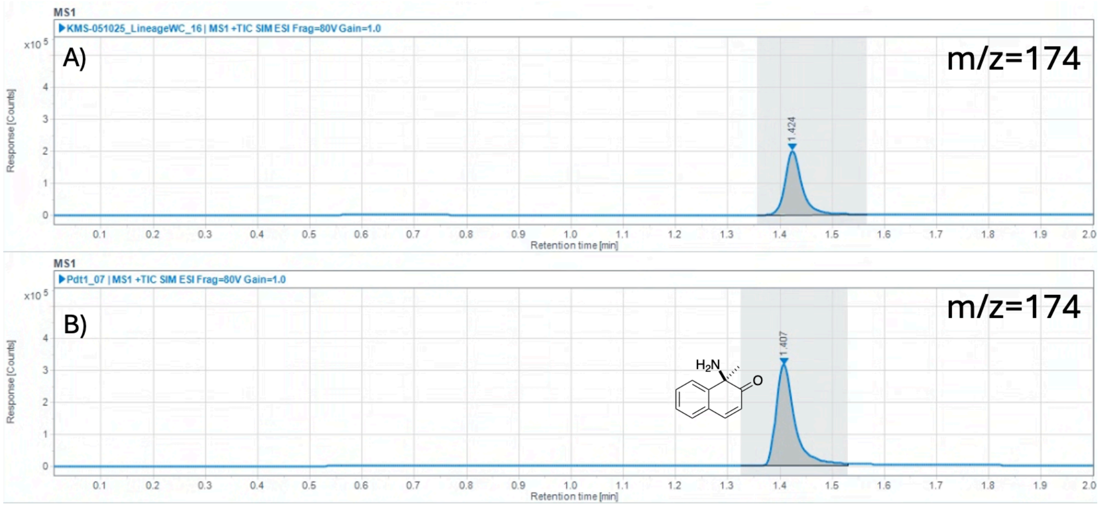


**Figure S53.** **(A)** Representative LC-MS trace for reaction **N1** with PromPgb. **(B)** LC-MS trace for a sample of the authentic standard of **N1-p**. The Selective Ion Monitoring channel for the [M+H]+ ion of **N1-p** (m/z = 174) is shown.

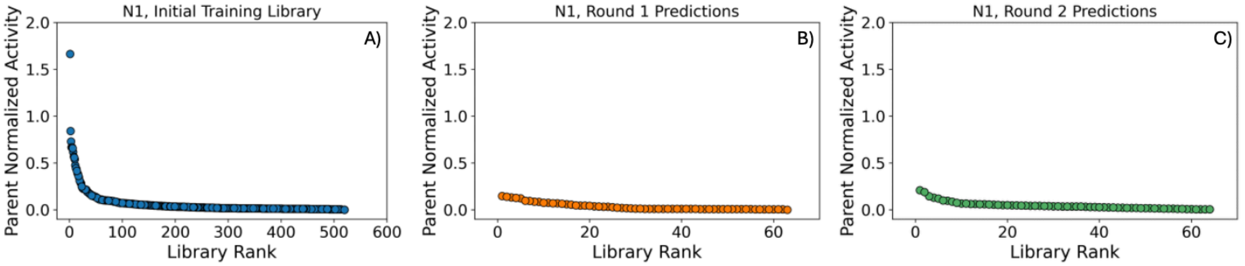


**Figure S54.** Retention of function plots for reaction **N1** activity for variants in the **(A)** initial training library, **(B)** the first round of predictions, and **(C)** the second round of predictions. Parent normalized activities are computed from the total formation of dearomatization product for each variant relative to PromPgb.

#### Reaction N2:

**Scheme S19.** Reaction conditions for reaction **N2** and expected products.

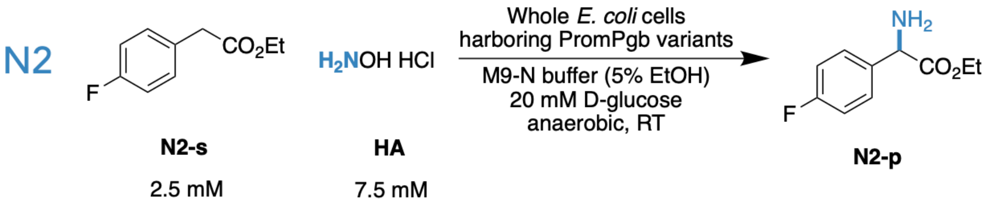


For library screening, reaction **N2** was analyzed by LC-MS. The authentic standard of **N2-p** was kindly provided by Dr. Edwin Alfonzo, who previously synthesized and characterized this compound in our laboratory. Full characterization data are reported in reference 13. Reaction **N2** was tested with the initial training library, round 1 of predictions, and round 2 of predictions. PromPgb was incapable of forming **N2-p** under these conditions.

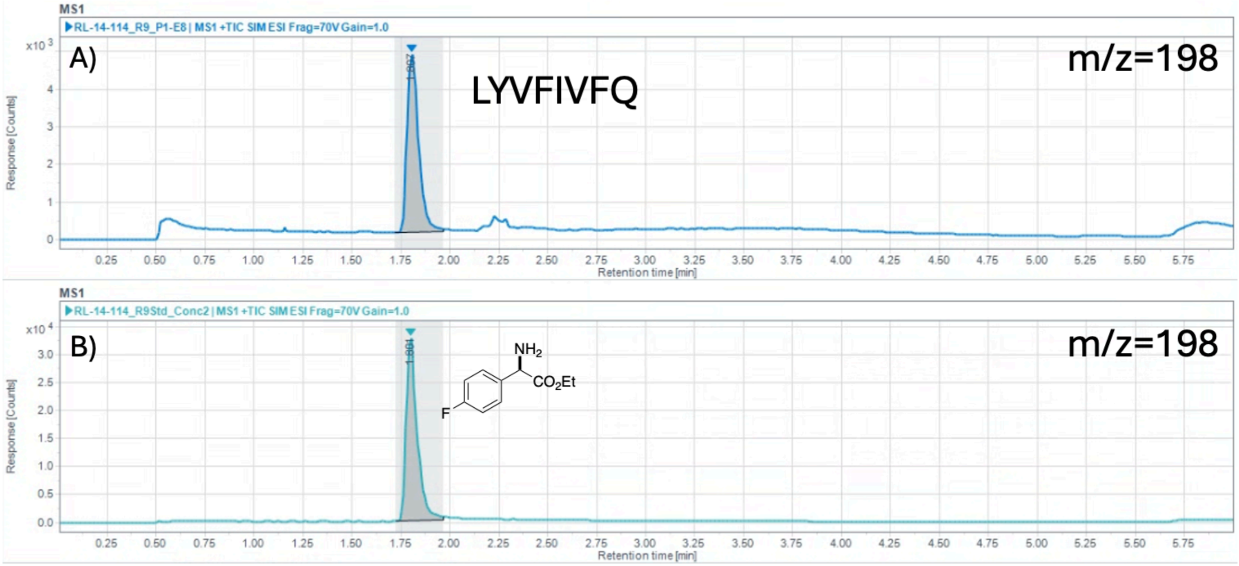


**Figure S55. (A)** LC-MS trace for reaction **N2** with variant LYVFIVFQ from the first round of predictions. **N2-p** formation is observed. **(B)** LC-MS trace for a sample of the authentic standard of **N2-p**. The Selective Ion Monitoring (SIM) channel for the [M+H]+ ion of **N2-p** (m/z = 198) is shown.

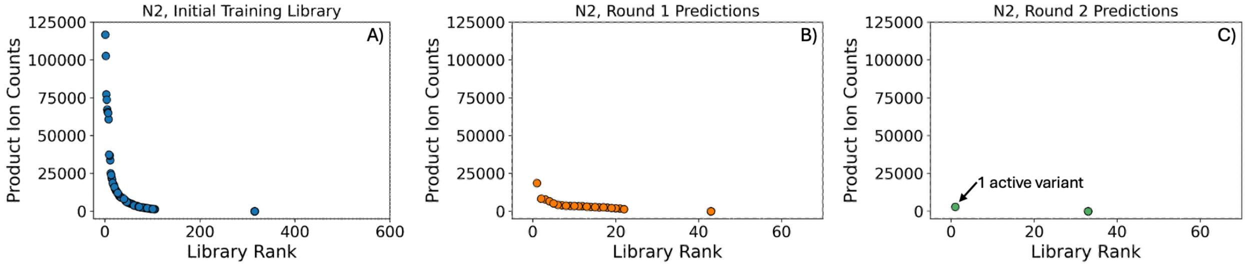


**Figure S56.** Retention of function plots for reaction **N2** activity for variants in the **(A)** initial training library, **(B)** the first round of predictions, and **(C)** the second round of predictions. Variants are ranked by measured ion counts corresponding to product **N2-p** in the SIM channel for the [M+H]+ ion (m/z = 198).

#### Reaction N3:

**Scheme S20.** Reaction conditions for reaction **N3** and expected products.

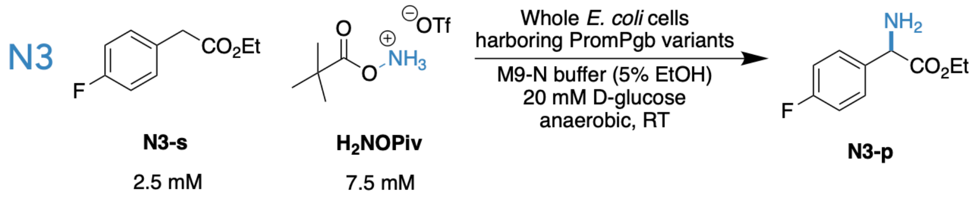


For library screening, reaction **N3** was analyzed by LC-MS. Authentic standards for this reaction were synthesized and characterized in our laboratory. Reaction **N3** was tested with the initial training library, round 1 of predictions, and round 2 of predictions.

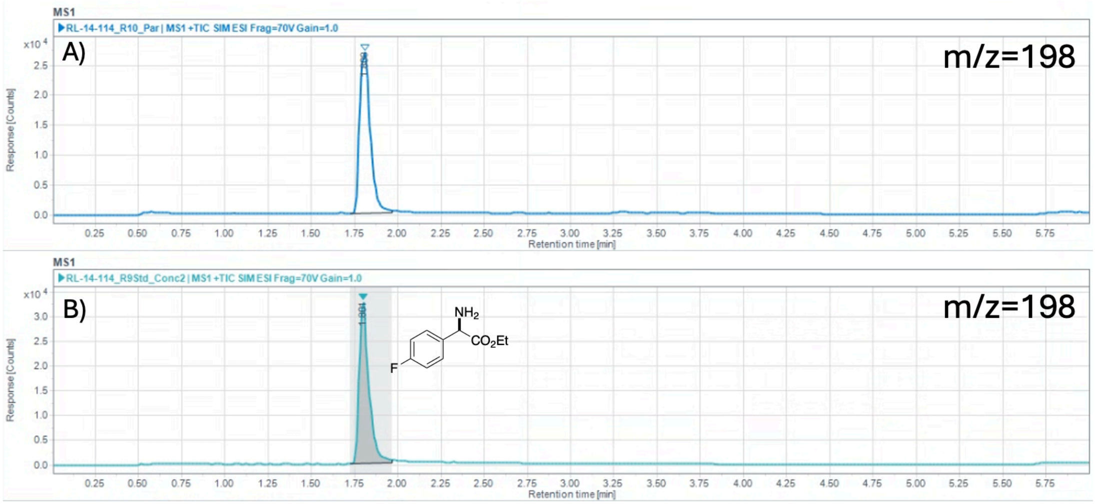


**Figure S57. (A)** Representative LC-MS trace for reaction **N3** with PromPgb. **(B)** LC-MS trace for a sample of the authentic standard of **N3-p**. The Selective Ion Monitoring (SIM) channel for the [M+H]+ ion of **N3-p** (m/z = 198) is shown.

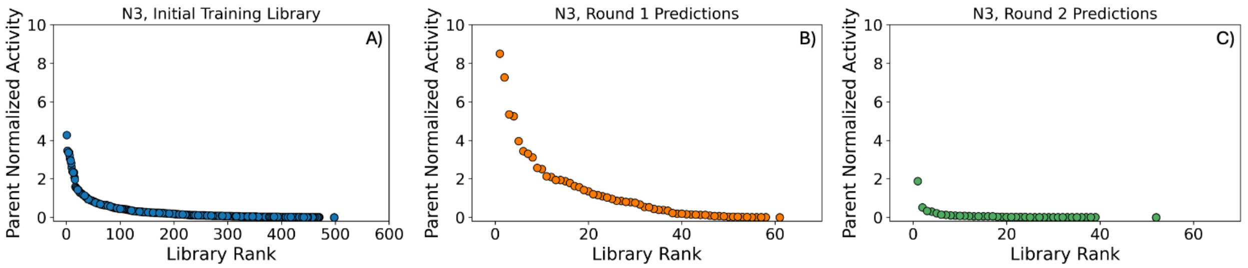


**Figure S58.** Retention of function plots for reaction **N3** activity for variants in the **(A)** initial training library, **(B)** the first round of predictions, and **(C)** the second round of predictions. Parent normalized activities are computed from the total formation of C–H amination product for each variant relative to PromPgb.

#### Reaction N4:

**Scheme S21.** Reaction conditions for reaction **N4** and expected products.

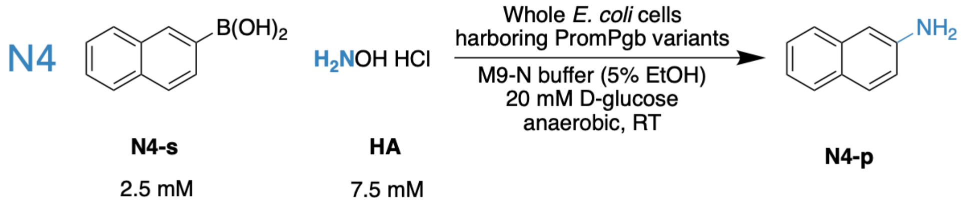


For library screening, reaction **N4** was analyzed by LC-MS. Authentic standards for this reaction were purchased from a commercial source. Reaction **N4** was tested with the initial training library, round 1 of predictions, and round 2 of predictions.

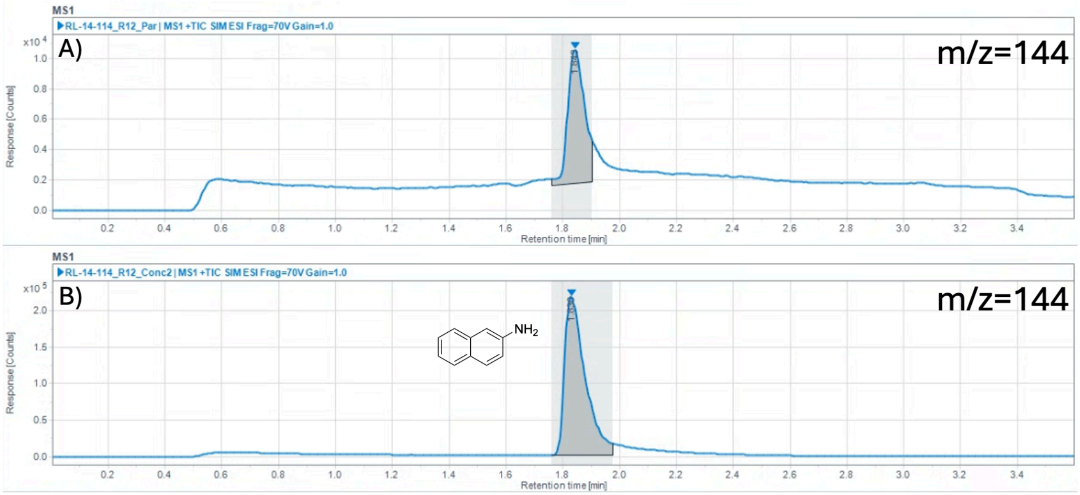


**Figure S59. (A)** Representative LC-MS trace for reaction **N4** with PromPgb. **(B)** LC-MS trace for a sample of the authentic standard of **N4-p**. The Selective Ion Monitoring (SIM) channel for the [M+H]+ ion of **N4-p** (m/z = 144) is shown.

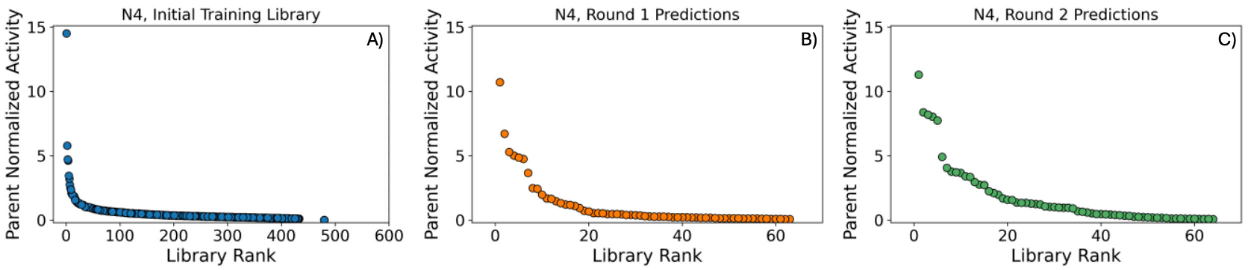


**Figure S60.** Retention of function plots for reaction **N4** activity for variants in the **(A)** initial training library, **(B)** the first round of predictions, and **(C)** the second round of predictions. Parent normalized activities are computed from the total formation of C–H amination product for each variant relative to PromPgb.

#### Reaction N5:

**Scheme S22.** Reaction conditions for reaction **N5** and expected products.

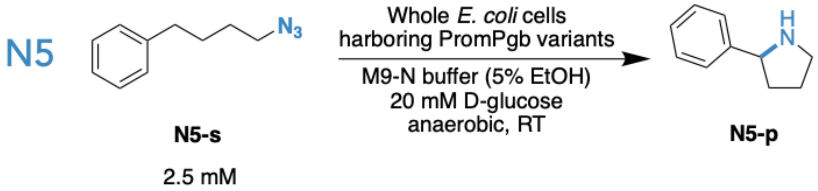


For library screening, reaction **N5** was analyzed by LC-MS. Authentic standards for this reaction were purchased from a commercial source. Reaction **N5** was tested with round 1 and round 2 of predictions.

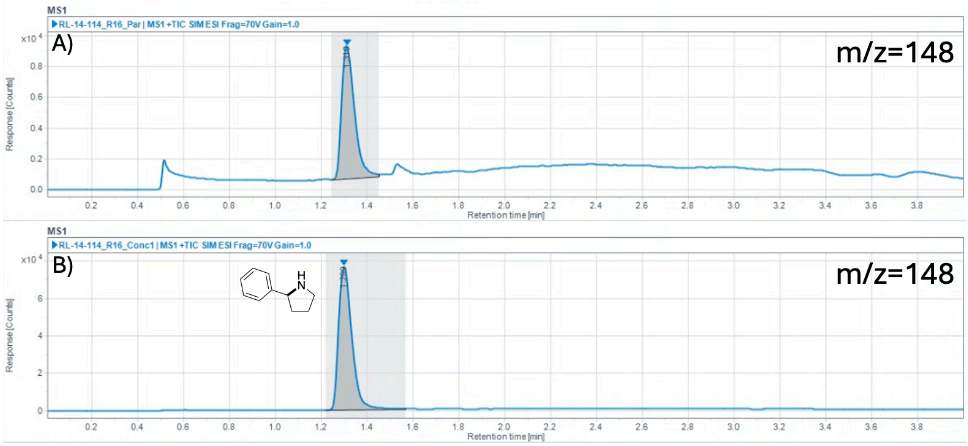


**Figure S61. (A)** Representative LC-MS trace for reaction **N5** with PromPgb. **(B)** LC-MS trace for a sample of the authentic standard of **N5-p**. The Selective Ion Monitoring (SIM) channel for the [M+H]+ ion of **N5-p** (m/z = 148) is shown.

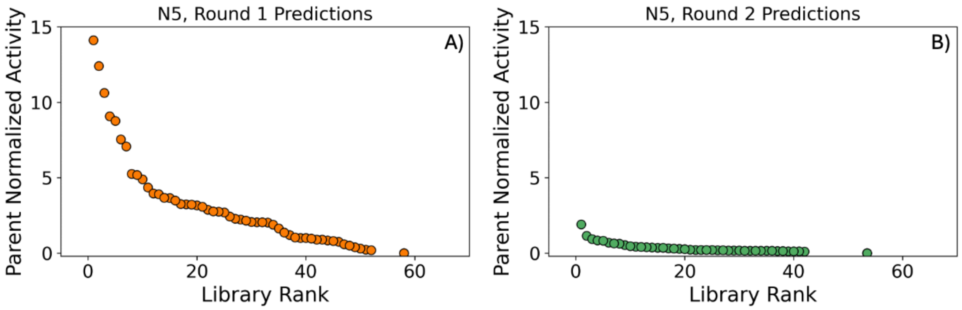


**Figure S62.** Retention of function plots for reaction **N5** activity for variants in the **(A)** first round of predictions and **(B)** the second round of predictions. Parent normalized activities are computed from the total formation of intramolecular reaction product for each variant relative to PromPgb.

#### Reaction N6:

**Scheme S23.** Reaction conditions for reaction **N6** and expected products.

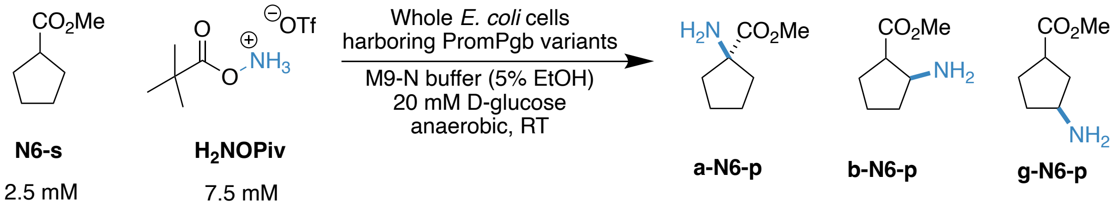


For library screening, reaction **N6** was analyzed by LC-MS. Authentic standards for the possible $\alpha$-, $\beta$-, $\gamma$-amination products for this reaction were purchased from a commercial source. Reaction **N6** was tested with round 1 and round 2 of predictions. PromPgb was incapable of catalyzing reaction **N6** under these conditions. In library screening, five variants were found in the first round of predictions which could generate trace quantities of **N6-p** isomers and no variants were found in the second round of predictions. For model updating, data from this reaction were treated as binary, where a value of 0 denoted no reaction, and a value of 1 indicated detectable product formation.

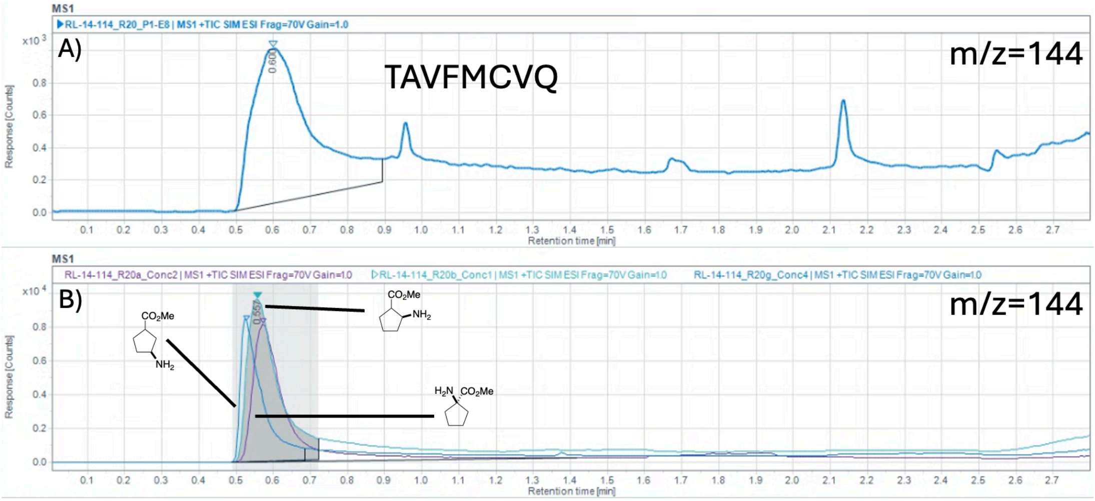


**Figure S63. (A)** LC-MS trace for reaction **N6** with variant TAVFMCVQ from the first round of predictions. The three possible regioisomers of **N6-p** formation could not be well-separated on the LC-MS column and so were treated as a single eluting peak. **(B)** LC-MS traces for samples of the authentic standards of **a-N6-p**, **b-N6-p**, and **g-N6-p**. The Selective Ion Monitoring (SIM) channel for the [M+H]+ ion of **N6-p** (m/z = 144) is shown.

#### Reaction N7:

**Scheme S24.** Reaction conditions for reaction **N7** and expected products.

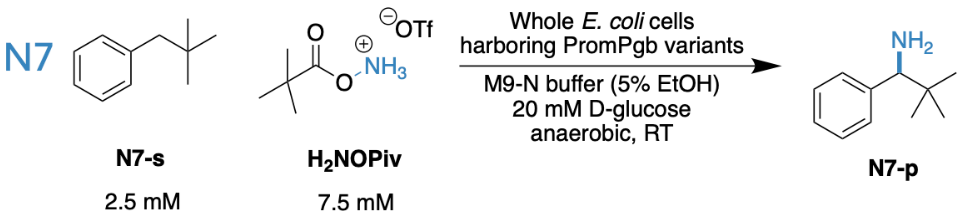


For library screening, reaction **N7** was analyzed by LC-MS. Authentic standards for this reaction were purchased from a commercial source. Reaction **N7** was tested with round 1 and round 2 of predictions. PromPgb was incapable of catalyzing reaction **N7** under these conditions. In library screening, three variants were found in the first round of predictions which could generate trace quantities of **N7-p** isomers and no variants were found in the second round of predictions. For model updating, data from this reaction were treated as binary, where a value of 0 denoted no reaction, and a value of 1 indicated detectable product formation.

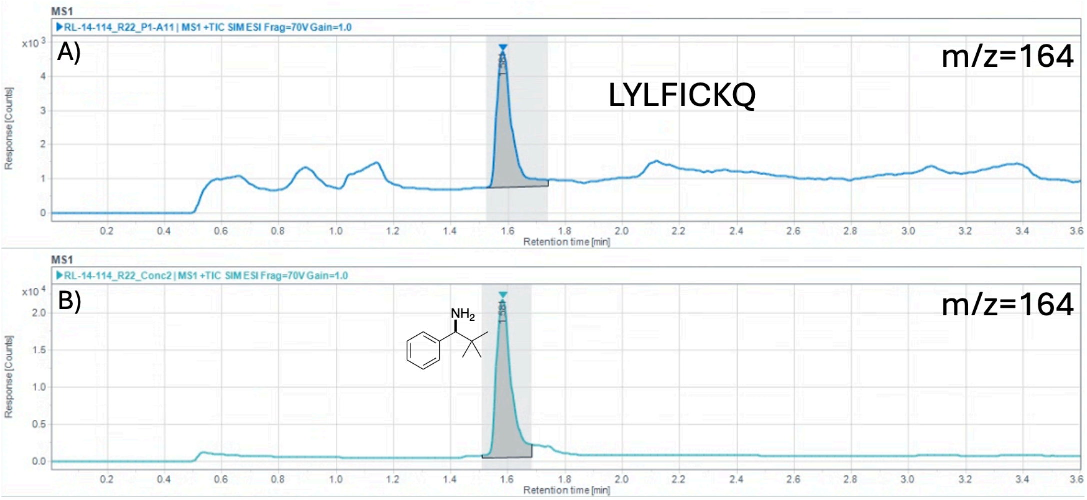


**Figure S64. (A)** Representative LC-MS trace for reaction **N7** with variant LYLFICKQ from the first round of predictions. **(B)** LC-MS trace for a sample of the authentic standard of **N7-p**. The Selective Ion Monitoring (SIM) channel for the [M+H]+ ion of **N7-p** (m/z = 164) is shown.

#### Reaction N8:

**Scheme S25.** Reaction conditions for reaction **N8** and expected products.

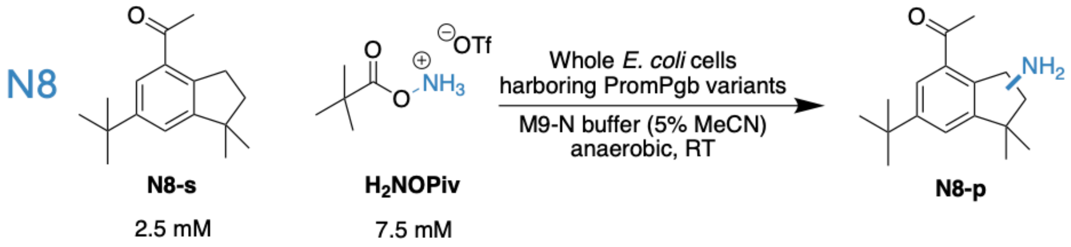


For library screening, reaction **N8** was analyzed by LC-MS. For reaction **N8**, the exact product structure was not determined; activity was quantified based on an LC–MS peak corresponding to the expected $[M+H]^+$ ion. Taken together, the data shown in **Figure S65** support the occurrence of a protoglobin-catalyzed nitrene transfer reaction with **N8-s**. The product peak observed at t = 1.73 min in panels **Figure S65A** and **Figure S65B** is detected only in reactions containing whole cells harboring protoglobin variants, and its abundance depends on the identity of the protoglobin. This peak is not observed in the absence of whole cells (**Figure S65C**) or in the presence of whole cells expressing TrpB, a PLP-dependent enzyme serving as our negative control (**Figure S65D**).

Reaction **N8** was tested with round 1 of predictions.

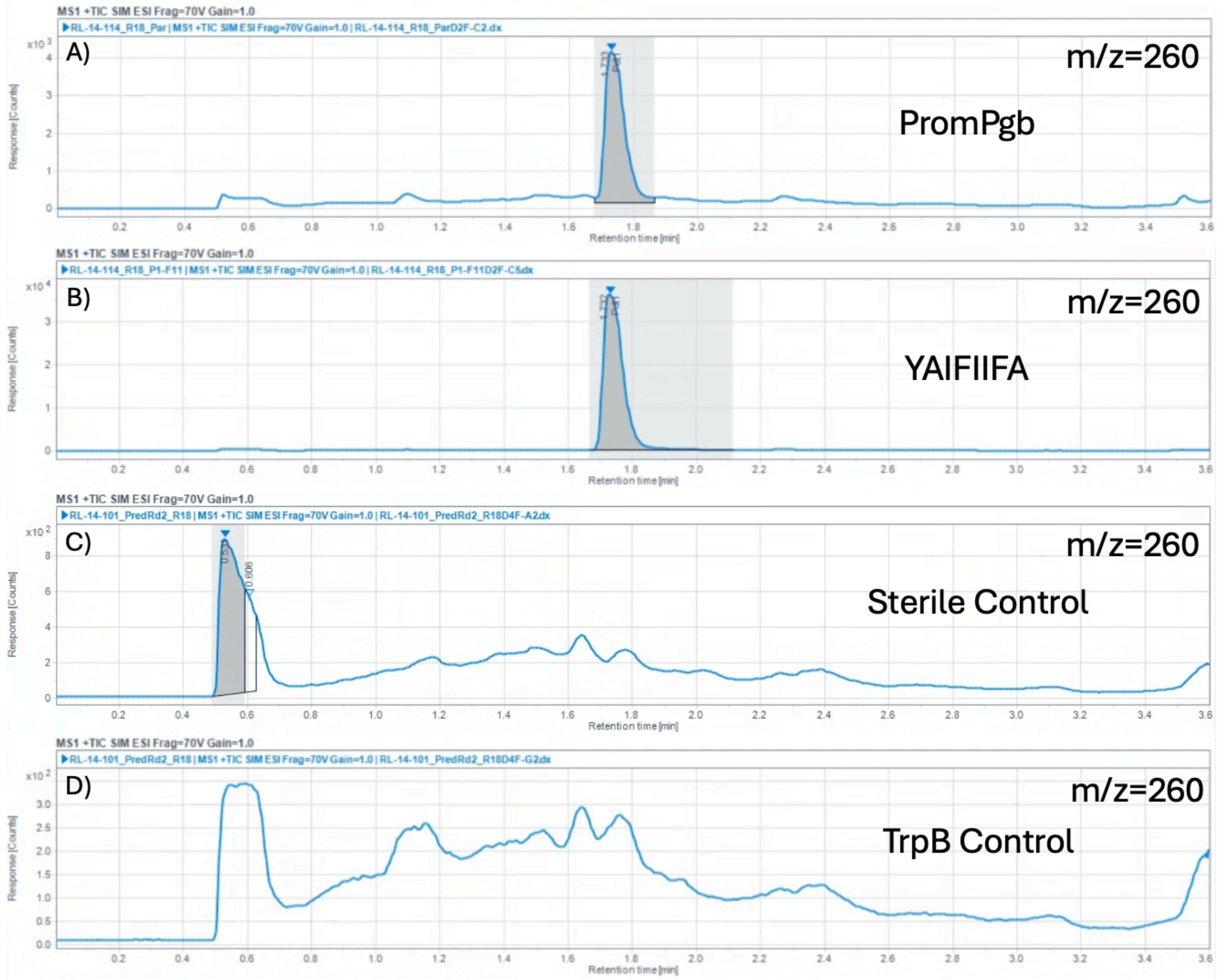


**Figure S65. (A)** Representative LC-MS trace for reaction **N8** with PromPgb. **(B)** LC-MS trace for reaction **N8** with the high-performing variant YAIFIIFA. **(C)** Representative LC-MS trace for reaction **N8** in the absence of any whole cells. All reaction conditions are otherwise kept the same. **(D)** Representative LC-MS trace for reaction **N8** in the presence of whole cells expressing The Selective Ion Monitoring (SIM) channel for the $[M+H]^+$ ion of **N8-p** (m/z = 260) is shown for all chromatograms.

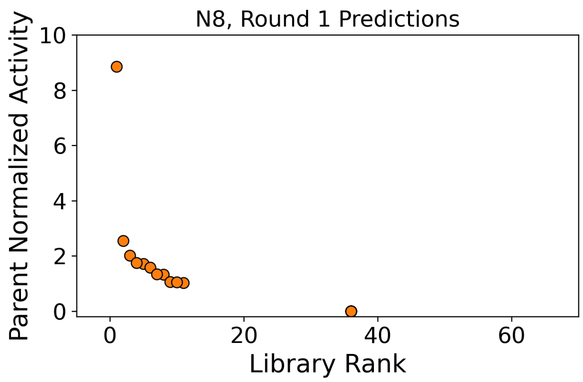


**Figure S66.** Retention of function plot for reaction **N8** activity for variants in the first round of predictions.

#### Reaction N9:

**Scheme S26.** Reaction conditions for reaction **N9** and expected products.

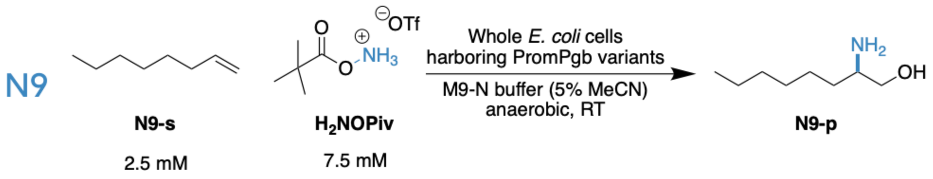


For library screening, reaction **N9** was analyzed by LC-MS. Authentic standards for this reaction were synthesized and characterized in our laboratory. Reaction **N9** was tested with round 1 and round 2 of predictions. During Round 1 data collection, four variants were initially assigned activity for reaction **N9**, and these measurements were incorporated into the model update. Subsequent reanalysis of these samples did not confirm product formation under the reported conditions. PromPgb was not capable of performing this reaction, and upon further screening no variants were found in either round of predictions which were capable of performing this transformation with detectable yield.

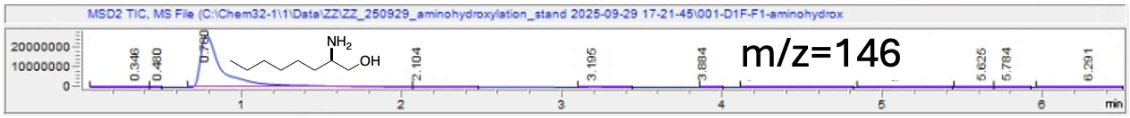


**Figure S67.** LC-MS trace for a sample of the authentic standard of **N9-p**. The Selective Ion Monitoring channel for the $[M+H]^+$ ion of **N9-p** (m/z = 146) is shown.

### Computed Summary Statistics

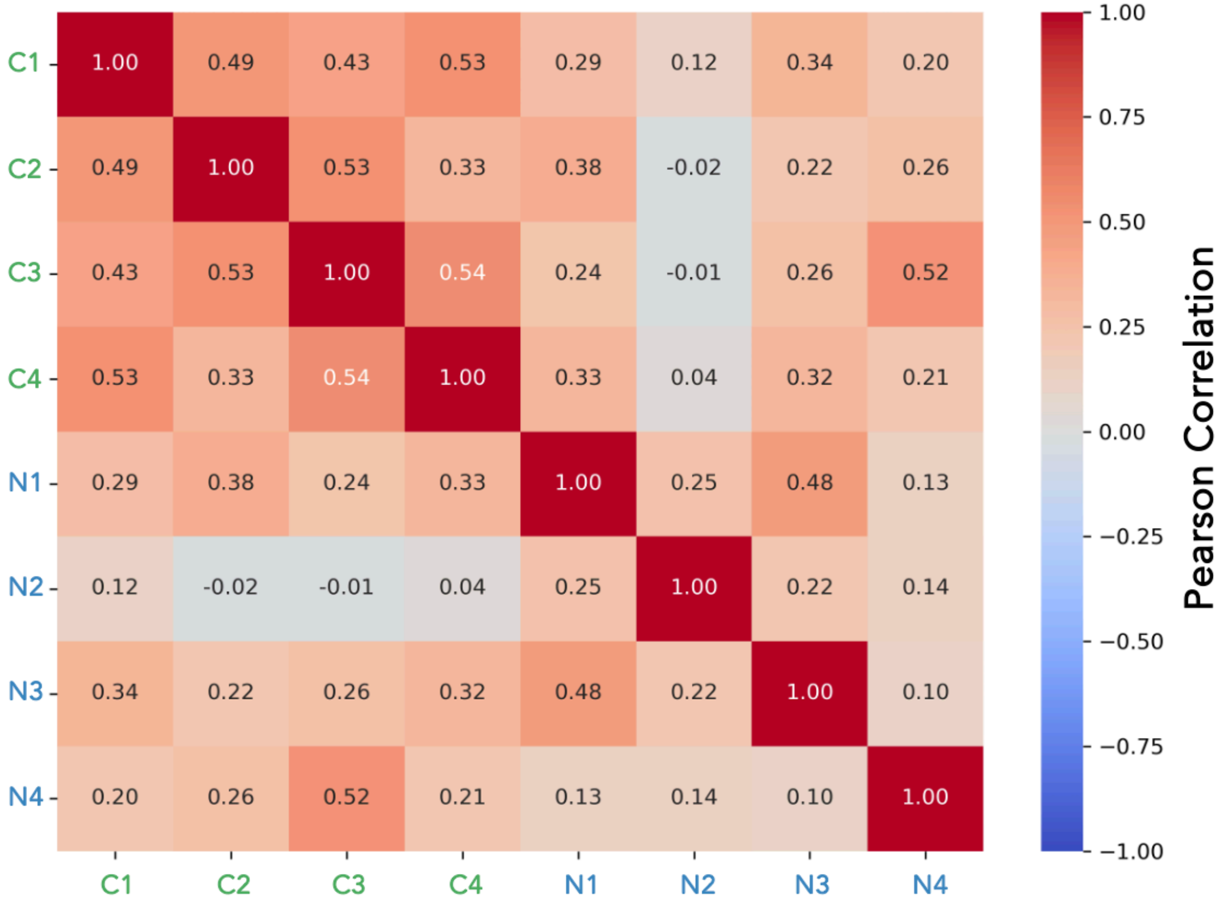


**Figure S68.** Computed pairwise Pearson correlation coefficients (r) between the fitness values for each of the eight ITRs, calculated using data from variants in the initial training library. Correlations were computed across all variants for which measurable activity was observed in both reactions. Positive values indicate that mutations tending to enhance activity in one reaction also enhance activity in the other, whereas low or near-zero correlations suggest reaction-specific mutational effects.

### Machine Learning Details

**Data Processing.** At the start of each round of the computational pipeline, sequences were paired with a set of respective experimental yields for carbene and nitrene reactions. The reaction yields were processed and normalized to the parent (starting) sequence, such that for a given reaction, 1.0 corresponded to the same yield as parent. For variants displaying activity for reaction **N2**, fold-improvements in activity are normalized to the yield of parent for reaction **N3**, which shares the same product. Outliers with very high yield (i.e., greater than 4 times that of parent) were rounded down to 4.0, to encourage promiscuity rather than optimizing for a specific reaction. For reactions with trace activity that were not detected in parent, a variant was assigned 1 if could perform that reaction and 0 otherwise. The two fitness objectives used as predictive outputs for ML model training were the mean fitness values of all carbene and nitrene reactions, respectively, thus weighting improvements to carbene reactions relatively equally to improvements to nitrene reactions.

**Surrogate Model Training.** Most Bayesian optimization algorithms consist of two main components: (1) a probabilistic surrogate model of the objective function and (2) an acquisition function. The surrogate model predicts the objective function values at unobserved inputs, while the acquisition function quantifies the potential benefit of evaluating any given batch of inputs based on these predictions. In each iteration of the Bayesian optimization loop, a new batch of inputs is selected by maximizing the acquisition function. After evaluating the objective function at these new inputs, the surrogate model is updated, and the process repeats. Let $\boldsymbol{X}$ denote the input space (i.e., the space of feasible protein sequences) and let $f: \boldsymbol{X} \rightarrow \boldsymbol{R}$ denote the objective function (i.e., the metric we wish to optimize). In this work, we used deep ensembles as the surrogate model, which is based on their high performance and calibrated uncertainty seen in past work.[8]

Deep ensembles are constructed by training identical deep neural network architectures multiple times, each with different random initializations of the weight parameters. Here, we train the deep ensembles with bootstrapping; each ensemble consists of 20 models where 90% of the total training data is randomly seen during training, and each model is a multilayer perceptron with two hidden layers and a hidden dimension of 50. The input to each model is a one-hot encoding of the 8 residues being studied, with dimension 160, and the outputs are two values, corresponding to the predictions for the two normalized carbene and nitrene reaction yields, described above. We surmised that performing multi-task training by sharing weights would improve regularization and reduce overfitting in the model, as the two distinct objectives are correlated. For each model, we performed training with a batch size of 128 and a learning rate of 0.001 for 100 epochs. Model training time was less than one minute on a single GPU. These independently trained networks were then collectively used as if they were samples from a Bayesian posterior distribution over the objective function $f$.

**Sample Acquisition.** We used the expected improvement (EI) acquisition function, which is given by $\alpha_n(x) = E_n[\{f(x) - f_n^*\}^+]$, where $f_n^* = \max_{i=1,\ldots,n} f(x_i)$ and the expectation is computed with respect to the posterior distribution given $\mathcal{D}_n$.[15] For Gaussian posterior distributions and noise-free observations (where $f_n^*$ is a constant rather than a random variable), the EI can be expressed in a closed form using the posterior mean and variance. In scenarios where these conditions do not hold, computing the EI often requires approximate calculation, typically through Monte Carlo sampling techniques.

In this case, we extended expected improvement to expected hypervolume improvement (eHVI), which corresponds to expected improvement in the pareto front "area" across the two objectives described above and is commonly used in multi-objective settings. We acquired a batch of new samples to maximize eHVI following a procedure similar to that used in MORBO[16] because this approach enables better scalability to large batch sizes, by exploiting the submodularity of the acquisition function. Thus, efficient approximation can be achieved through a greedy optimization strategy, selecting each input in the batch sequentially. The main difference between our approach and the approach used in MORBO is that we used Thompson sampling with a frequentist ensemble of neural networks rather than a Gaussian process as the surrogate function. We used a reference fitness value of 0 for both objectives in the expected improvement acquisition function. The algorithm we used to obtain a batch of 96 proposed samples was implemented using the botorch python package:[17]

1. Sample a function from the ensemble of supervised models (Thompson-style sampling)
2. Evaluate the predicted increase in hypervolume (across two objectives) for all points in the design space, using the sampled function
3. Query the next point in the batch as the point that increases the hypervolume the most
4. Repeat from step one until the entire batch is sampled, but the increase in area is evaluated with respect to expected gains from previously queried points

To speed up calculations, we only evaluated the acquisition function for a subset of the design space by only considering amino acid substitutions that corresponded to variants with some activity in the initial round of data collection, reducing the design space to about 180 million variants, based on a somewhat arbitrary cutoff. Evaluating the acquisition function 96 times took around 3 hours on a single H100 GPU.

### Quantitative Analysis of Select PromPgb Variants

Reactions **C1**, **C8**, **C13**, and **N5** were all validated under controlled conditions with small-scale protein expression of PromPgb and top-performing variants for each of these reactions.

**Protocols for Small-Scale Reaction Setup in GC Vials:**

*Small-Scale Protein Expression*

Single colonies from LB-Agar plates were picked using a sterile pipette tip and were used to inoculate 6 mL of LB-Amp in a 15-mL culture tube. Cultures were incubated at 37 ºC with shaking at 220 rpm overnight in an Innova 4000 shaking incubator. A 1-mL aliquot of each of these overnight cultures was used to inoculate 50 mL of Terrific Broth with 100 µg/mL of ampicillin (TB-Amp) (0.5% v/v starter culture in expression culture) in 125-mL unbaffled Erlenmeyer flasks. The expression cultures were incubated at 37 ºC and 220 rpm for 2.5 hours in an Innova 42 shaking incubator, at which point they are moved to room temperature for 25 minutes. Protein expression was then induced by direct addition of 50 µL of stock solutions containing 500 mM isopropyl-β-D-thiogalactoside (IPTG) and 1.0 M 5-aminolevulinic acid (ALA) such that the final concentrations were 0.5 mM and 1.0 mM, respectively. The cultures were shaken at 22 ºC and 140 rpm for 16–18 hours in an Innova 42 shaker.

*Small-Scale Biocatalytic Carbene Transfer Reaction Preparation*

The corresponding 50-mL expression cultures were pelleted (4,000 × g for 15 minutes at 4 ºC) and resuspended in 5 mL of M9-N buffer. The optical density at 600 nm ($OD_{600}$) of this suspension was measured and adjusted to $OD_{600} = 31.5$ with the addition of more M9-N buffer. A 380-µL aliquot of the cell suspension was added to 2-mL GC vials (Agilent). These whole-cell suspensions were transferred into a vinyl Coy anaerobic chamber, at which point 10 µL of a solution of the reaction substrate in MeCN followed by 10 µL of a solution of EDA in MeCN were added. The GC vials were tightly capped with screwcaps with a septum and were allowed to shake at room temperature for 16 hours.

Once complete, the reactions were transferred to 1.7-mL Eppendorf tubes and mixed with 600 µL of a 1:1 solution of ethyl acetate:cyclohexane with 1,3,5-trimethoxybenzene as an internal standard (1.0 mM concentration). The layers were vigorously mixed, and the samples were centrifuged (14,000 × g for 10 minutes at RT). Afterwards, an aliquot of the organic layer was subjected to GC analysis.

*Small-Scale Biocatalytic Nitrene Transfer Reaction Preparation*

The corresponding 50-mL expression cultures were pelleted (4,000 × g for 15 minutes at 4 ºC) and resuspended in 5 mL of M9-N buffer containing 20 mM D-glucose. The optical density at 600 nm ($OD_{600}$) of this suspension was measured and adjusted to $OD_{600}$ = 31.5 with the addition of more M9-N buffer. A 370 aliquot of the cell suspension was added to 2-mL GC vials (Agilent). These whole-cell suspensions were transferred into a vinyl Coy anaerobic chamber, at which point 20 µL of a solution of the reaction substrate in EtOH followed by 10 µL of a solution of nitrene precursor in $H_2O$ were added. The GC vials were tightly capped with screwcaps with a septum and were allowed to shake at room temperature for 16 hours.

Once complete, the reactions were transferred to 1.7-mL Eppendorf tubes and mixed with 800 µL of an acetonitrile solution containing papaverine as an internal standard (50 $\mu$M concentration). The layers are vigorously mixed after which the samples were stored at -20 °C for 1 hour. The samples were then centrifuged (14,000 × g for 10 minutes at RT). Afterwards, an aliquot of the resulting clarified solution was subjected to LC-MS analysis.

### Preparation of Calibration Curves for Analytical Yield Determination

All reactions with quantifiable yield were assayed either by gas chromatography equipped with flame ionization detection (GC-FID) or by liquid chromatography equipped with mass spectrometric detection (LC-MS). To quantify yields, calibration curves were prepared from synthesized authentic standards or products isolated from reactions. Calibration curves were constructed by one of two methods.

*GC-FID Calibration Curve Construction – Method A*

Method A was used for products which could be separated and detected by GC-FID. For these compounds, a 10 mM stock of authentic standard or isolated product was prepared in a solution of 1:1 solution of ethyl acetate:cyclohexane with 1,3,5-trimethoxybenzene as an internal standard (1.0 mM concentration). This stock solution was diluted in the same solution of ethyl acetate:cyclohexane solution containing 1.0 mM standard to a range of concentrations within the range of the measured enzymatic product concentration. In instances where two diastereomers could be formed, calibration curve samples were generated for each diastereomer. Samples were analyzed by GC-FID. Calibration curves were then generated by plotting the concentration of the product against the ratio of its signal (P) to the signal of the internal standard (IS).

*LC-MS Calibration Curve Construction – Method B*

Method B was used for products which could be separated and detected by LC-MS. For these compounds, 380 $\mu$L of M9-N buffer were mixed with 20 $\mu$L of the authentic product dissolved in ethanol at several unique concentrations such that the sample contained the product at final concentrations within the range of measured enzymatic formation. This mixture was then processed according to the protocol for working up enzymatic reactions for LC-MS quantification (**Small-Scale Biocatalytic Nitrene Transfer Reaction Preparation**).

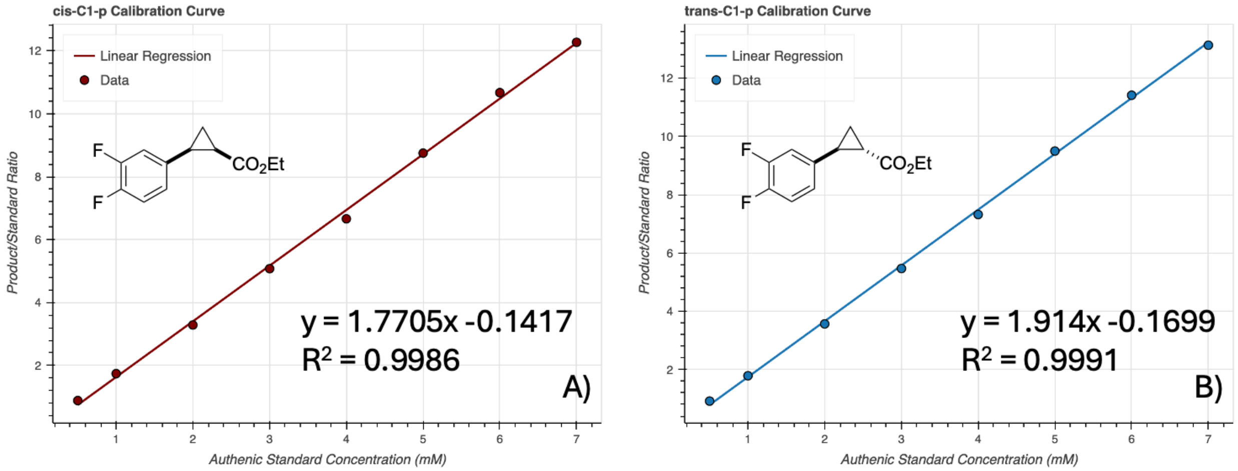


**Figure S69.** Calibration curves for the products of reaction **C1**. **(A)** Achiral GC-FID calibration curve for ***cis*-C1-p**. **(B)** Achiral GC-FID calibration curve for ***trans*-C1-p**. Samples were generated using method A.

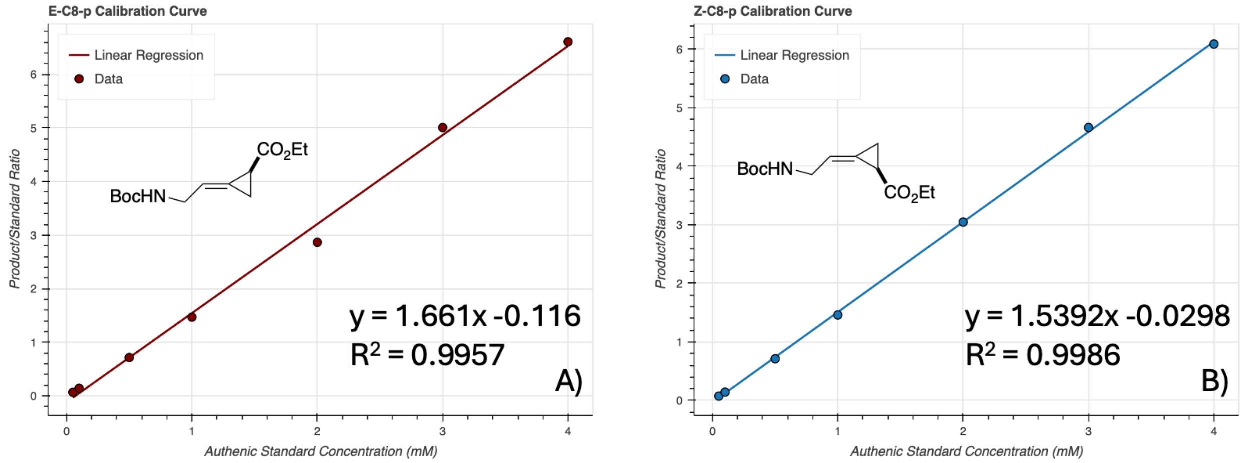


**Figure S70.** Calibration curves for the products of reaction **C8**. **(A)** Achiral GC-FID calibration curve for ***E*-C8-p**. **(B)** Achiral GC-FID calibration curve for ***Z*-C8-p**. Samples were generated using method A.

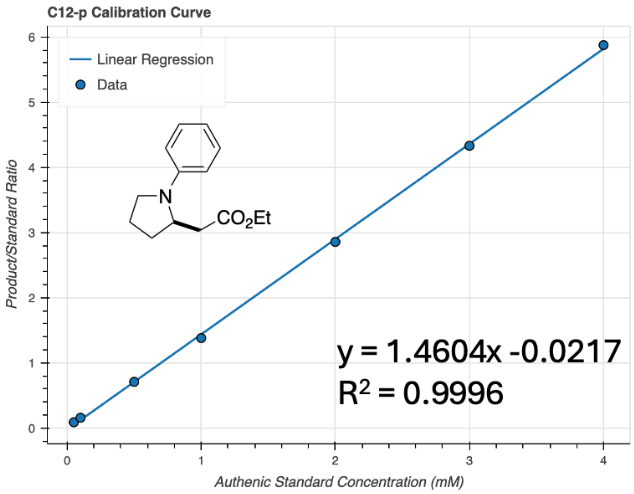


**Figure S71.** Achiral GC-FID calibration curve for **C12-p**. Samples were generated using method A.

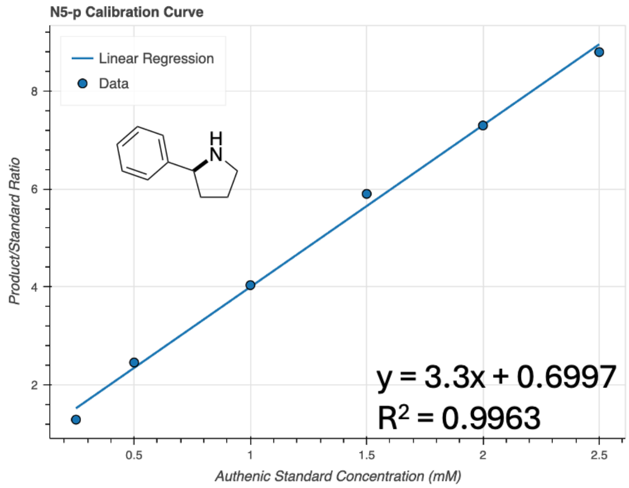


**Figure S72.** Achiral GC-FID calibration curve for **N5-p**. Samples were generated using method B.

### Preparation of Authentic Standards

#### *Synthesis of **cis-C1-p** and **trans-C1-p***

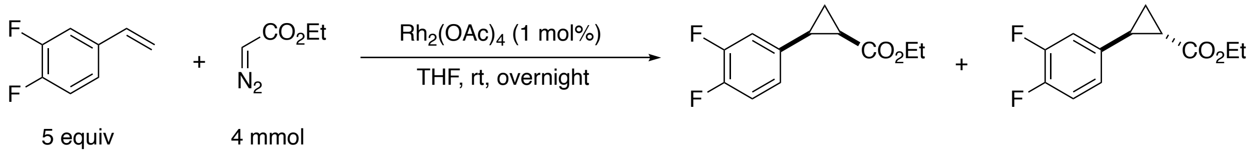


A solution of ethyl diazoacetate (1 equiv, 4 mmol, ≥13 wt.% DCM) in THF (1.2 ml) was added dropwise to a stirred mixture of 1,2-difluoro-4-vinylbenzene (5 equiv, 20 mmol) and rhodium (II) diacetate dimer (1.0 mol %, 0.04 mmol) in dry THF (2 ml) under an argon atmosphere. The reaction mixture was stirred for 16 h, then filtered through a short plug of alumina gel using diethyl ether as an eluent. The filtrate was concentrated *in vacuo*. The resulting residue was purified by column chromatography on silica gel in hexanes/EtOAc to afford the cyclopropanation products as separate diastereomers.

(±)-*cis*-ethyl 2-(3,4-difluorophenyl)cyclopropane-1-carboxylate ***cis-C1-p***

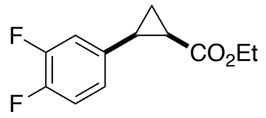


$^1$H NMR (400 MHz, $CDCl_3$) δ 7.12 – 6.93 (m, 3H), 3.93 (q, $J$ = 7.1 Hz, 2H), 2.58 – 2.43 (m, 1H), 2.08 (ddd, $J$ = 9.2, 7.9, 5.6 Hz, 1H), 1.63 (dt, $J$ = 7.4, 5.4 Hz, 1H), 1.35 (ddd, $J$ = 8.7, 7.9, 5.2 Hz, 1H), 1.05 (t, $J$ = 7.1 Hz, 3H).

(±)-*trans*-ethyl 2-(3,4-difluorophenyl)cyclopropane-1-carboxylate ***trans-C1-p***

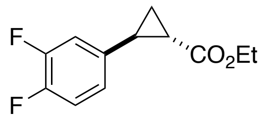


[1]H NMR (400 MHz, $CDCl_3$) δ 7.05 (dt, *J* = 10.2, 8.3 Hz, 1H), 6.93 – 6.79 (m, 2H), 4.17 (q, *J* = 7.3 Hz, 2H), 2.47 (ddd, *J* = 9.2, 6.4, 4.1 Hz, 1H), 1.84 (ddd, *J* = 8.5, 5.4, 4.2 Hz, 1H), 1.59 (ddd, *J* = 9.2, 5.3, 4.7 Hz, 1H), 1.28 (t, *J* = 7.2 Hz, 3H), 1.26 – 1.21 (m, 1H).

#### *Synthesis of* ***cis-C3-p*** *and* ***trans-C3-p***

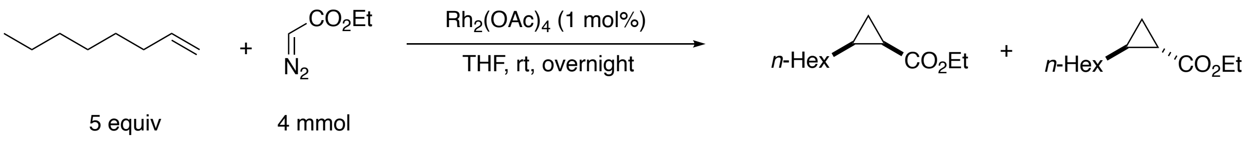


A solution of ethyl diazoacetate (1 equiv, 4 mmol, ≥13 wt.% DCM) in THF (1.2 ml) was added dropwise to a stirred mixture of 1-octene (5 equiv, 20 mmol) and rhodium (II) diacetate dimer (1.0 mol %, 0.04 mmol) in dry THF (2 ml) under an argon atmosphere. The reaction mixture was stirred for 16 h, then filtered through a short plug of alumina gel using diethyl ether as an eluent. The filtrate was concentrated *in vacuo*. The resulting residue was purified by column chromatography on silica gel in hexanes/EtOAc to afford the cyclopropanation products as a mixture of diastereomers.

ethyl 2-hexylcyclopropanecarboxylate ***C3-p***

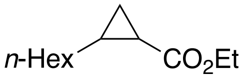


[1]H NMR (400 MHz, $CDCl_3$) δ 4.06 (q, *J* = 7.1 Hz, 2H), 1.59 (ddd, *J* = 8.9, 7.8, 5.5 Hz, 1H), 1.19 (td, *J* = 8.6, 4.2 Hz, 13H), 0.92 (td, *J* = 8.1, 4.4 Hz, 1H), 0.88 – 0.83 (m, 1H), 0.83 – 0.78 (m, 3H).

#### *Synthesis of* ***cis-C4-p*** *and* ***trans-C4-p***

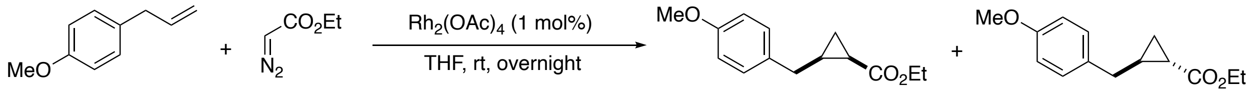


5 equiv 4 mmol

A solution of ethyl diazoacetate (1 equiv, 4 mmol, ≥13 wt.% DCM) in THF (1.2 ml) was added dropwise to a stirred mixture of 2-allylanisole (5 equiv, 20 mmol) and rhodium (II) diacetate dimer (1.0 mol %, 0.04 mmol) in dry THF (2 ml) under an argon atmosphere. The reaction mixture was stirred for 16 h, then filtered through a short plug of alumina gel using diethyl ether as an eluent. The filtrate was concentrated *in vacuo*. The resulting residue was purified by column chromatography on silica gel in hexanes/EtOAc to afford the cyclopropanation products as separate diastereomers.

(±)-*cis*-ethyl 2-(4-methoxybenzyl)cyclopropanecarboxylate ***cis-C4-p***

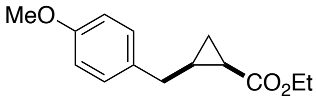


$^{1}$H NMR (400 MHz, $CDCl_3$) δ 7.17 – 7.08 (m, 2H), 6.88 – 6.78 (m, 2H), 4.11 (qd, $J$ = 7.2, 1.0 Hz, 2H), 3.79 (s, 3H), 2.70 (dd, $J$ = 14.8, 6.4 Hz, 1H), 2.52 (dd, $J$ = 14.8, 7.1 Hz, 1H), 1.71 – 1.61 (m, 1H), 1.49 (dt, $J$ = 8.5, 4.4 Hz, 1H), 1.31 – 1.16 (m, 4H), 0.81 (ddd, $J$ = 8.2, 6.4, 4.2 Hz, 1H).

(±)-*trans*-ethyl 2-(4-methoxybenzyl)cyclopropanecarboxylate ***trans-C4-p***

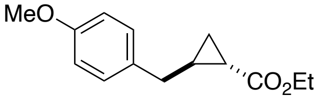


[1]H NMR (400 MHz, $CDCl_3$) δ 7.16 – 7.08 (m, 2H), 6.85 – 6.79 (m, 2H), 4.13 (q, $J$ = 7.1 Hz, 2H), 3.78 (s, 3H), 2.86 (dd, $J$ = 14.9, 6.9 Hz, 1H), 2.77 (dd, $J$ = 14.9, 7.6 Hz, 1H), 1.76 (ddd, $J$ = 8.8, 7.5, 5.8 Hz, 1H), 1.55 – 1.43 (m, 1H), 1.23 (d, $J$ = 7.2 Hz, 4H), 1.14 – 0.99 (m, 2H).

#### *Synthesis of* ***trans-C6-p***

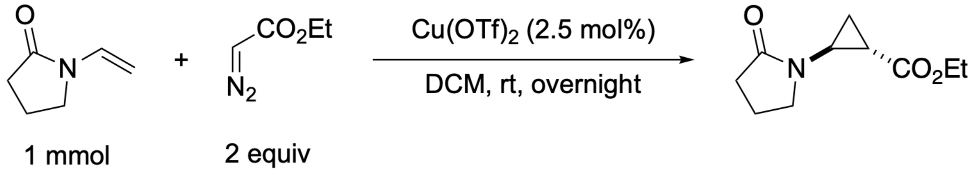


A solution of ethyl diazoacetate (2 equiv, 2 mmol, ≥13 wt.% DCM) in DCM (2 ml) was added dropwise to a stirred mixture of *N*-vinyl-pyrrolidone (1 mmol) and copper(II) trifluoromethanesulfonate (2.5 mol %, 0.025mmol) in dry DCM (5 ml) under an argon atmosphere. The reaction mixture was stirred for 16 h, then filtered through a short plug of silica gel using EtOAc as an eluent. The filtrate was concentrated *in vacuo*. The resulting residue was purified by column chromatography on silica gel in hexanes/EtOAc to afford the cyclopropanation product.

Ethyl 2-(2-oxopyrrolidin-1-yl)cyclopropane-1-carboxylate ***trans*-C6-p**

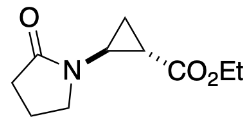


Yellow oil. $^{1}$H NMR (400 MHz, $CDCl_3$) δ 4.13 (qt, $J$ = 7.1, 1.2 Hz, 2H), 3.30 (dd, $J$ = 7.4, 6.6 Hz, 2H), 3.21 – 3.10 (m, 1H), 2.38 (t, $J$ = 8.1 Hz, 2H), 1.99 (dddd, $J$ = 14.9, 8.3, 7.3, 0.9 Hz, 2H), 1.83 (dddd, $J$ = 9.1, 5.9, 3.1, 0.7 Hz, 1H), 1.50 – 1.34 (m, 2H), 1.25 (td, $J$ = 7.1, 0.9 Hz, 3H). $^{13}$C NMR (101 MHz, $CDCl_3$) δ 176.2, 172.3, 61.0, 47.3, 34.1, 31.7, 19.8, 18.1, 14.2, 14.1.

#### *Synthesis of **C12-p***

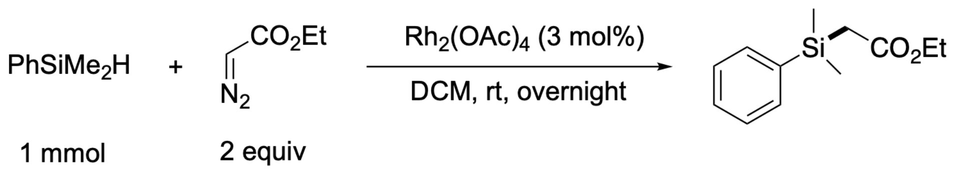


A solution of ethyl diazoacetate (2 equiv, 2 mmol, ≥13 wt.% DCM) in DCM (2 ml) was added dropwise to a stirred mixture of dimethylphenylsilane (1 mmol) and rhodium(II) acetate (3 mol%, 0.03 mmol) in dry DCM (5 ml) under an argon atmosphere. The reaction mixture was stirred for 16 h, then filtered through a short plug of silica gel using EtOAc as an eluent. The filtrate was concentrated *in vacuo*. The resulting residue was purified by column chromatography on silica gel in hexanes/EtOAc to afford the product.

Ethyl 2-(dimethyl(phenyl)silyl)acetate **C12-p**

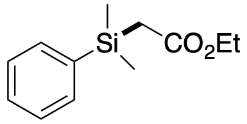


Colorless oil. $^{1}H$ NMR (400 MHz, $CDCl_3$) δ 7.59 – 7.48 (m, 2H), 7.44 – 7.32 (m, 3H), 4.04 (q, $J$ = 7.1 Hz, 2H), 2.11 (s, 2H), 1.16 (t, $J$ = 7.1 Hz, 3H), 0.41 (s, 6H). $^{13}C$ NMR (101 MHz, $CDCl_3$) δ 172.7, 137.1, 133.6, 129.6, 128.0, 60.1, 26.4, 14.4, -2.6.

#### *Synthesis of* ***C13-p***

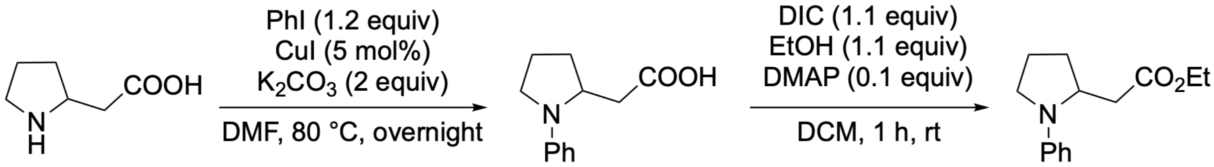


Step 1: To a sealed tube flushed with nitrogen were added pyrrolidine (5 mmol), potassium carbonate (2 equiv, 10 mmol), copper (I) iodide (0.25 mmol, 5 mol%), iodobenzene (1.2 equiv, 6 mmol) and DMF (7.5 ml). The mixture was heated at 90°C for 48 hours, then cooled to room temperature. Water was added, and the pH value was adjusted to <3 with concentrated HCl. The aqueous phase was extracted four times with ethyl acetate. The combined organic layers were washed with brine, dried over magnesium sulfate, filtered and concentrated *in vacuo*. Purification by silica gel chromatography (0 to 100% EtOAc/hexane gradient) afforded the product.

Step 2: A round-bottom flask was charged with carboxylic acid (1.0 equiv), ethanol (1.1 equiv), and DMAP (0.1 equiv). Dichloromethane was added (0.4 M), and the mixture was stirred vigorously. DIC (1.1 equiv) was then added dropwise via syringe, and the reaction mixture was stirred until consumption of the acid was complete, as determined by TLC. The mixture was filtered through a fritted funnel and rinsed with $CH_2Cl_2/Et_2O$. The solvent was removed under reduced pressure, and purification by column chromatography afforded the corresponding ester.

Ethyl 2-(1-phenylpyrrolidin-2-yl)acetate **C13-p**

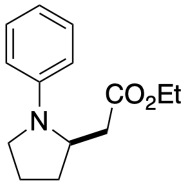


Yellow oil. $^1H$ NMR (400 MHz, $CDCl_3$) δ 7.26 – 7.22 (m, 2H), 6.71 – 6.65 (m, 1H), 6.65 – 6.55 (m, 2H), 4.20 – 4.15 (m, 3H), 3.46 – 3.39 (m, 1H), 3.22 – 3.13 (m, 1H), 2.79 (dd, *J* = 15.0, 2.9 Hz, 1H), 2.26 – 2.17 (m, 1H), 2.17 – 1.94 (m, 4H), 1.28 (t, *J* = 7.1 Hz, 3H). $^{13}C$ NMR (101 MHz, $CDCl_3$) δ 172.1, 146.5, 129.5, 129.4, 121.6, 116.0, 111.9, 55.4, 47.9, 37.8, 31.0, 23.1, 14.3.

*Synthesis of* ***C14-p***

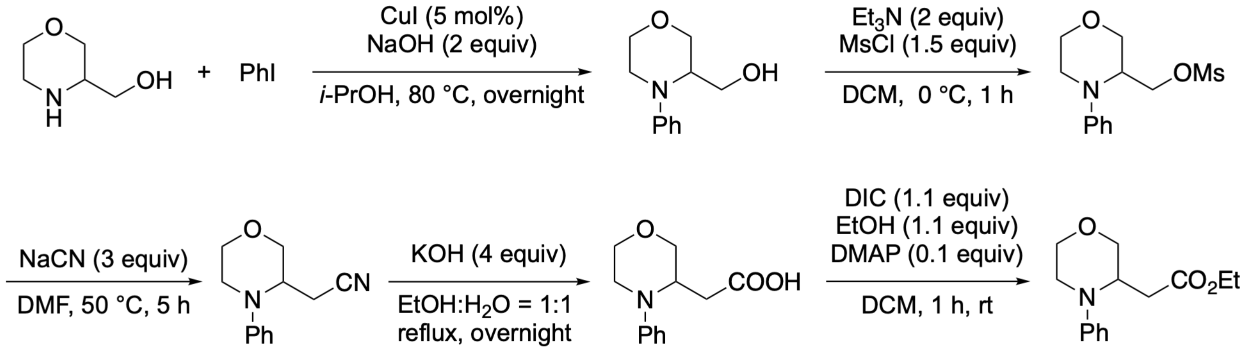


Step 1: A seal tube with a magnetic stirring bar was charged with CuI (1 mmol, 0.05 equiv.), powder NaOH (40 mmol, 2 equiv), and morpholin-3- ylmethanol (20 mmol, 1 equiv). To the tube were added isopropyl alcohol (5 mL, 0.8 M) and iodobenzene (24 mmol, 1.2 equiv) via syringe. The reaction was heated to 80 °C and left to stir overnight. The following day the reaction was reduced *in vacuo* and reconstituted in DCM and water. The mixture was introduced into a

separatory funnel and the organic layer was separated. The water layer was extracted twice with DCM, and the combined organic layers were washed with brine and dried over $MgSO_4$. The solution was reduced *in vacuo* and purified via flash chromatography, affording the arylation product as an oil.

Step 2: A round-bottom flask with a magnetic stirring bar was charged with (4- phenylmorpholin-3-yl)methanol (10 mmol, 1 equiv) and placed under an argon atmosphere. Thereafter, DCM (0.2 M) was added via syringe and the reaction was stirred and cooled to 0 °C (ice/water bath). Triethylamine (2 equiv, 2.0 mmol) and methanesulfonyl chloride (MsCl) (1.5 equiv, 1.5 mmol) were added dropwise in that order. The reaction was left to warm to room temperature and stirred for 1 hour at that temperature. The reaction was then introduced into a separatory funnel, washed with saturated $NaHCO_3$, brine, and dried over $MgSO_4$. The resultant oil was used immediately in the following reaction.

Step 3: A round-bottom flask with a magnetic stirring bar and the resultant oil from the previous reaction was charged with DMF (0.2 M) and NaCN (30 mmol, 3 equiv.). The reaction was sealed, blanketed with an argon atmosphere, and heated to 60 °C for 3 h. The reaction was cooled to room temperature and diluted with ethyl acetate and water. The organic layer was washed several times with brine, dried over $MgSO_4$, and reduced *in vacuo*. The resultant residue was purified using flash chromatography to afford the product as an oil.

Step 4: To a solution of 2-(4-phenylmorpholin-3-yl)acetonitrile (5 mmol) in ethanol (5 mL) and water (5 mL) was added potassium hydroxide (20 mmol) and the resulting mixture was refluxed overnight. The ethanol was removed under reduced pressure, and then the solution was cooled to below 10 °C. and acidified with concentrated HCl to pH 1. The mixture was extracted with EtOAc and the combined organic extracts were dried over anhydrous sodium sulfate and concentrated under vacuum to afford the carboxylic acid as a yellow oil.

Step 5: A round-bottom flask was charged with carboxylic acid (1.0 equiv, 3 mmol), ethanol (1.1 equiv), and DMAP (0.1 equiv). Dichloromethane was added (0.4 M), and the mixture was stirred vigorously. DIC (1.1 equiv) was then added dropwise via syringe, and the reaction mixture was stirred until consumption of the acid was complete, as determined by TLC. The mixture was filtered through a fritted funnel and rinsed with $CH_2Cl_2/Et_2O$. The solvent was removed under reduced pressure, and purification by column chromatography afforded the corresponding ester.

Ethyl 2-(4-phenylmorpholin-3-yl)acetate **C14-p**

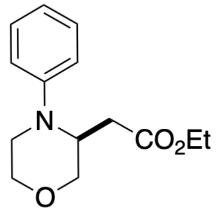


Yellow oil. $^1$H NMR (400 MHz, $CDCl_3$) δ 7.31 – 7.25 (m, 2H), 6.92 – 6.82 (m, 3H), 4.19 – 4.14 (m, 1H), 4.09 – 3.98 (m, 3H), 3.92 (dt, *J* = 11.6, 1.5 Hz, 1H), 3.85 (ddd, *J* = 11.5, 2.8, 1.6 Hz, 1H), 3.72 (td, *J* = 11.3, 3.4 Hz, 1H), 3.19 (ddd, *J* = 12.7, 3.4, 1.8 Hz, 1H), 3.10 (ddd, *J* = 12.2, 11.3, 3.7 Hz, 1H), 2.86 (dd, *J* = 15.8, 10.2 Hz, 1H), 2.36 (ddd, *J* = 15.8, 3.4, 1.6 Hz, 1H), 1.21 (t, *J* = 7.1 Hz, 3H). $^{13}$C NMR (101 MHz, $CDCl_3$) δ 172.6, 149.3, 129.8, 120.1, 115.9, 69.8, 67.3, 60.9, 52.5, 43.4, 30.5, 14.5.

*Synthesis of* ***C10-p***

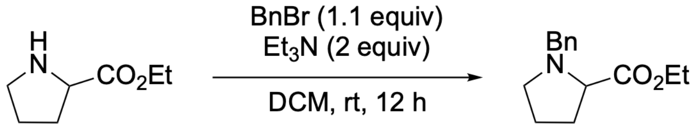


Triethylamine (2 equiv, 2 mmol) was added to a solution of pyrrolidine (1 mmol) in 10 mL DCM at room temperature. Benzyl bromide (1.1 equiv, 1.1 mmol) was added dropwise to the reaction mixture and the resulting solution was refluxed for 12 hrs. After cooling, sodium hydroxide (10 mL of 1 N solution) was added to the reaction mixture, and the product was extracted with DCM (2 x 30 mL). All the organic layers were combined, washed with brine, dried over $MgSO_4$, and concentrated under reduced pressure to afford the crude product. The resulting residue was purified by column chromatography on silica gel in hexanes/EtOAc to afford the product.

Ethyl benzyl-prolinate **C10-p**

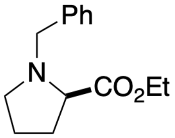


$^1$H NMR (400 MHz, $CDCl_3$) δ 7.31 – 7.21 (m, 4H), 7.21 – 7.14 (m, 1H), 4.06 (qd, *J* = 7.1, 1.9 Hz, 2H), 3.86 (d, *J* = 12.7 Hz, 1H), 3.49 (d, *J* = 12.7 Hz, 1H), 3.17 (dd, *J* = 8.8, 6.3 Hz, 1H), 2.98 (ddd, *J* = 9.0, 7.6, 3.1 Hz, 1H), 2.32 (td, *J* = 8.8, 7.8 Hz, 1H), 2.13 – 2.01 (m, 1H), 1.95 – 1.77 (m, 2H), 1.77 – 1.65 (m, 1H), 1.19 (t, *J* = 7.1 Hz, 3H). $^{13}$C NMR (101 MHz, $CDCl_3$) δ 174.3, 138.5, 129.3, 128.3, 127.2, 60.6, 58.8, 53.3, 29.4, 23.0, 14.4.

#### *Synthesis of* ***N9-p***

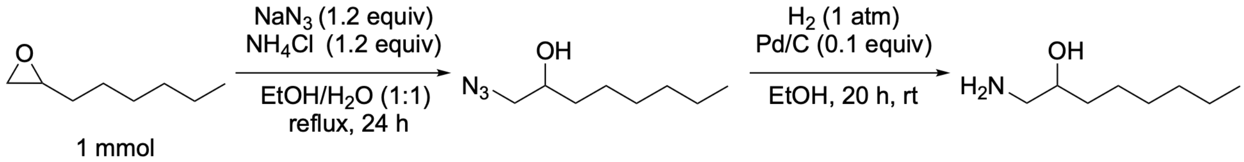


$NaN_3$ (1.2 equiv, 1.2 mmol), $NH_4Cl$ (1.2 equiv, 1.2 mmol) and a 1,2-epoxy-*n*-alkane (1 mmol) were dissolved in sufficient EtOH/$H_2O$ (1:1 v:v) to give a homogeneous solution. The solution was refluxed for 24 h, then the EtOH was removed *in vacuo.* The residual aq. soln. was extracted with $Et_2O$ (3 × 10 ml) and the combined organic extracts washed with water and dried ($MgSO_4$). After filtration and concentration *in vacuo*, purification by flash column chromatography (hexanes/EtOAc) gave a colorless oil. The azidoalcohol and Pd/C catalyst (0.1 equiv, w:w) were placed in a flask with EtOH (3 ml). The mixture was degassed by three freeze-pump-thaw cycles from an $H_2$ atmosphere. The mixture was then stirred under $H_2$ (balloon, 1 atm.) at room temperature for 20 h. After filtration through Celite®, the EtOH was removed *in vacuo* to give the product as a colorless oil. No further purification was necessary.

1-Aminooctan-2-ol **N8-p**

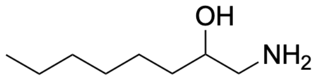


Colorless oil. $^1$H NMR (400 MHz, $CDCl_3$) δ 3.81 – 3.71 (m, 1H), 3.38 (dd, *J* = 12.4, 3.3 Hz, 1H), 3.25 (dd, *J* = 12.4, 7.4 Hz, 1H), 2.05 (s, 1H), 1.46 (ddt, *J* = 15.8, 11.7, 5.1 Hz, 3H), 1.36 – 1.23 (m, 7H), 0.92 – 0.85 (m, 3H). $^{13}$C NMR (101 MHz, $CDCl_3$) δ 71.0, 57.2, 34.4, 31.8, 29.3, 25.5, 22.7, 14.2.

### Preparative-Scale Enzymatic Reactions

**Protocols for Preparative-Scale Enzymatic Reactions:**

*Liter-Scale Protein Expression*

Single colonies from LB-Agar plates were picked using a sterile pipette tip and were used to inoculate 50 mL of LB-Amp in a 125-mL Erlenmeyer flask. Cultures were incubated at 37 ºC with shaking at 220 rpm overnight in an Innova 4000 shaking incubator. A 25-mL aliquot of each of these overnight cultures was used to inoculate 1 L of Terrific Broth with 100 μg/mL of ampicillin (TB-Amp) (0.5% v/v starter culture in expression culture) in a 3.2-L unbaffled Erlenmeyer flask. The expression cultures were incubated at 37 ºC and 220 rpm for 2.5 hours in an Innova 42 shaking incubator, at which point they were moved to room temperature for 25 minutes. Protein expression was then induced by direct addition of 50 μL of stock solutions containing 500 mM isopropyl-β-D-thiogalactoside (IPTG) and 1.0 M 5-aminolevulinic acid (ALA) such that the final concentrations were 0.5 mM and 1.0 mM, respectively. The cultures were shaken at 22 ºC and 140 rpm for 16–18 hours in an Innova 42 shaker.

*Small-Scale Biocatalytic Carbene Transfer Reaction Preparation*

The corresponding l-L expression cultures were pelleted (4,000 × g for 30 minutes at 4 ºC) and resuspended in 100 mL of M9-N buffer. The optical density at 600 nm ($OD_{600}$) of this suspension was measured and adjusted to $OD_{600} = 31.5$ with the addition of more M9-N buffer. A 38-mL aliquot of the cell suspension was added to a 60-mL test tube with screw-cap seal. These whole-cell suspensions were transferred into a vinyl Coy anaerobic chamber, at which point 1 mL of a solution of the reaction substrate in MeCN was added. Subsequently, 1 mL of a solution of EDA in MeCN was added dropwise to the reaction vessel to prevent rapid release of $N_2$ gas upon diazo activation. The GC vials were tightly capped with screw caps equipped with septum and were allowed to shake at room temperature for 16 hours.

Once complete, the reaction was split into two 20-mL aliquots across two 50-mL falcon tubes. The suspensions are extracted three times with ethyl acetate as follows: 20 mL of ethyl acetate were added to each tube and the phases were mixed by hand-shaking the tubes. Upon thorough mixing, the phases were separated by centrifugation (5000 × g, 5 minutes, RT) and the organic phase was siphoned off. The combined organics were dried over sodium sulfate; once dry, the drying agent was decanted, and the volatiles removed under vacuum. The crude residue was purified by silica gel column chromatography to yield the titled compounds.

*Characterization of isolated products*

Ethyl (*E*)-2-(2-((*tert*-butoxycarbonyl)amino)ethylidene)cyclopropane-1-carboxylate ***E*-C8-p**

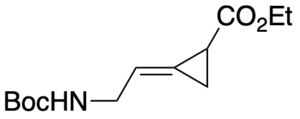


Colorless oil. $^{1}$H NMR (400 MHz, $CDCl_3$) δ 5.91 (ddt, $J$ = 5.7, 3.7, 2.1 Hz, 1H), 4.67 (s, 1H), 4.16 (qd, $J$ = 7.2, 1.2 Hz, 2H), 3.85 (d, $J$ = 6.2 Hz, 2H), 2.29 (dddd, $J$ = 6.9, 4.6, 2.2, 1.1 Hz, 1H), 1.73 (td, $J$ = 5.5, 2.3 Hz, 1H), 1.61 (tt, $J$ = 6.6, 2.2 Hz, 1H), 1.44 (s, 9H), 1.27 (t, $J$ = 7.2 Hz, 3H). $^{13}$C NMR (101 MHz, $CDCl_3$) δ 172.2, 155.9, 123.5, 116.4, 79.5, 61.0, 41.6, 36.7, 28.5, 24.8, 23.4, 17.4, 14.3, 10.6. HRMS (FI+) *m/z*: $[M]^{\cdot+}$ calcd. for $C_{13}H_{21}NO_4$ 255.1465, found: 255.1463.

Ethyl (*Z*)-2-(2-((*tert*-butoxycarbonyl)amino)ethylidene)cyclopropane-1-carboxylate ***Z*-C8-p**

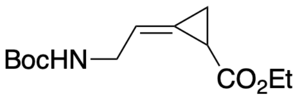


Colorless oil. $^1$H NMR (400 MHz, $CDCl_3$) δ 5.91 (ddq, $J$ = 5.7, 4.1, 2.2 Hz, 1H), 4.75 (s, 1H), 4.13 (qd, $J$ = 7.1, 0.8 Hz, 2H), 3.91 (d, $J$ = 7.3 Hz, 2H), 2.26 – 2.16 (m, 1H), 1.81 (ddd, $J$ = 8.9, 4.5, 2.4 Hz, 1H), 1.63 (ddd, $J$ = 8.6, 7.5, 2.2 Hz, 1H), 1.44 (s, 9H), 1.26 (d, $J$ = 7.1 Hz, 3H). $^{13}$C NMR (101 MHz, $CDCl_3$) δ 172.2, 123.4, 116.9, 79.5, 60.9, 41.5, 28.5, 16.9, 14.3, 10.9. HRMS (FI+) *m/z*: $[M]^{\cdot+}$ calcd. for $C_{13}H_{21}NO_4$ 255.1465, found: 255.1468.

Ethyl (*E*)-2-(cyclohexylmethylene)cyclopropane-1-carboxylate ***E*-C9-p**

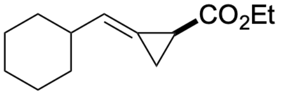


Colorless oil. $^1$H NMR (400 MHz, $CDCl_3$) δ 5.87 – 5.69 (m, 1H), 4.12 (q, $J$ = 7.1 Hz, 2H), 2.26 – 2.11 (m, 2H), 1.83 – 1.60 (m, 7H), 1.34 – 1.15 (m, 8H). $^{13}$C NMR (101 MHz, $CDCl_3$) δ 172.7, 125.3, 119.2, 60.3, 39.8, 32.4, 32.2, 26.0, 25.8, 25.8, 16.4, 14.0, 11.4.

Diethyl 1-phenylbicyclo[1.1.0]butane-2,4-dicarboxylate **C7-p**

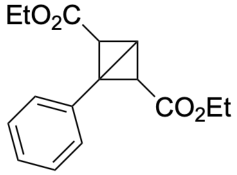


Colorless oil. $^1$H NMR (400 MHz, $CDCl_3$) δ 7.56 – 7.49 (m, 2H), 7.33 – 7.26 (m, 3H), 4.08 – 3.93 (m, 4H), 3.40 (s, 1H), 1.81 (s, 2H), 1.57 (s, 2H), 1.08 (t, $J$ = 7.1 Hz, 6H). $^{13}$C NMR (101 MHz, $CDCl_3$) δ 167.3, 131.7, 129.3, 128.1, 127.9, 60.8, 40.5, 25.6, 15.1, 14.1.

### NMR Spectra of Authentic Standards and Isolated Products

[1]H NMR (400 MHz, $CDCl_3$) of ***trans*-C6-p**

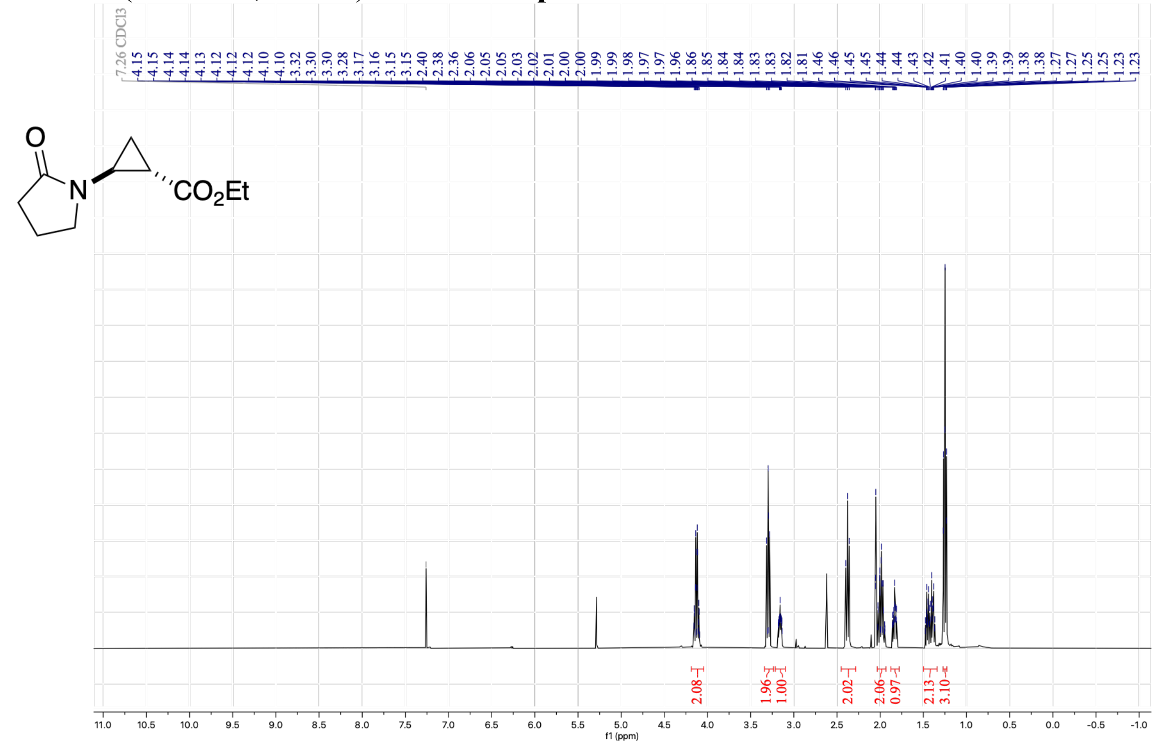


[13]C NMR (101 MHz, $CDCl_3$) of ***trans*-C6-p**

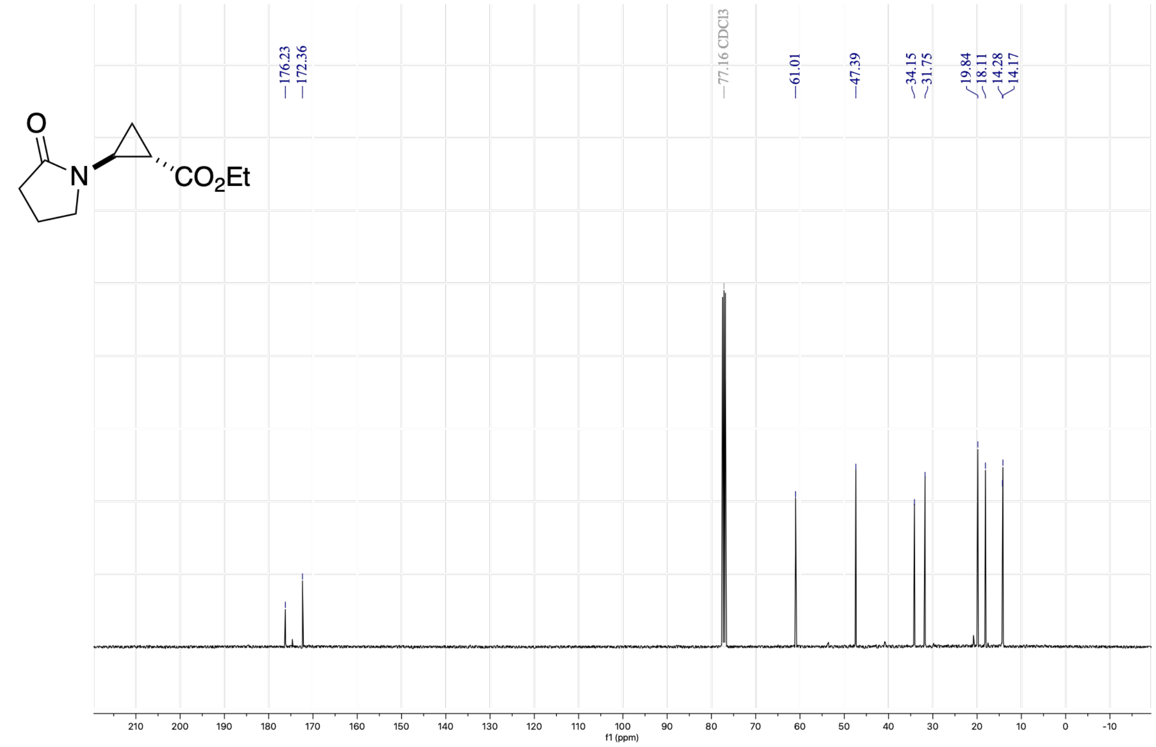

$^{1}H$ NMR (400 MHz, $CDCl_3$) of **C12-p**

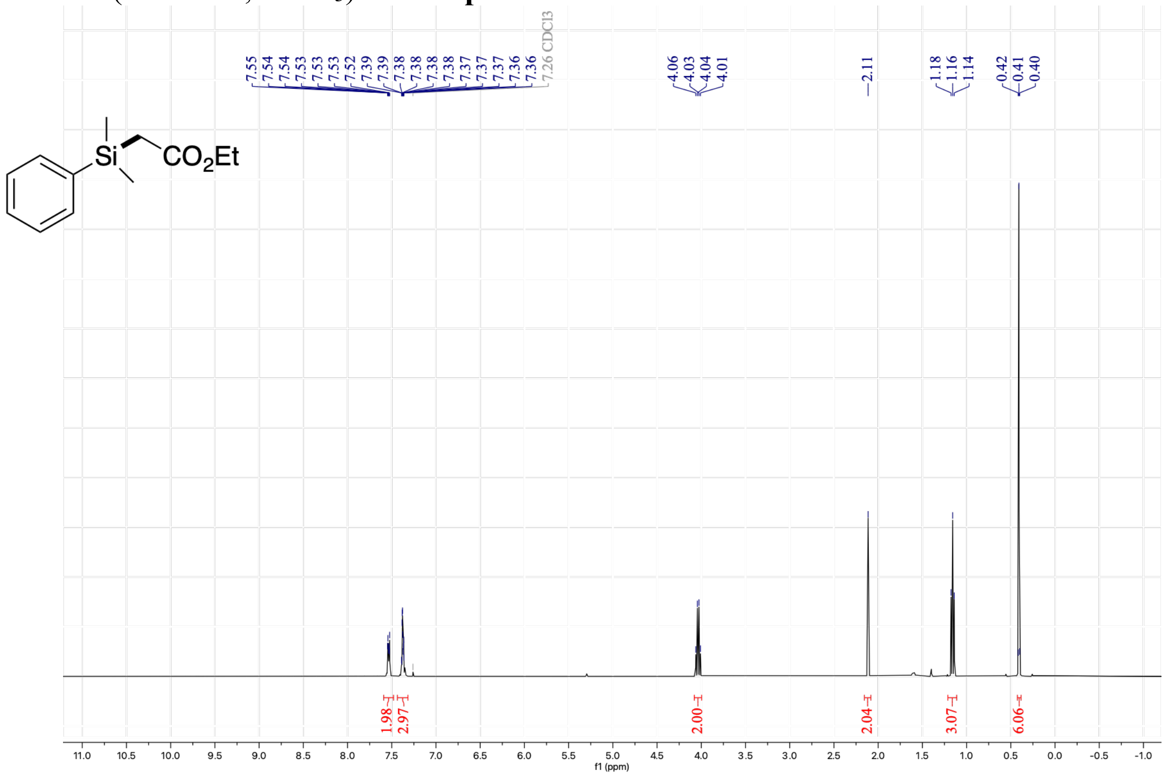


$^{13}C$ NMR (101 MHz, $CDCl_3$) of **C12-p**

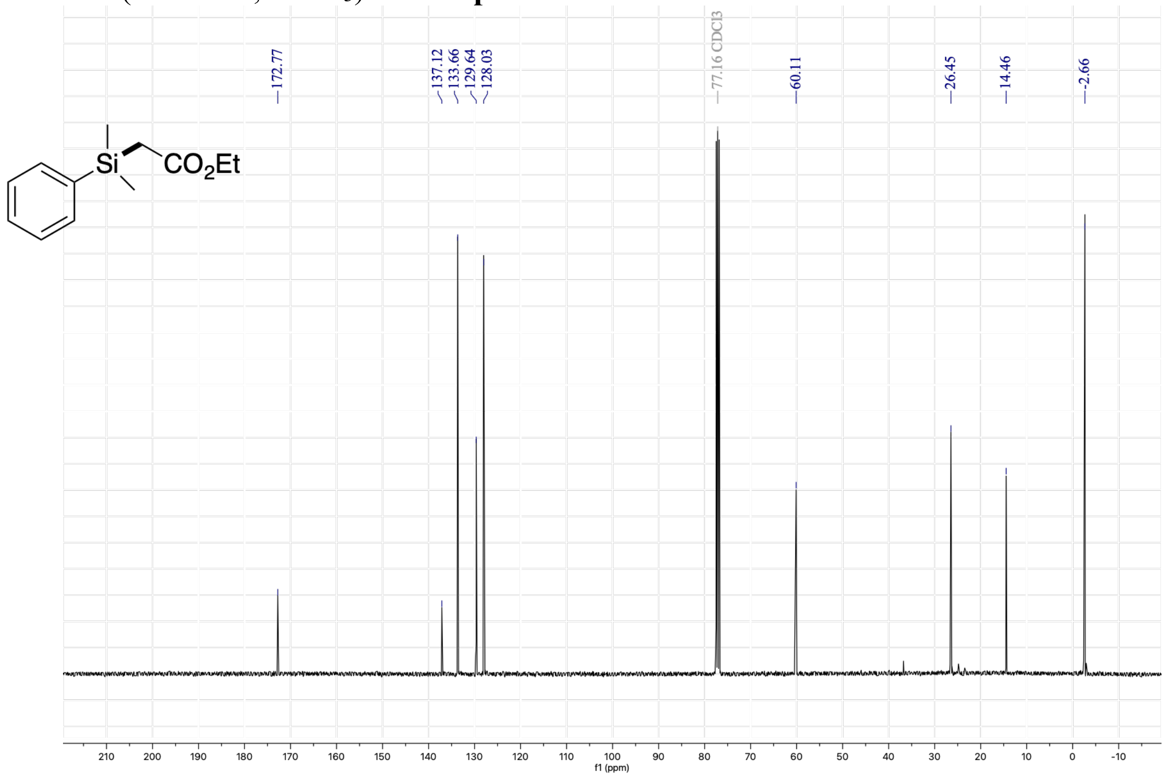

[1]H NMR (400 MHz, $CDCl_3$) of **C13-p**

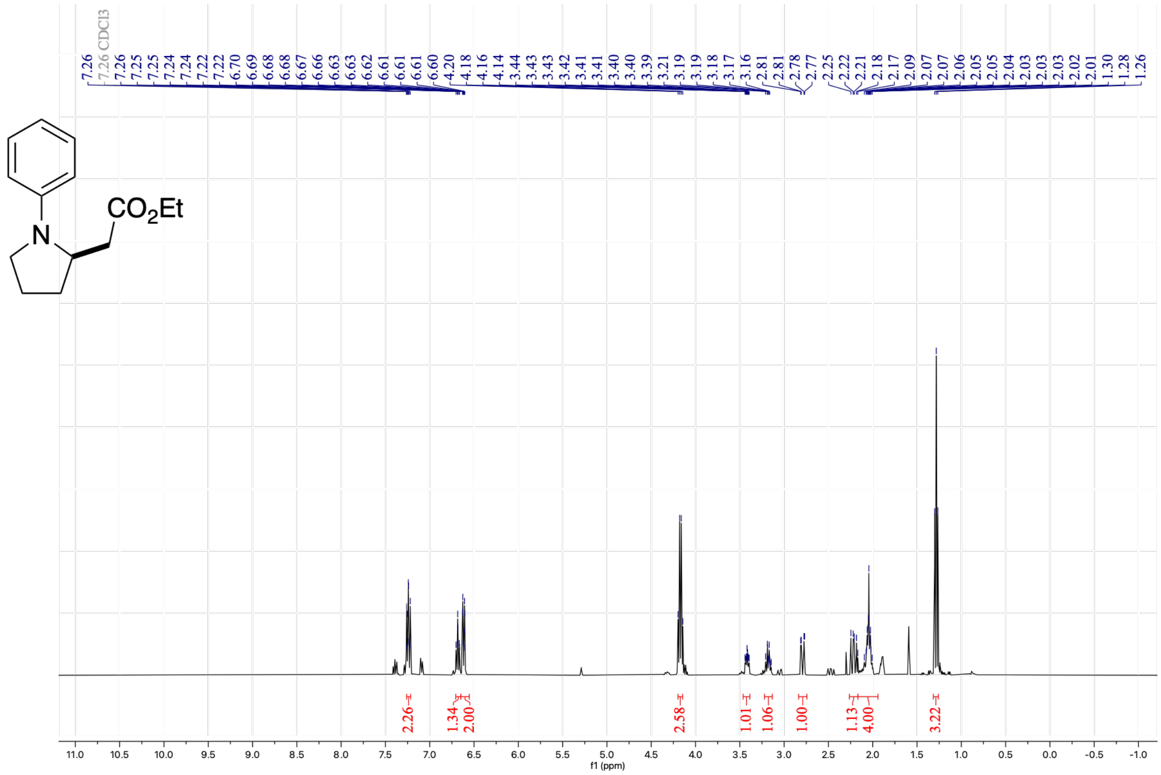


[13]C NMR (101 MHz, $CDCl_3$) of **C13-p**

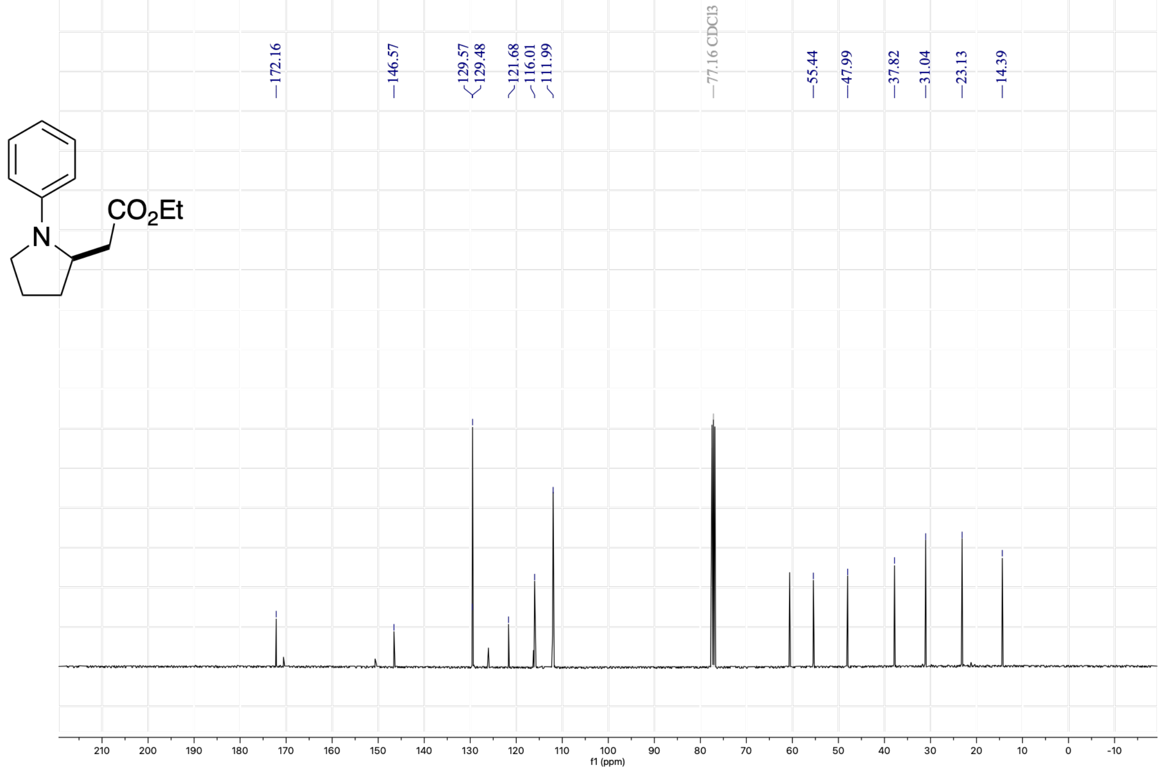

$^{1}$H NMR (400 MHz, $CDCl_3$) of **C10-p**

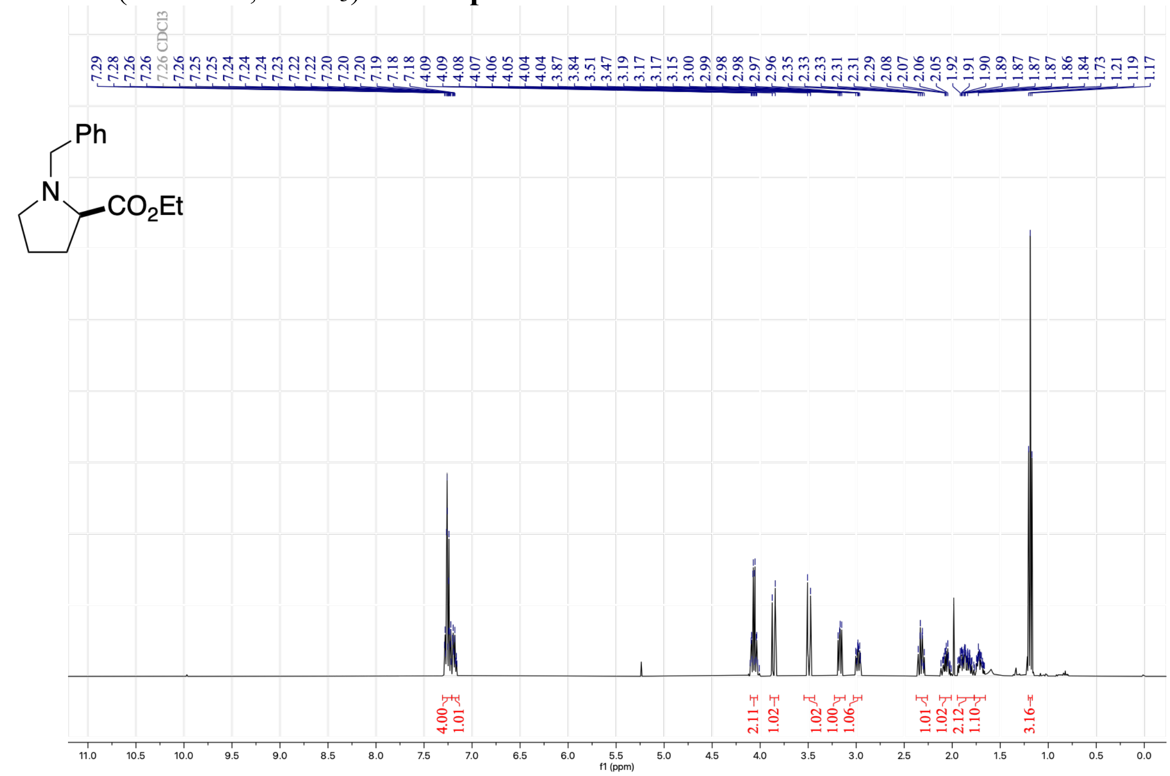


$^{13}$C NMR (101 MHz, $CDCl_3$) of **C10-p**

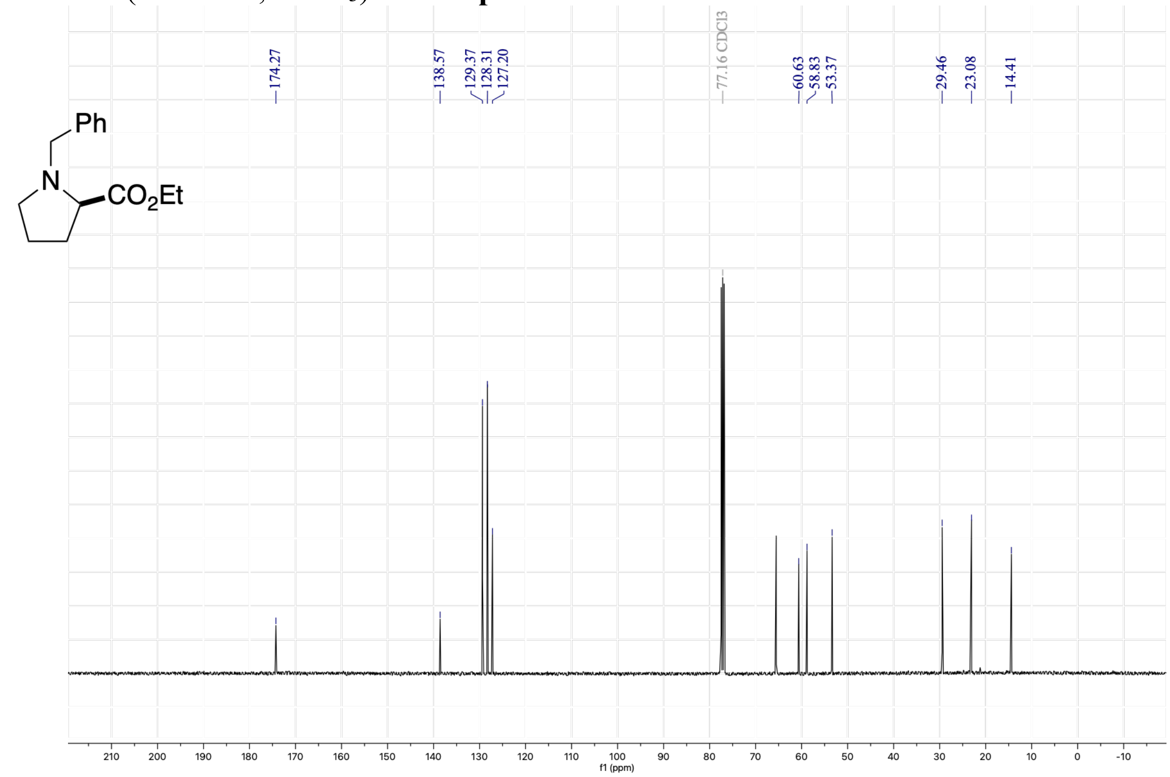

[1]H NMR (400 MHz, $CDCl_3$) of **C14-p**

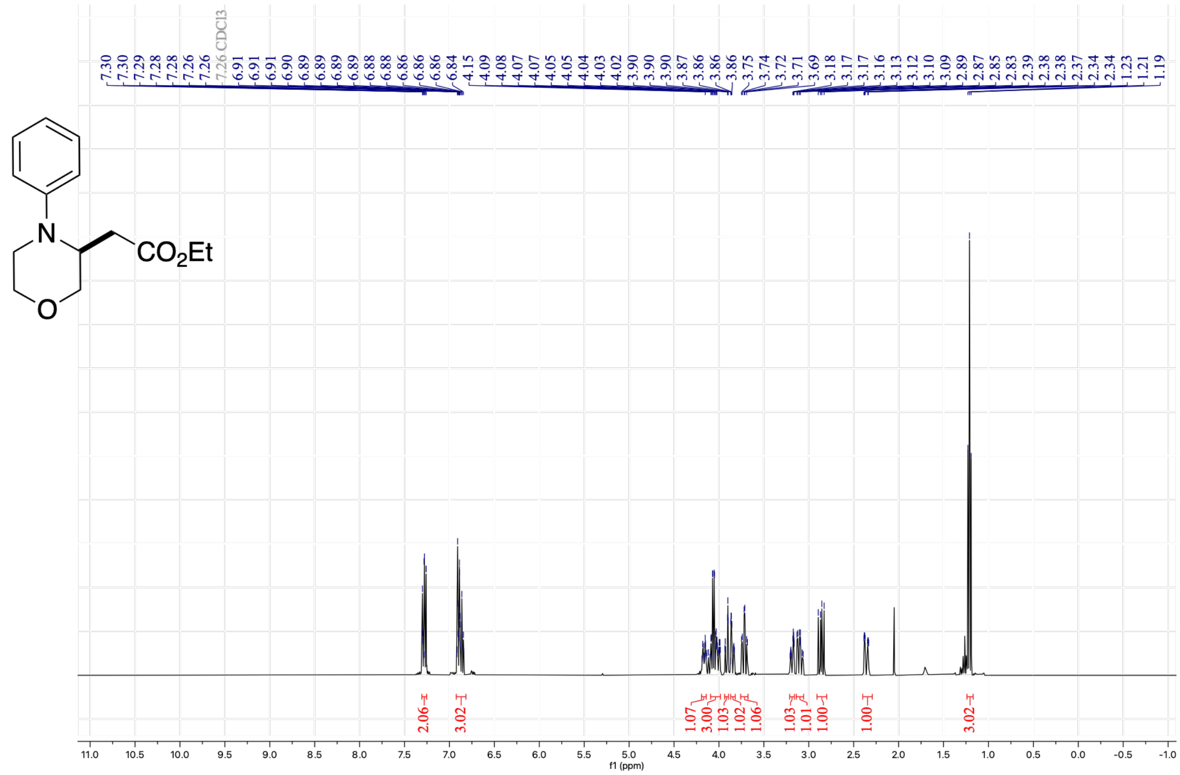


[13]C NMR (101 MHz, $CDCl_3$) of **C14-p**

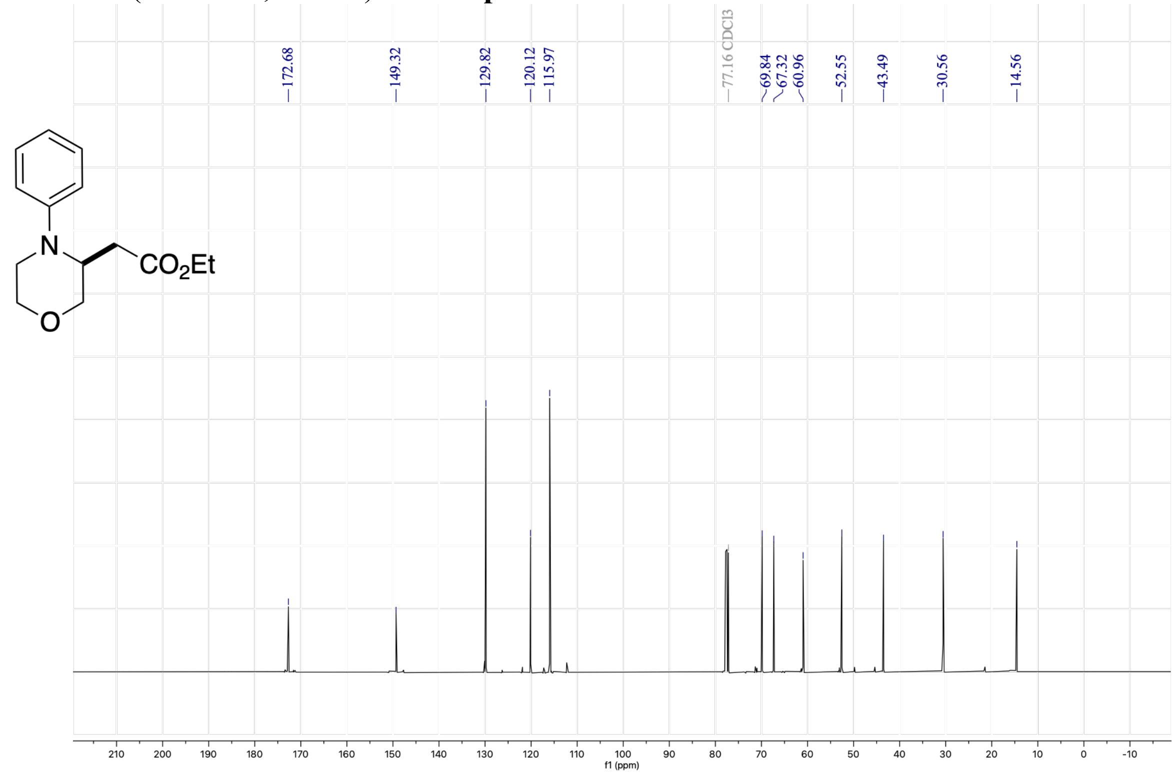

[1]H NMR (400 MHz, $CDCl_3$) of **N9-p**

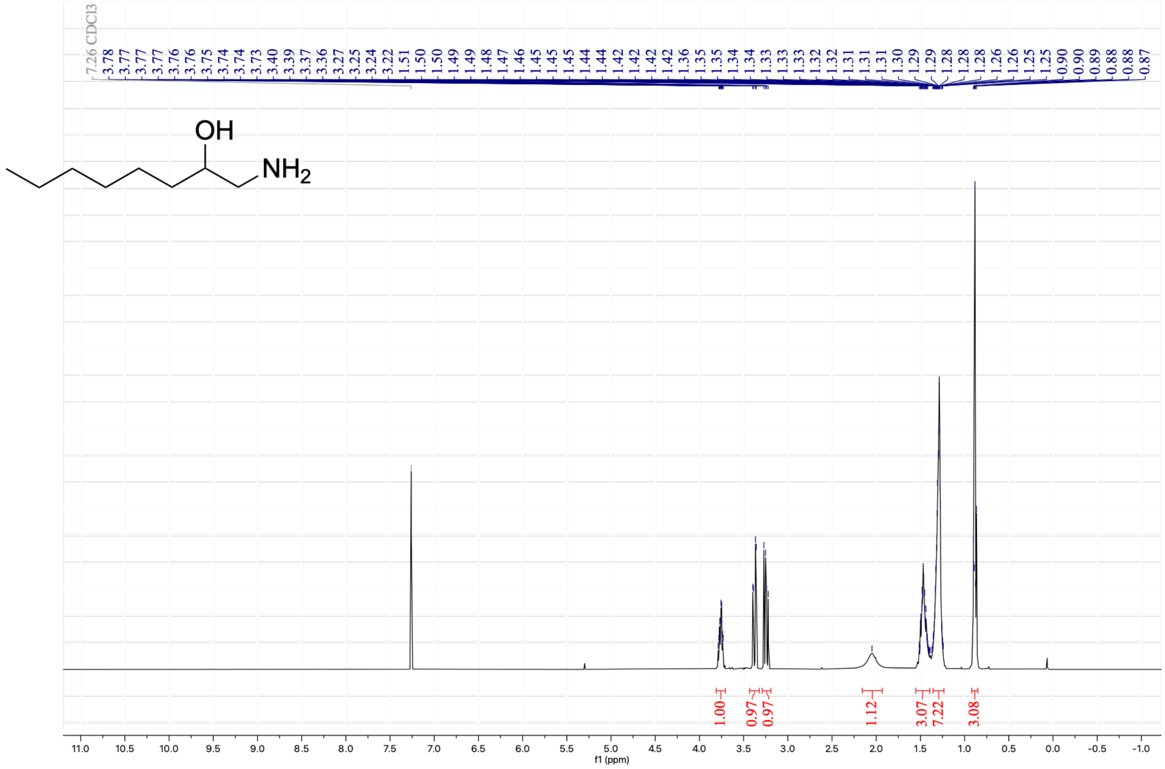


[13]C NMR (101 MHz, $CDCl_3$) of **N9-p**

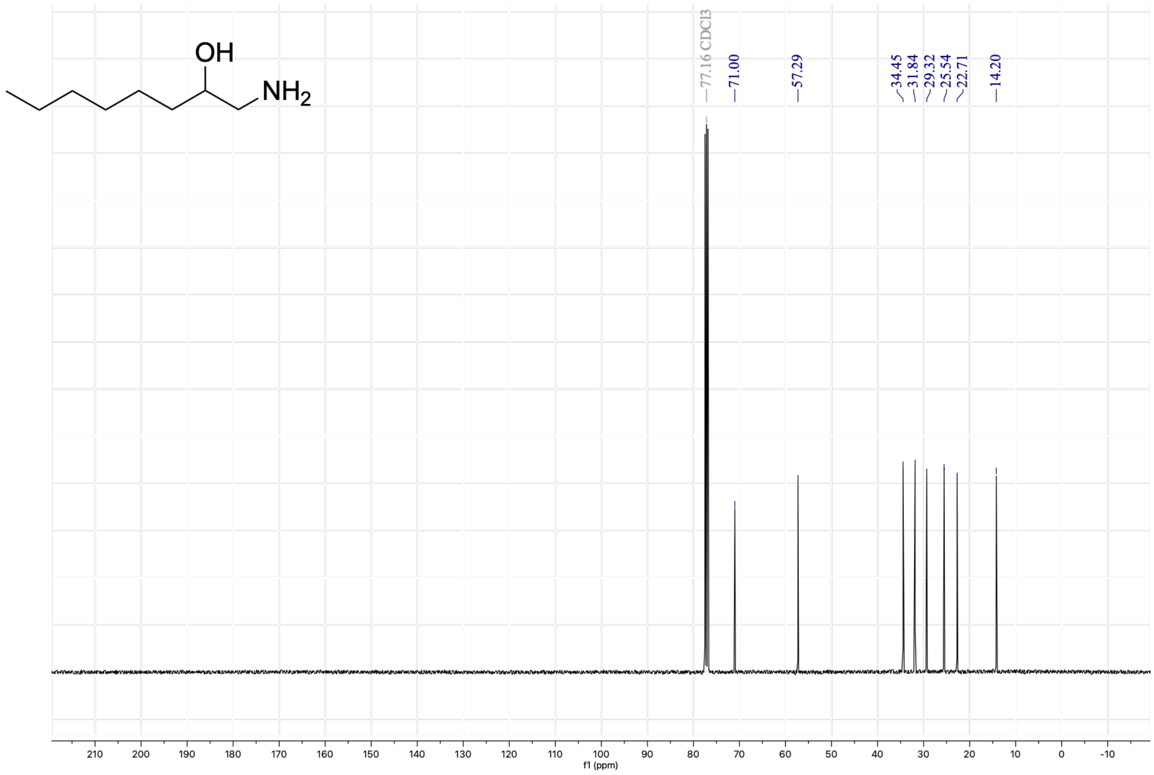

[1]H NMR (400 MHz, $CDCl_3$) of ***E*-C8-p**

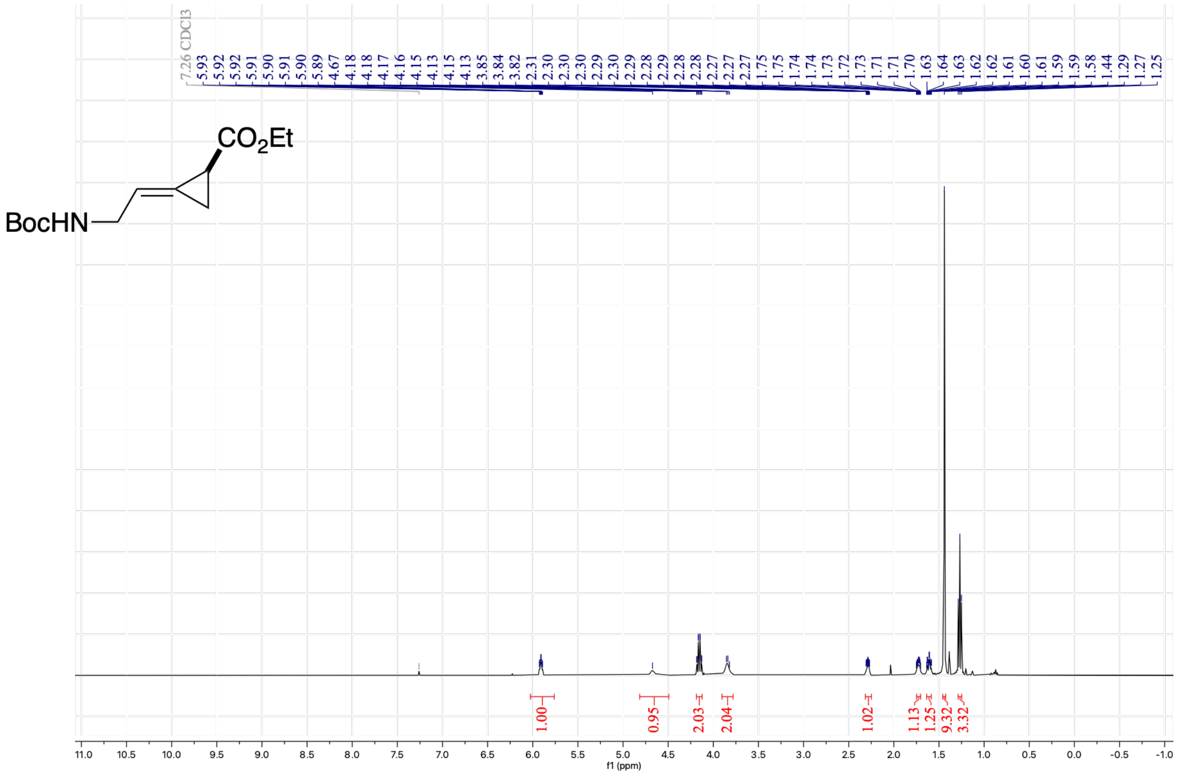

[13]C NMR (101 MHz, $CDCl_3$) of ***E*-C8-p**

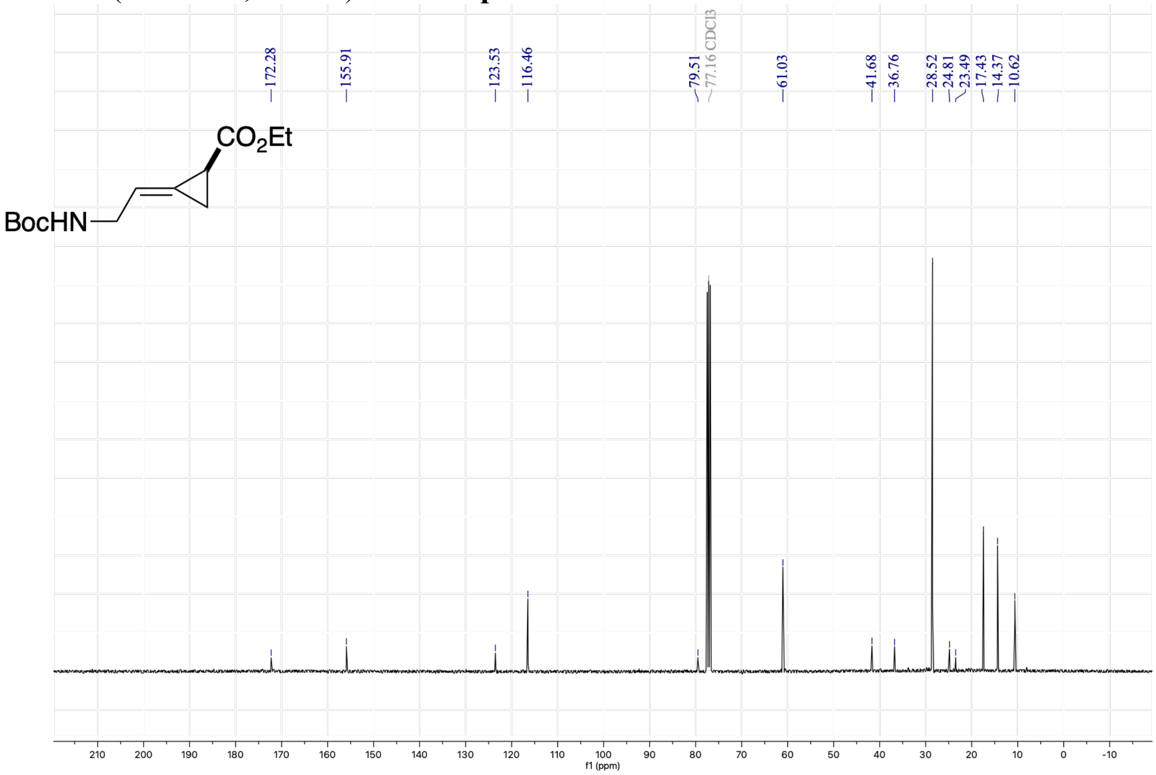

[1]H NMR (400 MHz, $CDCl_3$) of ***Z*-C8-p**

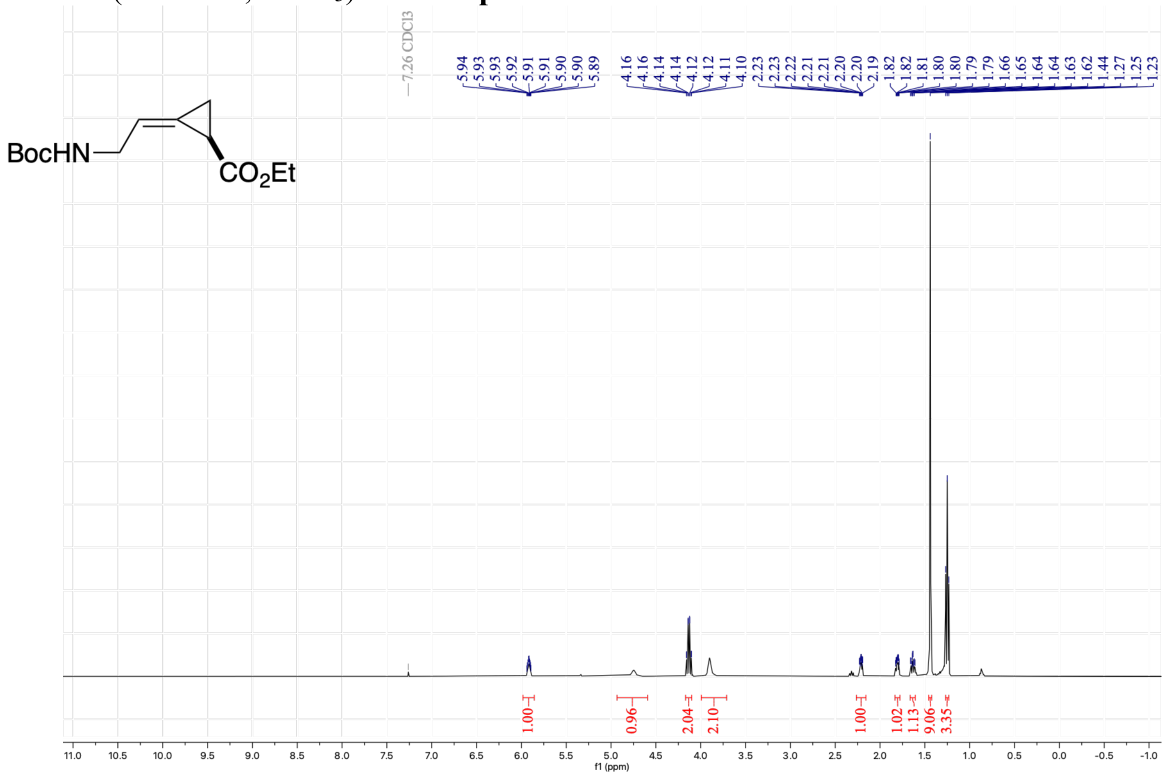


[13]C NMR (101 MHz, $CDCl_3$) of ***Z*-C8-p**

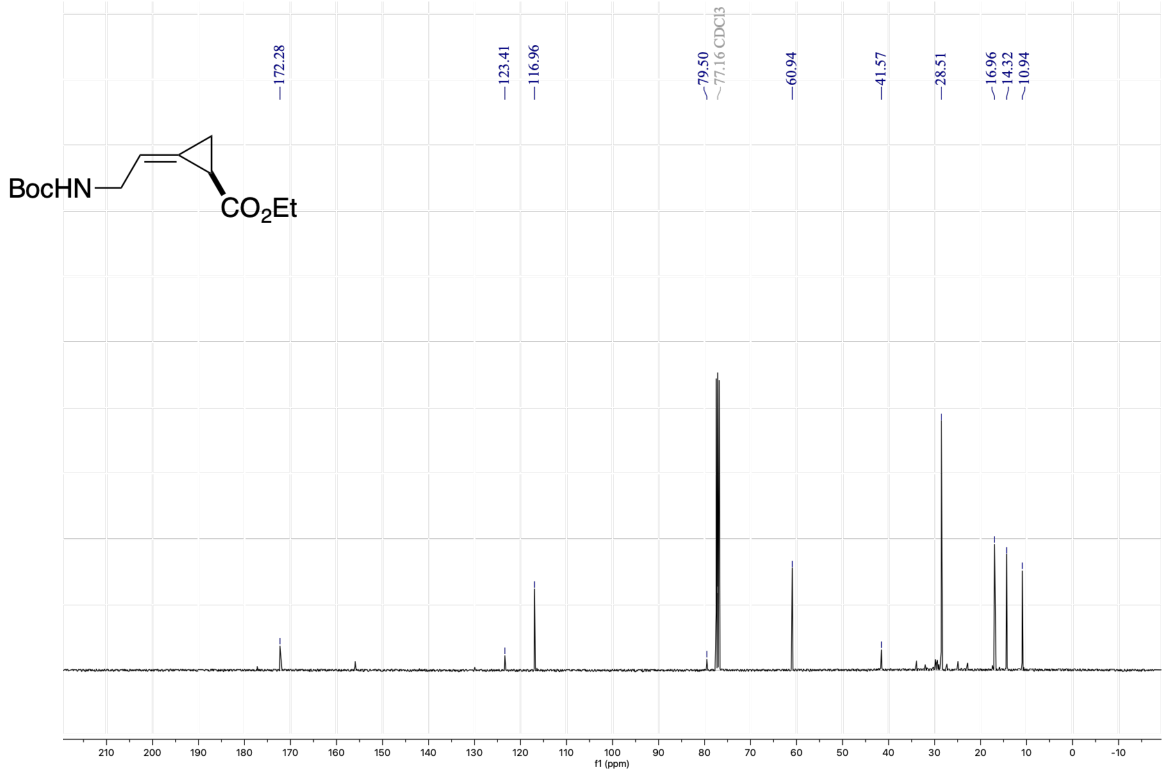

[1]H NMR (400 MHz, $CDCl_3$) of ***E*-C9-p**

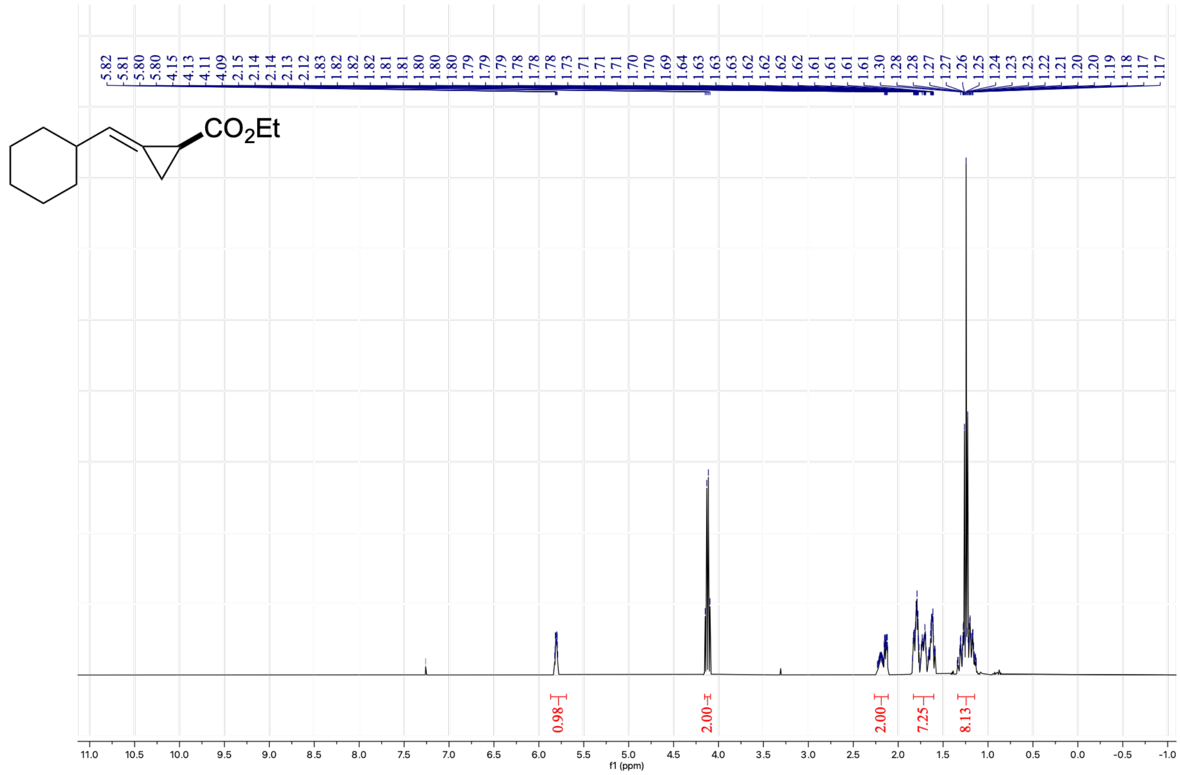


[13]C NMR (101 MHz, $CDCl_3$) of ***E*-C9-p**

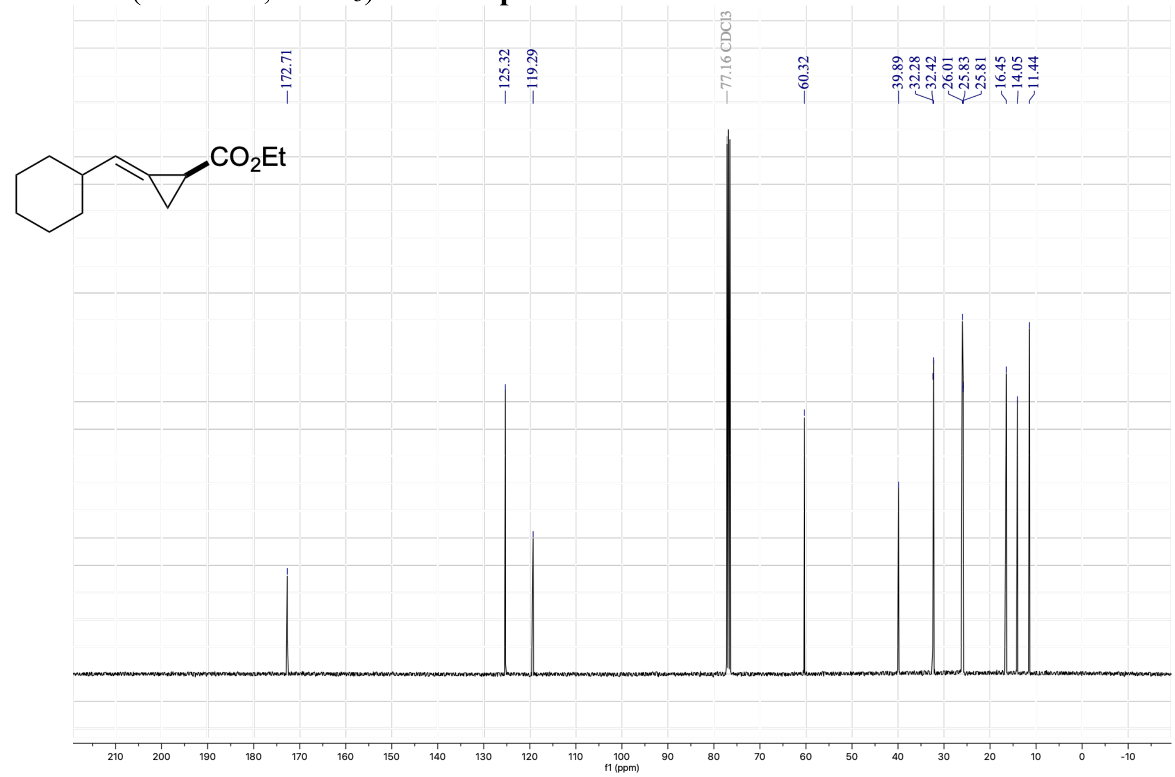

[1]H NMR (400 MHz, $CDCl_3$) of **C7-p**

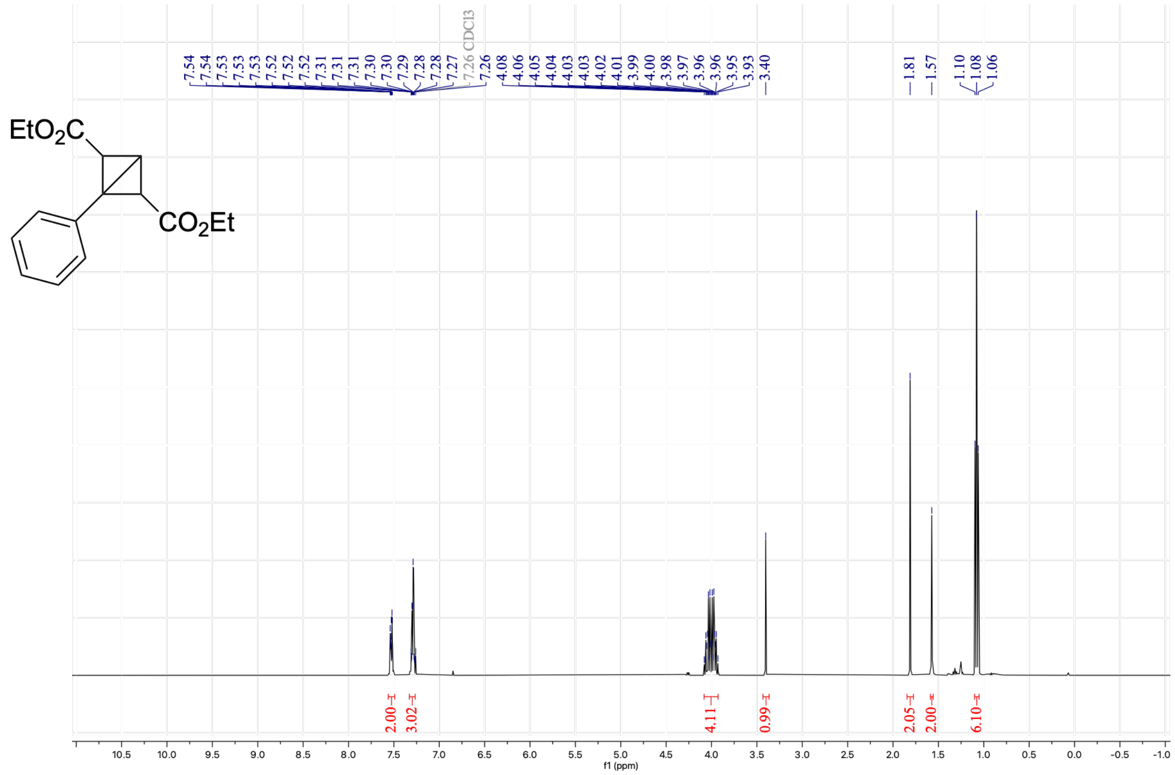


[13]C NMR (101 MHz, $CDCl_3$) of **C7-p**

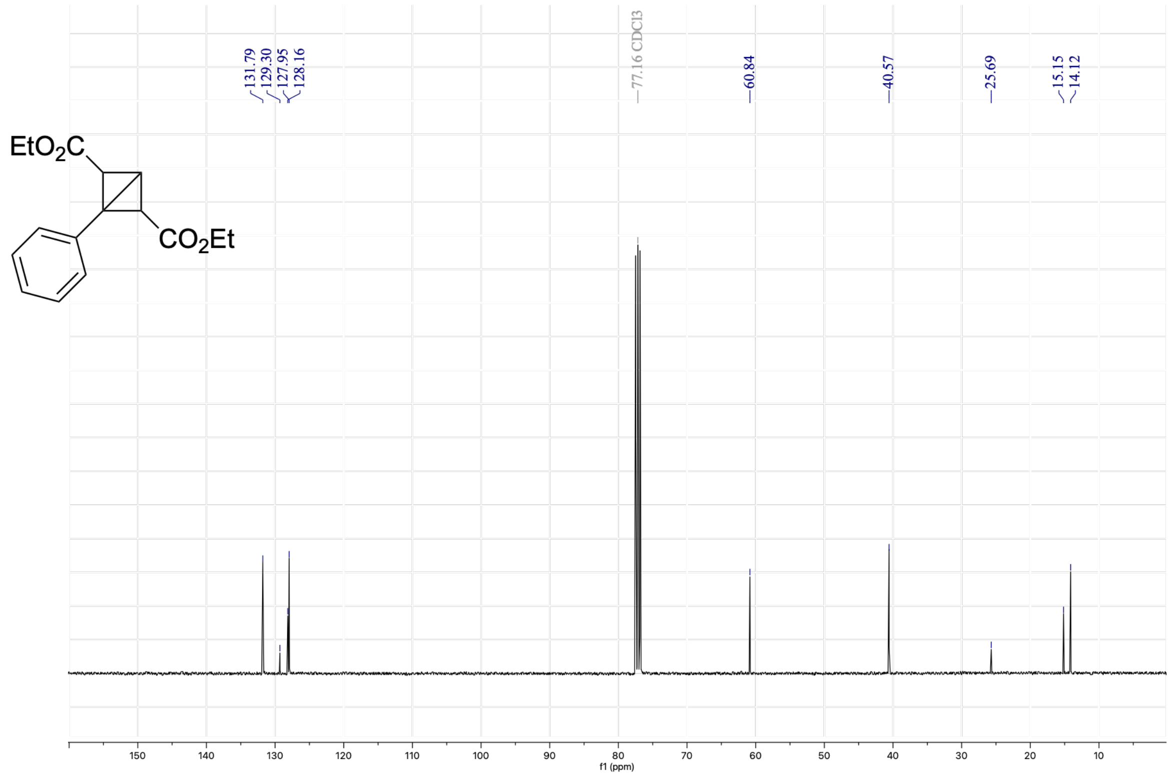
